# Parametrically guided design of beta barrels and transmembrane nanopores using deep learning

**DOI:** 10.1101/2024.07.22.604663

**Authors:** David E. Kim, Joseph L. Watson, David Juergens, Sagardip Majumder, Ria Sonigra, Stacey R. Gerben, Alex Kang, Asim K. Bera, Xinting Li, David Baker

**Affiliations:** Department of Biochemistry, University of Washington, Seattle, WA 98195; Institute for Protein Design, University of Washington, Seattle, WA 98195; HHMI, University of Washington, Seattle, WA 98195

**Author notes:** Contributed equally to this work (David E. Kim, Joseph L. Watson, David Juergens, Sagardip Majumder).

**Keywords:** protein design, machine learning, beta barrels, nanopores

## Abstract

Francis Crick’s global parameterization of coiled coil geometry has been widely useful for guiding design of new protein structures and functions. However, design guided by similar global parameterization of beta barrel structures has been less successful, likely due to the deviations from ideal barrel geometry required to maintain inter-strand hydrogen bonding without introducing backbone strain. Instead, beta barrels have been designed using 2D structural blueprints; while this approach has successfully generated new fluorescent proteins, transmembrane nanopores, and other structures, it requires expert knowledge and provides only indirect control over the global shape. Here we show that the simplicity and control over shape and structure provided by parametric representations can be generalized beyond coiled coils by taking advantage of the rich sequence-structure relationships implicit in RoseTTAFold based design methods. Starting from parametrically generated barrel backbones, both RFjoint inpainting and RFdiffusion readily incorporate backbone irregularities necessary for proper folding with minimal deviation from the idealized barrel geometries. We show that for beta barrels across a broad range of beta sheet parameterizations, these methods achieve high in silico and experimental success rates, with atomic accuracy confirmed by an X-ray crystal structure of a novel barrel topology, and de novo designed 12, 14, and 16 stranded transmembrane nanopores with conductances ranging from 200 to 500 pS. By combining the simplicity and control of parametric generation with the high success rates of deep learning based protein design methods, our approach makes the design of proteins where global shape confers function, such as beta barrel nanopores, more precisely specifiable and accessible.

**Significance:** De novo beta barrel proteins have previously been designed using “blueprint” based methods which require expert knowledge of the rules of folding and provide only indirect control of the overall barrel shape by specifying structural features such as glycine kinks and beta bulges. The barrel shape can be directly modeled using global parametric methods, but to date such methods have not succeeded in generating folded proteins, likely due to the absence of such structural features. Here, we describe methods that combine the simplicity and control of parametric barrel specification with the high success rates of deep learning based protein design methods to successfully design new beta barrel folds of different and pre-specified sizes, including both soluble designs and transmembrane nanopores.

## Introduction

Generation of protein backbones using the Crick parametrization of coiled coils^1^ followed by loop building and amino acid sequence design has been used to design a wide variety of structures^2,3^ and functions^4,5^. However, a similar approach based on parametrically generated beta barrel backbone structures did not yield designs which folded when produced in the lab, even when the backbones were relaxed to improve hydrogen bonding using Rosetta based structural relaxation^6^. Instead, beta barrels with new structures and functions ranging from fluorescent sensors^6^ to transmembrane channels^9,10^ have been designed using Rosetta “blueprints.” While powerful, this approach can control the overall shape only indirectly through explicit specification of blueprint features such as the strand number, strand lengths, barrel shear^6–8^, backbone hydrogen bonding arrangement, positioning of irregularities such as glycine kinks and beta bulges required for continuous hydrogen bonding and backbone strain reduction^6,11^, and tyrosine corners^6^. This complexity of blueprint design limits exploration of beta barrel structure space even with considerable human expertise.

In parallel, deep-learning methods have revolutionized de novo protein design, with methods such as Hallucination^12–16^, RFjoint inpainting^13^, and more recently generative models such as RFdiffusion^17^ demonstrating improved performance across a range of protein design tasks as compared to classical energy-based methods such as Rosetta^18^. These neural network based methods are capable of implicitly parameterizing the “rules” of protein folding, and are thus able to incorporate necessary irregularities into designed beta barrels that fold accurately^7^. To date, however, precise parameter based control over global fold geometry has not been achievable with deep-learning methods. There is therefore an unmet need to marry the benefits of classical design (precise control of structure parameters) with the benefits of deep-learning (simplicity of use, higher computational and experimental success rates, and higher design complexity).

We reasoned that the rich understanding of sequence structure relationships implicit in deep learning based design approaches and their ability to be guided by partial structural information could enable beta barrel design based on global parameterization of barrel size and shape by first generating backbones using parametric design, and then refining them using deep learning approaches. We investigated both inpainting and diffusion approaches for designing beta barrels from global parameterizations of beta barrel shape.

### Guiding RFjoint2 inpainting with global fold parameterizations

We first explored the ability of RFjoint2, an improved version of RoseTTAFold based RFjoint inpainting (see Supplementary text and Figure S1), to design beta barrels with imperfect templates as input structures. RFjoint2 was trained to address the motif-scaffolding problem, where the network is provided with a protein substructure (the “motif”), and is tasked with generating the rest of the protein (the “scaffold”; we refer to this process as “inpainting” but in other fields it is referred to as outpainting). RFjoint2 is able to generate large scaffolds around a small input motif described using 2D inter-residue pairwise distances, and pairwise dihedral angles. The underlying RoseTTAFold model can take as input homologous structures (“templates”), accompanied by a one-dimensional “confidence” feature, a measure of how similar the provided template is to the query sequence. This notion of an “imperfect” template input is maintained in RFjoint2; thus, RFjoint2 can refine imperfect input structures provided to the network along with confidence values.

For input templates, we systematically generated Cɑ atom cylinders^6,19^ with defined parameter ranges in the number of strands (n), the shear number (S; the register shift between the first and last strands), and the strand length (l) (see Methods). In the context of a protein beta barrel structure, the radius and core packing can be described by n and S, and together with l, the height of the cylinder can be specified^20,21^. These Cɑ positions approximate the beta barrel fold to be designed, but are not optimal since variations in beta strand lengths and imperfections such as glycine kinks and beta bulges are typically necessary for proper barrel closure, backbone hydrogen bonding, and hydrophobic core packing^6,11^. We hypothesized that RFjoint2 would be able to introduce the necessary features and imperfections to permit proper folding while closely maintaining the desired parameters given a cylinder with imperfect confidence as the input. Loops connecting the strands and terminal extensions were not provided in the input template, but were fully inpainted by RFjoint2. Because the template input to RoseTTAFold, as well as the initialized 3D frames, require the N and C backbone atoms, these were added to the Ca cylinders using BBQ (Backbone Building from Quadrilaterals)^22^. Although these positions are not physically plausible, we again reasoned they should be refinable by RFjoint2. No sequence input was provided to RFjoint2, leveraging the fact that RFjoint2 can condition on structure inputs in the absence of sequence.

### Guiding RFdiffusion with global fold parameterizations

RFdiffusion is a probabilistic diffusion model with excellent performance across a broad range of protein design tasks^16,17^. RFdiffusion outperforms RFjoint2 at motif scaffolding, and has been used in a wider range of protein design applications. We therefore developed a second, RFdiffusion-based method for designing pre-specified beta barrels. RFdiffusion is able to generate specific folds through conditioning on secondary structure and “block-adjacency” information^17,23^, but this is insufficient to specify all barrel parameters (for example the barrel shear; Figure S2). In RFdiffusion, protein backbone frames (N-Cɑ-C atoms, represented by a translational and rotational component) are randomly sampled from a reference distribution (3D Gaussian distribution for translations, uniform distribution on SO3 for rotations) at time *t*=*T*, and are then iteratively “de-noised” over *T* timesteps to generate novel protein backbones (at *t*=0). During training, protein structures from the Protein Data Bank^24^ (PDB) are sampled, and such structures are “noised” to a random degree (either a small amount, through the addition of a small number of noising steps, or more extensively, through the addition of a large number of noising steps). RFdiffusion is then trained to predict the de-noised structure. While, after training, RFdiffusion is generally used to design completely novel protein backbones^17^, Vázquez Torres et al introduced the concept of “partial diffusion” with RFdiffusion, whereby a specific protein backbone is “resampled” through the addition of just a few forward noising steps^16^. The fewer noising steps added to a given backbone, the closer the resulting structure is to the original input protein^16^, thus giving a means of tuning the divergence of the resulting structure from the input protein.

We sought to use partial diffusion to design parametrically-specified beta barrels. We again treat the design problem as a refinement of parametrically generated backbone cylinders. The non-physically realizable beta barrel cylinders were noised with a variable number of noising steps, and RFdiffusion was then used to “de-noise” these to generate viable barrels with parameters similar to those input to the network. During training, RFdiffusion learns the expected deviation from the “true” (*t*=0) structure to expect at each timestep t, and we found it optimal to place each residue in the input structure (pre-noising) in a close-to-realistic position, so that the partially diffused *t*=*t* input is compatible with a beta barrel at *t*=0. We hence first roughly modeled loops and terminal extensions to the cylinder inputs with PyRosetta^25^.

### Beta barrel design using RFjoint2 and RFdiffusion

The two approaches are summarized in Figure 1a. Provided with an input cylinder specifying beta barrel parameters (which without refinement are not designable; Figure S3), both methods were capable of refining these inputs into beta barrels passing stringent in silico metrics (Figure 1b-d) previously found to correlate with experimental success. As desired, the barrels very closely match the specified parameters (Figure 1b,c).

**Figure 1:**
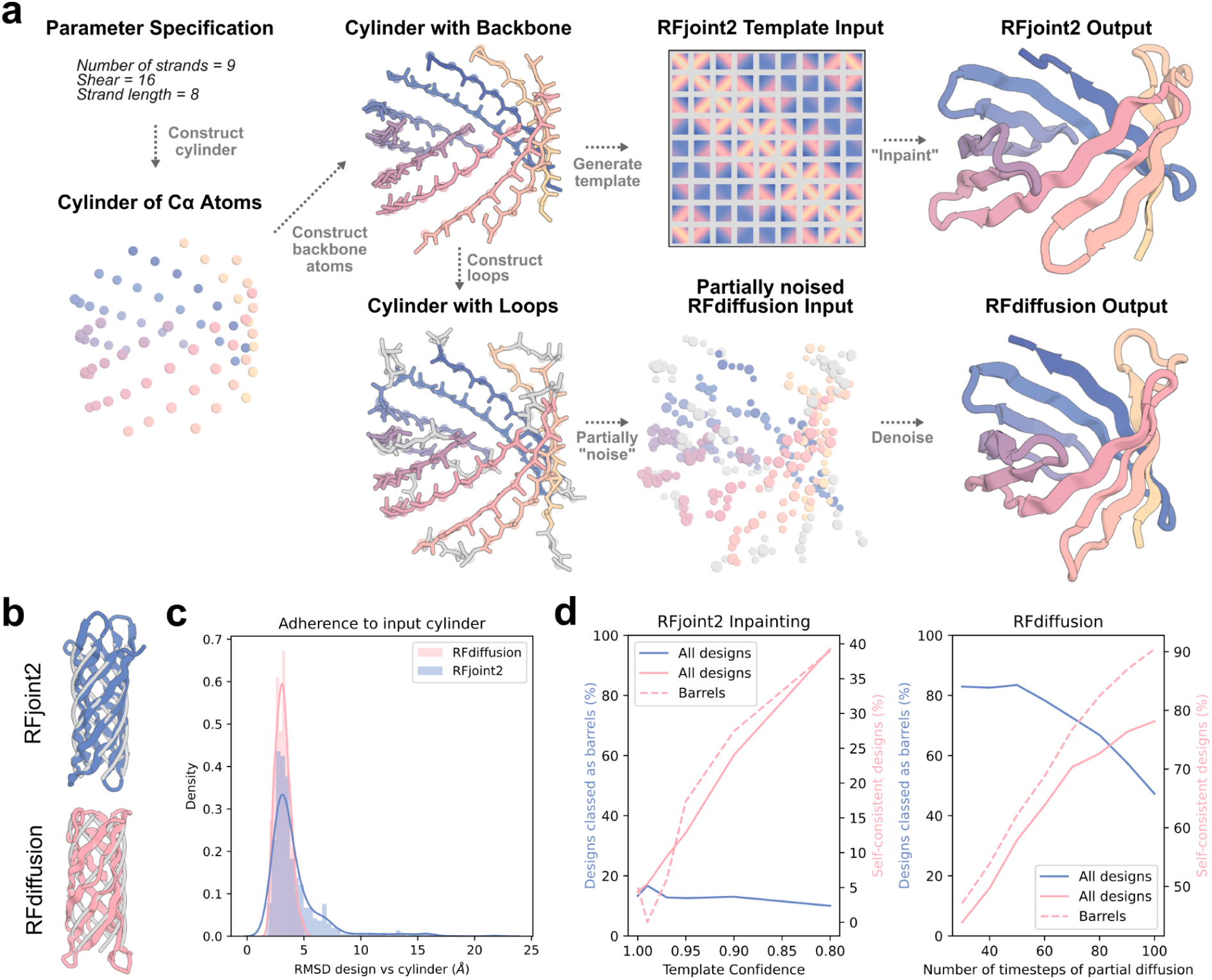
Generation of parametrically-defined beta barrels with RFjoint2 and RFdiffusion. Given a specification of global fold parameters, Cα-atoms are placed and then connected into protein backbones. In the RFjoint2 approach, (top row), these are used as low confidence inputs for protein inpainting (Methods) which outputs complete structures closely matching the input parametric structures with de novo connecting loops. In the RFdiffusion approach (bottom row), parametric backbones are completed with loops and partially noised followed by de-noising to generate more realistic protein backbones that closely match the input parameters. **b**) Representative parameter-matched barrels (colors) overlaid on the input cylinder (gray) for RFjoint2 (top) and RFdiffusion (bottom). **c**) Of the designs that are classified as barrels, the adherence to the input cylinder (i.e. the specified barrel parameters) is similar between RFjoint2 and RFdiffusion. The examples in **b** represent the “median” similarity of barrel to input cylinder (< 0.2 Å from respective medians in **c**). **d**) Dependence of design output on input parameters. Left: a high (> 0.5) template confidence was required in RFjoint2 for outputs to resemble barrels (blue line). However, self-consistency progressively decreases with increasing template confidence (solid pink line). The optimal template confidence in 0.8, where a good fraction of outputs resemble barrels, and self-consistency is still tolerable. Right: the more steps of partial diffusion (i.e., the more noise added to the input cylinder), the lower the rate of beta barrels observed (blue line). However, AF2 self-consistency increases (pink lines). 50 steps of partial diffusion were used for subsequence analyses. Success is defined as AF2 pLDDT > 80, RMSD between design and AF2 prediction < 2 Å, and both the FastRelaxed^18^ design and FastRelaxed AF2 prediction having hydrogen-bonding patterns consistent with barrel formation (Methods).

Both approaches have means of tuning the design process. In RFdiffusion, the number of steps of noise added to the input cylinder is expected to control the deviation of the output structure from the original input cylinder, and indeed, while at low (30-50) steps of added noise, over 80% of outputs were beta barrels, at higher step numbers (80-100) this fell sharply (Figure 1d right, blue curve). On the other hand, consistent with the failure of global parametric methods to generate designable structures, the fewer the steps of noise added and hence the more closely tied the design is to the input cylinder, the lower the in silico success rate (Figure 1d right, pink curves). There is thus a tradeoff between constraining the design by the desired cylinder parameters and generating high quality design outputs. For all further analyses in the paper, 50 steps of noise addition was used as a compromise between adherence to the input cylinder and in silico success rate. In RFjoint2, the design process can be tuned by varying the template confidence associated with the input cylinder. A higher template confidence more strongly constrains the design, with the output more closely matching the input cylindrical parameters. Indeed, the proportion of outputs classed as beta barrels (i.e., constrained by the input cylinder) increased approximately linearly with the template confidence value. (Figure 1d left, blue curve), but overall a far smaller fraction of designs were classed as barrels as compared to RFdiffusion. In a similar manner to the RFdiffusion approach, the more constrained (barrel-like) the outputs, the lower the in silico success rate (Figure 1d left, pink curves). The optimal template confidence was approximately 0.8, where the proportion of outputs classed as barrels was relatively high, while maintaining reasonable in silico success.

Although both methods were able to generate in silico successful beta barrels across nearly the whole spectrum of parameters tested, the RFdiffusion-based approach outperformed RFjoint2, in many cases generating successful designs with > 50% in silico success rate (Figure S4a-c). The two approaches have similar compute requirements, with each design taking around 1 minute on an NVIDIA A4000 GPU. Beta barrels generated using both methods closely adhered to the input cylinder (Figure 1c), highlighting that, although empirical success rates differ, both methods achieve the goal of allowing design of parameter-specified beta barrels. Inspection of the output structures demonstrated that both deep-learning methods have implicitly learned the requirement for backbone abnormalities (glycine kinks, beta bulges) and variable strand lengths in barrel folding; rules that were explicitly incorporated with previous blueprint energy-based approaches.

While the RFdiffusion-based method outperformed the RFjoint2-based method, the latter has its own advantages. First, with autoregressive sequence design in RFjoint2 (see Supplementary text), the network is able to generate high quality sequences competitive with ProteinMPNN^26^ while simultaneously designing protein structure. Second, while for the partial diffusion approach in RFdiffusion, every residue in the input barrels must be in the neighborhood of a physically plausible position such that the partially-diffused inputs could have emanated from a beta barrel, RFjoint2 only requires partial template specification so input loops and termini are not required. This additional flexibility could be advantageous in design scenarios where a portion of a protein is specified parametrically and the rest is free to fully reconfigure around this substructure.

### Experimental characterization of parametrically-defined beta barrels

Because there has been quite extensive experimental validation of RFdiffusion^16,17^, we focused on designs generated with RFjoint2. Designs with confident in silico metrics (see Methods) were selected for in vitro biophysical characterization. Synthetic genes encoding the designed sequences were ordered and the proteins were expressed in *Escherichia coli* and purified using immobilized metal affinity chromatography (IMAC). Out of 96 designs, 72 expressed, were soluble, and mostly monomeric as suggested by size exclusion chromatography (SEC; Figure S4). 32 designs expressed in sufficient quantities in a 96 well format for far-UV circular dichroism (CD); 29 of the 32 had CD spectra suggesting folded structures with minima in the range indicating beta structure, and 14 appeared to be stable at 95°C (Figure S5). An additional design of a small four stranded barrel had a structure and CD spectrum typical of a Src homolog 3 (SH3) beta-barrel fold^7^ with a maxima at around 230nm (Figure 2a). Experimental results from soluble monomeric designs made with varying parameters are shown in Figure 2.

**Figure 2:**
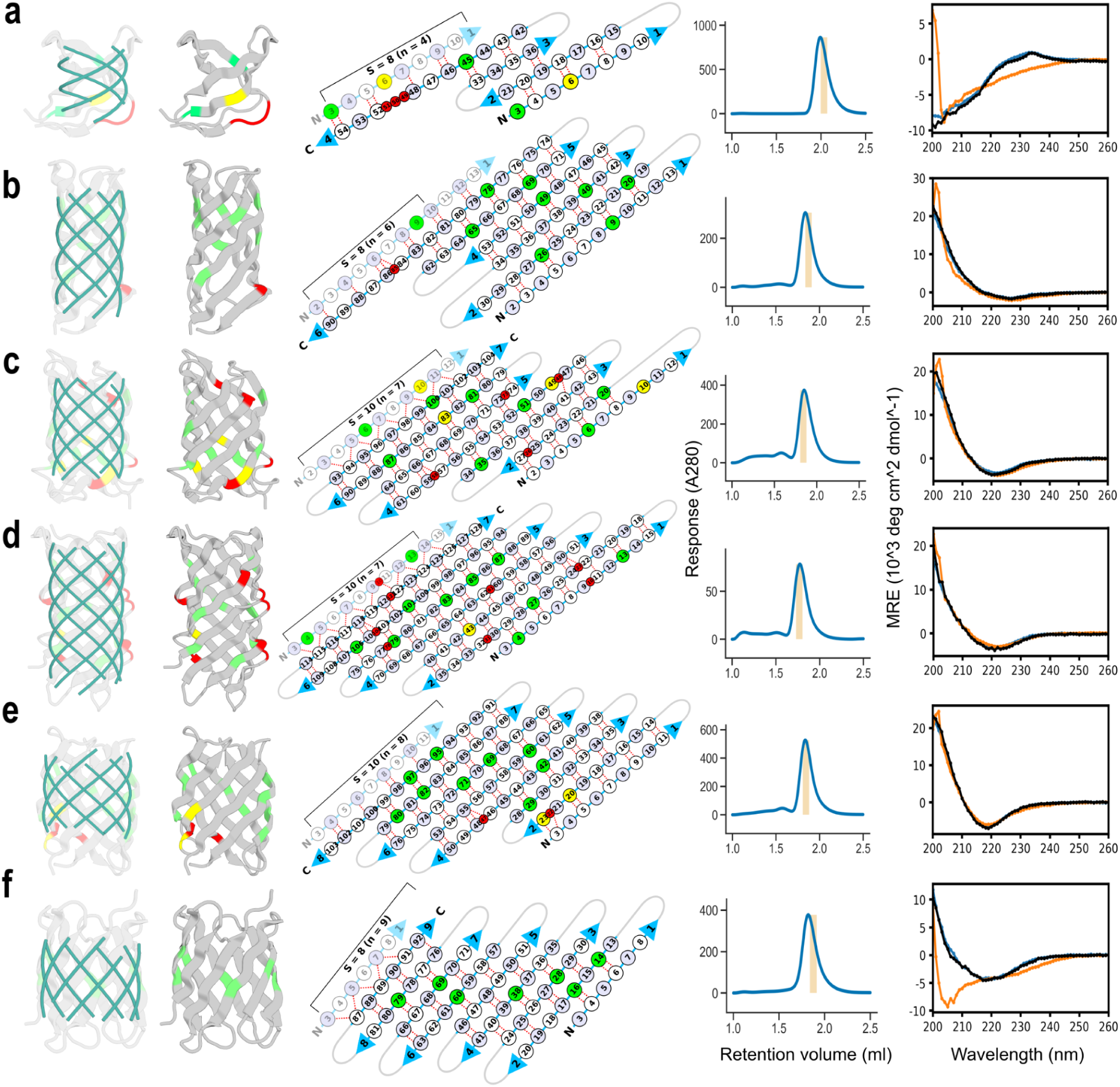
Experimental characterization of representative designs. Columns from left to right display parametric cylinders used as inputs for RFjoint2 inpainting overlaid on resulting designs (translucent), backbone cartoon traces of each design with residue features important for proper barrel formation highlighted using the following color code: red for beta-bulges, yellow for glycines with positive phi backbone torsion angles (e.g. glycine kinks) and green for remaining glycines, 2D beta-strand pairing maps using the same color code, SEC traces of IMAC purified designs with expected elution volumes for monomers highlighted with vertical bars, and far-UV CD spectra at 25°C (blue), 95°C (red), and back to 25°C (black). Beta strand residues in the 2D map are represented as circles shaded light blue and unshaded for exposed and core positions, respectively, and small red circles for beta-bulges. Backbone hydrogen bonds between strands are represented as broken lines. The parameters n (strand count), S (shear), and strand length used to generate the input cylinders for each design are **a**) 4, 10, 9, **b**) 6, 8, 10 **c**) 7, 10, 10, **d**) 7, 10, 13, **e**) 8, 12, 8, and **f**) 9, 10, 6, respectively.

6 designs were chosen for crystallographic screening based on protein expression levels and parameter representation and 2 designs produced crystals providing diffraction data with less than 3.5 Å resolution. We succeeded in determining the structure of one design solved at 2 Å resolution, BBn6 (6 strands, shear number 8, and strand length 10; Figures 2b and 3), whose solution structure was nearly identical to the computational design with a Cɑ rmsd of 0.9 Å and 1.9 Å to the input parametric cylinder (Figure 3b,c). Nearly all the designed core side chain conformations closely matched the crystal structure. The 2-dimensional beta strand pairing map and backbone hydrogen bonds specifying the barrel of the crystal structure matched closely to the design with identical shear number, and a beta-bulge at position 85 (Figure 3a). CD spectra indicated the protein was stable at 95°C (Figure 2b). Structure comparison searches using FoldSeek^29^ indicated that the design’s simple 6 stranded up-and-down barrel topology was not represented in the PDB and AlphaFold models databases^27,28^ (a total of 200 million structures), however, 5 models from the MIGNIFY_ESM30 microbiome database and 1 from the GMGCL_ID microbial metagenomic database had similar topologies with TM-scores > 0.6. Querying the designed sequence against the non-redundant protein sequence database (nr) using PSI-BLAST^30^ with an e-value threshold of 0.05 resulted in no significant similarity. The experimental structural validation of design BBn6 suggests that the 30 designs which pass stringent in silico selection metrics including AlphaFold2 (AF2)^27^ validation, form monomers by SEC, and have beta secondary structure by CD are likely to match their designed 3-dimensional structures; these designs span the full range of barrel topologies with n = 4,6-9 and S = 8,10,12,14.

**Figure 3:**
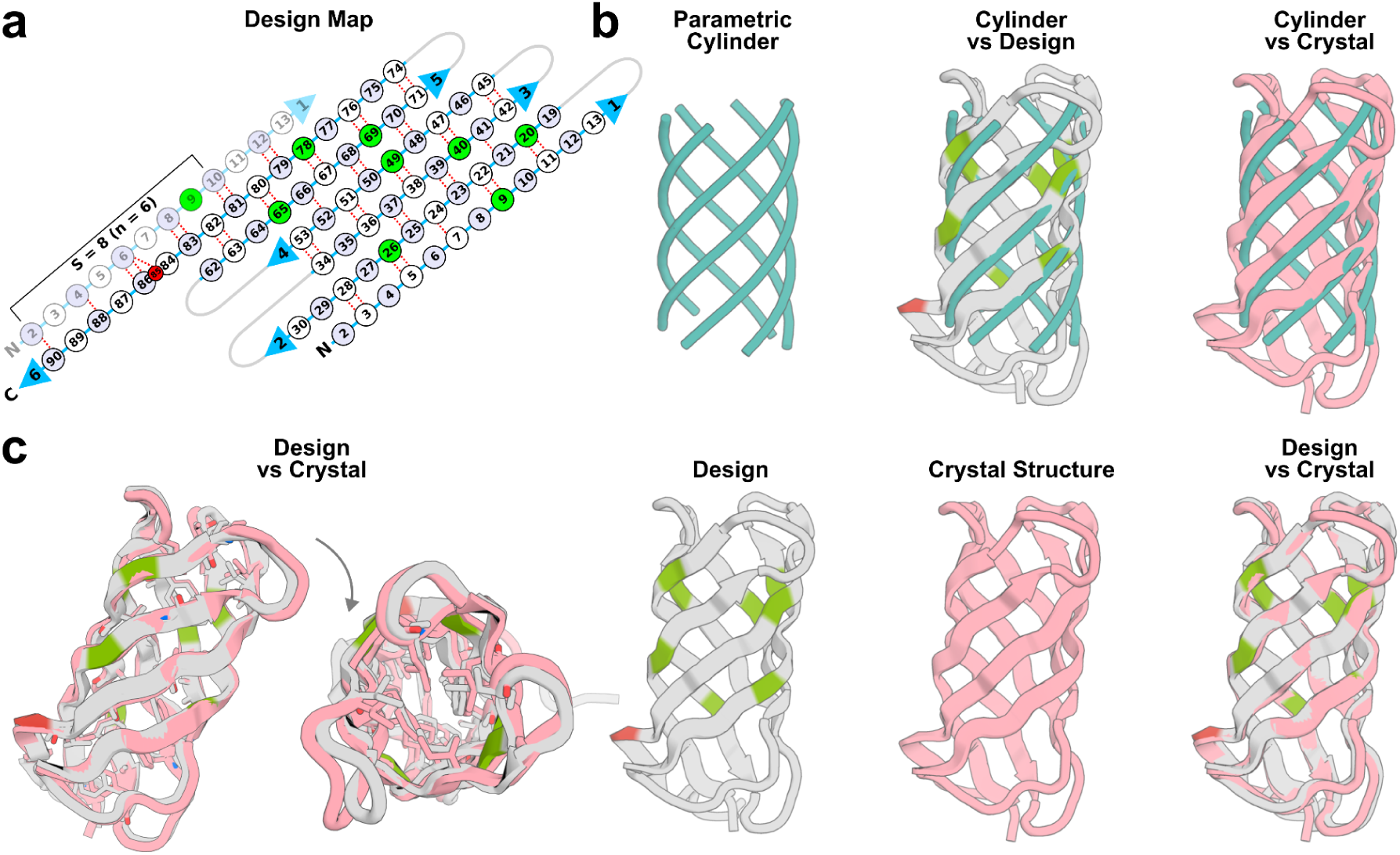
X-ray crystal structure of design BBn6 closely matches the design model. **a**) 2D beta-strand pairing map as described in Figure 2. **b**) left: parametric cylinder generated with n, S and strand length parameters 6, 8, and 10, respectively. Superposition of the parametric cylinder on the design model (middle) and crystal structure (right). **c**) Superposition of the design model with the crystal structure with Cɑ rmsd of 0.9 Å and 1.9 Å to the input parametric cylinder. Note the presence of a beta bulge (red) at position 85 in both the design and crystal structure.

We next explored parametric design of beta barrel nanopores. Protein nanopore engineering has primarily involved making small numbers of mutations in naturally occurring pore-forming proteins, and hence the range of applications has been limited to molecules with roughly the same sizes and diameters of these natural pores. Using a traditional “blueprint” based approach, we have successfully designed de novo beta barrel nanopores with different sizes and shapes^10^, however, the process to generate large beta barrel backbones with control over key features such as beta bulges and register shifts between adjacent strands is difficult and requires significant human expertise. We explored the design of transmembrane beta barrels using the combination of global parametric backbone generation and RFjoint2 described above. We first generated backbones using the aforementioned strategy spanning a wide range of overall shapes with 12, 14 and 16 beta strands. For sequence design^10^, we used a previously described approach which favors hydrophobic residues on the surface of the beta barrels and mostly hydrophilic residues within the pore lining. To compare pore conductances with our previously generated beta-barrel pores made using Rosetta blueprint approaches, we made pore backbones with 12 and 14 strands and a combination of different shear numbers and number of residues per strand. Sequences were designed using ProteinMPNN and Rosetta as described previously^10^. After filtering with AF2 (for predicted structure matching the design model) and selecting for sequences with low beta-sheet propensities (calculated using Raptor-X as described previously^9^), we selected 81 designs for experimental characterization.

19 of the 81 designs were found to express at high levels in E coli and purified from inclusion bodies as stated previously^9,10^. 10 of those designs had SEC peaks at the expected volume fractions. Upon dilution to low nM concentrations, spontaneous insertions for all ten designs into planar lipid bilayer membranes were observed with 3 12-stranded and 2 14-stranded pores showing consistent single channel conductance jumps akin to native pore-forming proteins. Current traces and estimated conductances based on smallest current jumps for representative designs are shown in Figure 4a,b. The 12-stranded designs had conductances between 200-300 pS which were observed for different shapes of 12-stranded pores made using the Rosetta blueprint approach^10^. The single-channel conductances for the RFjoint2 generated 14-stranded designs (370,420 pS) were also close to that observed for one of the 14-stranded designs (470 pS) from our previous study.

**Figure 4:**
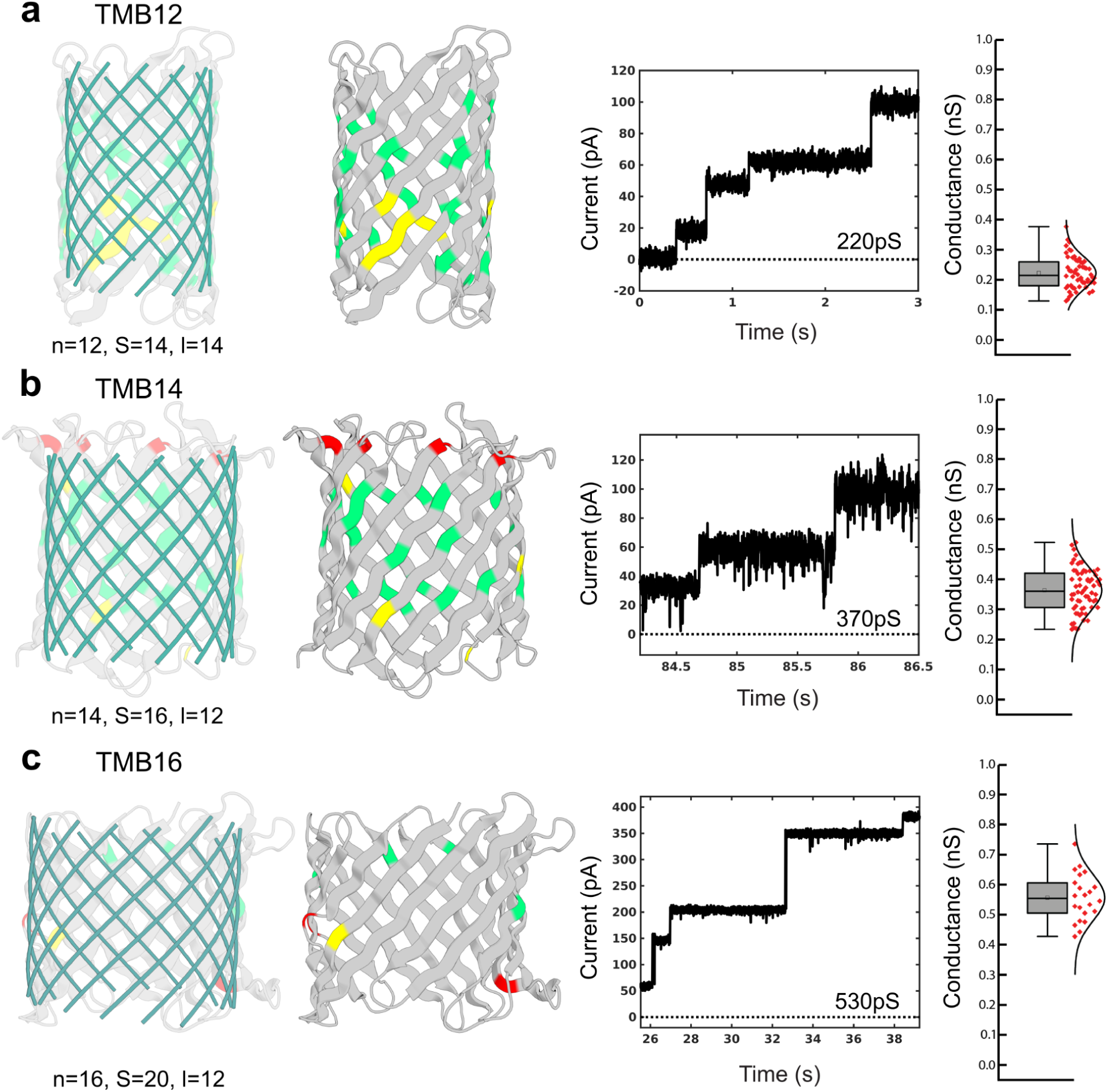
Design of transmembrane beta-barrel pores with different sizes and conductances. From left to right, example input parametric cylinders, designs with features colored using the same scheme as in Figure 2, corresponding current traces at 100mV on a planar bilayer setup, and conductance histograms of three different transmembrane beta-barrel pores **a**) TMB12 **b**) TMB14 and **c**) TMB16. The parameters used for generating the backbones are n (number of strands), S (shear number) and l (number of residues per strand). Current traces shown were recorded at 10kHz and downsampled to 1 kHz and plotted to show multiple current jumps indicating spontaneous pore insertions. Mean conductances calculated from lowest current jumps are indicated in the current trace plot insets. All measurements were carried out in 500mM Potassium Chloride at 100mV constant applied voltage. For the I/V plots, the voltage was varied alternatively in steps of 10 from −100 mV to 100mV.

Motivated by this initial success, we investigated the generation of larger 16 stranded beta-barrel pores and observed higher fidelity of RFjoint2 outputs to the input parametric backbone coordinates possibly due to lower curvature of individual beta-strands in this larger size regime. Following the sequence design, we tested 32 16-stranded designs and found 8 designs that had peaks at the expected SEC fractions. Conductance experiments yielded 3 designs that showed spontaneous insertion and multiple conductance jumps, an example of which is shown in Figure 4c. Unlike the 12 and 14 stranded designs, the conductance jumps for the 16-stranded pores are equally distributed between the potential single-channel jump amplitude and integer multiples of this value, perhaps, reflecting joint insertion of multiple barrels into the planar lipid bilayer membrane. The ∼2 nm diameter and greater than 500pS conductance of these designs are considerably larger than achieved with the previously described Rosetta approach which requires expert placement of glycine kinks and bulges.

## Discussion

The central hypothesis guiding this work was that deep-learning design methods such as RFdiffusion and RFjoint2 have a sufficiently rich understanding of protein folding to be able to refine physically non-realizable inputs into designable outputs. We chose to test this hypothesis on beta barrels because they have traditionally been very challenging to design computationally. The extensive presence of backbone abnormalities in the outputs from both methods (glycine kinks, beta bulges), together with the experimental characterization of folding and the x-ray crystal structure validate our central hypothesis. Both these models have implicitly learned the “rules” required for beta barrel folding that previously had to be empirically discovered one by one. By providing simple-to-generate cylinder inputs to RFdiffusion or RFjoint2, one can now generate beta barrels with unprecedented speed and success rates (> 50% in silico success in many cases) across a broad range of barrel parameters.

Control of the beta barrel pore radius and geometry is of vital importance for downstream applications. The beta barrel central cavity can host small molecule binding/catalytic sites^6,31,32^ and can function as transmembrane nanopores for DNA sequencing and other applications^10,9,33^. The methods described here now permit the design of beta barrels with precisely tunable overall shapes rapidly and without requiring specialist knowledge. The ability of RFdiffusion to generate symmetric oligomers, coupled with the approach described here, should permit the design of transmembrane oligomeric pores, which could be designed to assemble in response to a specific trigger. More broadly, for folds such as beta solenoids and beta propellers, that are still challenging to design de novo^34–36^, the approaches described here could enable design without the need to train dedicated neural networks or to explicitly determine the “rules” of folding such structures. Our design of a 16 stranded beta barrel nanopore with observed single-channel conductance of 555 pS, considerably larger than achieved with the previous Rosetta based approach, highlights the power and simplicity of using global parameterization to define overall protein shape and deep learning generative modeling to fill in the fold specific features. Further development of current AI models is needed to make robust beta-barrels in the larger size range such as the TMB16s owing to the long range interactions which are harder to encode at the backbone and sequence level.

Both RFjoint2 and RFdiffusion are fine-tuned directly from the RoseTTAFold structure prediction network, and thereby directly leverage the representations learned during structure prediction training, without requiring the extensive compute required for initial training. In this study, we use both of these design methods in a manner tangential to the tasks for which they were trained, in a similar spirit to previous work repurposing existing protein networks without retraining^12,13,15,37,38^. In a similar manner to how large language models have been used as the foundation for domain-specific fine-tuned models^39^, protein structure prediction models are emerging as foundation models for protein design.

## Materials and Methods

### Generation of parameter-defined cylinders

Input cylinder backbones for both RFjoint2 and RFdiffusion were generated using a geometric model of an intertwined coil where each coil represents a beta-strand as previously described^40^. The tilt of each beta-strand and the radius of the cylinder with respect to the central axis were computed from the strand count (n) and shear number (S), and the placement of C-alpha atoms within each coil was determined by inter-strand H-bond geometry. The complete backbone was generated given the C-alphas as input using BBQ^22^. For RFdiffusion, 2 residue N and C termini and 3 residue loops connecting the strands were generated for each cylinder using PyRosetta^25^. The script used to generate barrel cylinders and backbones is available on Github (https://github.com/davidekim/parametric_barrels).

### Design with RFjoint2 and selection for experimental characterization

To generate parameter-specified barrels with RFjoint2, the parametrically-defined cylinders (without added loops) were provided to RFjoint2 along with an intermediate template-confidence value (0.9). The barrel strands were provided interspersed with loops (at both the N- and C-termini, and in between the strands) of specified length ranges (0-4 residues at N- and C-termini, and 3-6 residues between strands). Because RFjoint2 is deterministic (until the random-order autoregressive sequence design), for each design for a specific parameter set, loop and termini lengths were randomly and independently sampled from the provided length ranges.

Parameter ranges investigated for experimental characterization of soluble barrels were 4-9 strands (n), 8-18 shear values (S), and strand lengths of 5-20 residues. For each parameter group, 100 inpainted designs were generated and structures with closed barrels based on backbone hydrogen bonding were selected for ProteinMPNN sequence (re)design followed by AF2 validation (See In silico analysis of barrel design performance below). 96 designs were selected for experimental characterization based on the Cɑ rmsd of the design to the AF2 model (< 0.985 Å) and the input cylinder (<5.0 Å), AF2 pLDDT (>90), net charge at pH 8 (absolute value > 2), and the spatial aggregation propensity (SAP) score^41^ (< 39).

### Design with RFdiffusion

To generate parameter-specified barrels with RFdiffusion, the parametrically-defined cylinders (with added loops and termini) were provided to RFdiffusion, and 50 steps of noise were added to the input structure, before denoising. Note that in this paper we use T=200 as the total number of timesteps defining the full forward noising process (the schedule RFdiffusion is trained with). Hence, 50 steps of noise addition represents a quarter of the noising trajectory. Empirically, RFdiffusion performs as well with a noising schedule of T=50 timesteps (giving a 4X computational speedup)^17^, so the partial-diffusion approach described here could likely be performed with an equivalent speedup (with 12 steps of noise added with T=50). All other parameters matched the RFdiffusion default parameters. Because RFdiffusion is not deterministic, for all analyses in this paper, fixed loop lengths were used for each set of input parameters (2 residues at N- and C-terminus, and 3 residues between adjacent strands).

### In silico analysis of barrel design performance

Downstream in silico analysis of design performance was performed essentially as described previously^13,17^. Briefly, ProteinMPNN was used to (re)design the protein sequences (8 sequences per backbone; temperature = 0.1), and AF2^27^ *model_4_ptm* was used in single sequence mode to subsequently predict the structure of that designed sequence. The AF2 confidence (pLDDT) and the root mean square deviation (RMSD) between the AF2 prediction and the design model was calculated.

The definition of “in silico success” closely followed previous work^13,17^, with the AF2 pLDDT > 80 and RMSD between AF2 prediction and design < 2 Å. Additionally, to check for barrel construction and closure, we assessed backbone hydrogen bonding between adjacent strands, and required at least 10 percent of the strand length parameter participating in hydrogen bonding between each pair of strands (including the N and C terminal strands to complete barrel closure), both in the design model and AF2 prediction. Because ProteinMPNN is used to design the sequence on the backbone, we used Rosetta FastRelax^18^ to relax the design backbones with ProteinMPNN sequence, as well as the AF2 predictions prior to detecting these hydrogen bonds. The script used to check the extent of barrel formation and closure is available on Github (https://github.com/davidekim/parametric_barrels).

### Protein expression and purification

RFjoint2 designs were synthesized as eBlock gene fragments by Integrated DNA Technologies (IDT), and cloned into the pET29b+ vector using Golden Gate assembly. Plasmids were transformed into chemically competent Lemo21(DE3) *E. coli* (NEB) and protein expression for each design was carried out overnight at 37 ℃ in 1mL/well of autoinduction medium consisting of Terrific Broth II (MP biomedicals), 2 mM MgS04, 1x 5052, and 30 µg/mL kanamycin in a 96-deep well round-bottom 2 mL/well plate on a platform shaker (1000 rpm). Cells were lysed using BugBuster (Millipore) in the presence of DNAse and protease inhibitors, and immobilized metal affinity chromatography (IMAC) purification proceeded in a 96-well (800 µL/well) plate with a 24-µm polyethylene frit (Agilent Technologies) loaded with 75 µL/well Ni-NTA affinity resin (Qiagen). The eluted purified protein was exchanged into 20 mM Tris buffer (150 mM NaCl, pH 8.0) and further purified by Akta Pure FPLC (GE Healthcare) SEC using a Superdex 75 increase 5/150 GL column. Large scale expressions for crystallography screens were performed in 2L baffled culture flasks using 500mL of autoinduction medium. Following cell lysis by sonication, the 6x Histidine tag was removed by SNAC-tag chemical cleavage^42^ and further purified by SEC using a Superdex 75 increase 10/300 GL column. Fractions for monodispersed peaks that were monomeric based on calibration standards were collected and kept at 4 ℃ for biophysical characterization experiments. Protein concentrations were determined with predicted extinction coefficients^43^ using a NanoDrop spectrophotometer (ThermoScientific) to measure absorbance at 280 nm. Purified proteins were verified using liquid chromatography mass spectrometry (LC/MS).

The membrane pore beta barrels were expressed and purified using auto-induction media. All proteins were washed thoroughly with Triton X-100 and Brij-35 detergents and refolded from inclusion bodies using dodecyl-phosphatidylcholine (DPC) detergent and were subsequently run through an SEC column (Superdex S200 from Cytiva). Appropriate fractions were diluted to nanomolar or picomolar concentrations and tested for membrane insertion and conductance as described below.

### Ionic current measurements in planar lipid bilayer membranes

Ion current measurements were carried out as previously described^10^ using the Nanion Orbit 16TC instrument (https://www.nanion.de/products/orbit-16-tc/) on MECA chips. Lipid stock solutions were freshly made in dodecane at a final concentration of 5mg/mL. Di-phytanoyl-phosphatidylcholine (DPhPC) lipids were used for all experiments. Designed proteins were diluted in a buffer containing 0.05% DPC (∼ 1 critical micelle concentration), 25 mM Tris-Cl pH 8.0 and 150 mM NaCl to a final concentration of ∼100 nM. Subsequently, 0.5 μL or less of this stock was added to the cis chamber of the chip containing 200 μL of buffer while simultaneously making lipid bilayers using the in-built rotating stir-bar setup. All measurements were carried out at 25 ℃ and with a positive potential bias of 100 mV. Spontaneous insertions were recorded over multiple rounds of bilayer formation. All chips were washed with multiple rounds of ethanol and water and completely dried before testing subsequent designs. A 500 mM NaCl buffer was used on both sides of the membrane for all current recordings. Raw signals were recorded at a sampling frequency of 5 kHz. Only current recordings from bilayers whose capacitances were in the range 15-25 pF were used for subsequent analysis. The raw signals at 5 kHz were downsampled to 1000 Hz using an 8-pole bessel filter. Estimation of current jumps were carried out using a custom script with appropriate thresholds. Analysis scripts for processing ion conductance data as presented in this manuscript are also available on Github (https://github.com/sagardipm/denovoPores) and archived in Zenodo (DOI: 10.5281/zenodo.10939541).

### Circular dichroism (CD)

Purified designs were prepared at 0.2-0.4 mg/mL in 20 mM Tris buffer (150 mM NaCl, pH 8.0). Wavelength scans from 200-260 nm were recorded at 25°C, 95°C, and back to 25°C using a Jasco J-1500 CD spectrophotometer with a 1 mm path length cuvette.

### Crystallization and structure determination

All crystallization experiments were conducted using the sitting drop vapor diffusion method. Crystallization trials were set up in 200 nL drops using the 96-well plate format at 20 °C.

Crystallization plates were set up using a Mosquito LCP from SPT Labtech, then imaged using UVEX microscopes and UVEX PS-256 from JAN Scientific. Diffraction quality crystals formed in 0.2 M AmSO4, 30 %(w/v) PEG 8000. Crystals were looped and flash cooled in liquid nitrogen using PEG 200 as cryoprotectant.

Diffraction data was collected at the NSLS2 FMX beamline. X-ray intensities and data reduction were evaluated and integrated using XDS^44^ and merged/scaled using Pointless/Aimless in the CCP4 program suite^45^. Structure determination and refinement starting phases were obtained by molecular replacement using Phaser^46^ using the designed model for the structures. Following molecular replacement, the models were improved using phenix.autobuild^47^; with simulated annealing. Structures were refined in Phenix^47^. Model building was performed using COOT^48^. The final model was evaluated using MolProbity^49^. Data collection and refinement statistics are recorded in Table S3. Data deposition, atomic coordinates, and structure factors reported in this paper have been deposited in the PDB^24^.

## Code availability

See Supplementary information.

## Acknowledgements

We thank Luki Goldschmidt and Patrick Vecchiato for computer and network infrastructure support, Kandise VanWormer for managing the wet labs at the UW Institute for Protein Design (IPD), and the staff at the IPD for help with protein production, LC/MS, and crystal screening. This research used resources of the National Energy Research Scientific Computing Center (NERSC), a U.S. Department of Energy Office of Science User Facility located at Lawrence Berkeley National Laboratory, operated under Contract No. DE-AC02-05CH11231 using NERSC award BER-ERCAP0022018, and the FMX beamline of the National Synchrotron Light Source II, a U.S. Department of Energy (DOE) Office of Science User Facility operated for the DOE Office of Science by Brookhaven National Laboratory under Contract No. DE-SC0012704. The Center for BioMolecular Structure (CBMS) is primarily supported by the National Institutes of Health, National Institute of General Medical Sciences (NIGMS) through a Center Core P30 Grant (P30GM133893), and by the DOE Office of Biological and Environmental Research (KP1607011). This work was supported by the Institute for Protein Design Breakthrough Fund for “De novo design of selective pores” (D.E.K., S.M. and D.B.)

## Author Contributions

D.E.K. conceived the original idea; D.E.K., J.L.W., D.J., S.M. and D.B. designed research; D.E.K., J.L.W., D.J., S.M, R.S., S.R.G, A.K, and A.K.B. performed research; D.E.K., J.L.W., D.J., S.M. and A.K.B analyzed data; and D.E.K., J.L.W., D.J., S.M and D.B. wrote the paper.

## References

1. Crick, F. H. C. The Fourier transform of a coiled-coil. Acta Crystallogr. 6, 685–689 (1953).

2. Huang, P.-S. et al. High thermodynamic stability of parametrically designed helical bundles. Science 346, 481–485 (2014).

3. Grigoryan, G. & DeGrado, W. F. Probing Designability via a Generalized Model of Helical Bundle Geometry. J. Mol. Biol. 405, 1079–1100 (2011).

4. Grigoryan, G. et al. Computational design of virus-like protein assemblies on carbon nanotube surfaces. Science 332, 1071–1076 (2011).

5. Chen, Z. et al. Programmable design of orthogonal protein heterodimers. Nature 565, 106–111 (2019).

6. Dou, J. et al. De novo design of a fluorescence-activating β-barrel. Nature 561, 485–491 (2018).

7. Kim, D. E., et al. De novo design of small beta barrel proteins. Proc. Natl. Acad. Sci. 120, e2207974120 (2023).

8. Liu, W. M. Shear numbers of protein beta-barrels: definition refinements and statistics. J. Mol. Biol. 275, 541–545 (1998).

9. Vorobieva, A. A. et al. De novo design of transmembrane β barrels. Science 371, eabc8182 (2021).

10. Berhanu, S. et al. Sculpting conducting nanopore size and shape through de novo protein design. 2023.12.20.572500 Preprint at 10.1101/2023.12.20.572500 (2023).

11. Salemme, F. R. Conformational and geometrical properties of beta-sheets in proteins. III. Isotropically stressed configurations. J. Mol. Biol. 146, 143–156 (1981).

12. Anishchenko, I. et al. De novo protein design by deep network hallucination. Nature 600, 547–552 (2021).

13. Wang, J. et al. Scaffolding protein functional sites using deep learning. Science 377, 387–394 (2022).

14. Wicky, B. I. M. et al. Hallucinating symmetric protein assemblies. Science 378, 56–61 (2022).

15. Frank, C. et al. Efficient and scalable de novo protein design using a relaxed sequence space. 2023.02.24.529906 Preprint at 10.1101/2023.02.24.529906 (2023).

16. Torres, S. V. et al. De novo design of high-affinity binders of bioactive helical peptides. Nature (2023) doi:10.1038/s41586-023-06953-1.

17. Watson, J. L. et al. De novo design of protein structure and function with RFdiffusion. Nature 620, 1089–1100 (2023).

18. Leaver-Fay, A. et al. ROSETTA3: an object-oriented software suite for the simulation and design of macromolecules. Methods Enzymol. 487, 545–574 (2011).

19. Lasters, I., Wodak, S. J., Alard, P. & van Cutsem, E. Structural principles of parallel beta-barrels in proteins. Proc. Natl. Acad. Sci. U. S. A. 85, 3338–3342 (1988).

20. Murzin, A. G., Lesk, A. M. & Chothia, C. Principles determining the structure of β-sheet barrels in proteins I. A theoretical analysis. J. Mol. Biol. 236, 1369–1381 (1994).

21. Murzin, A. G., Lesk, A. M. & Chothia, C. Principles determining the structure of beta-sheet barrels in proteins. II. The observed structures. J. Mol. Biol. 236, 1382–1400 (1994).

22. Gront, D., Kmiecik, S. & Kolinski, A. Backbone building from quadrilaterals: a fast and accurate algorithm for protein backbone reconstruction from alpha carbon coordinates. J. Comput. Chem. 28, 1593–1597 (2007).

23. Anand, N. & Achim, T. Protein Structure and Sequence Generation with Equivariant Denoising Diffusion Probabilistic Models. Preprint at 10.48550/ARXIV.2205.15019 (2022).

24. Berman, H. M. et al. The Protein Data Bank. Nucleic Acids Res. 28, 235–242 (2000).

25. Chaudhury, S., Lyskov, S. & Gray, J. J. PyRosetta: a script-based interface for implementing molecular modeling algorithms using Rosetta. Bioinforma. Oxf. Engl. 26, 689–691 (2010).

26. Dauparas, J. et al. Robust deep learning-based protein sequence design using ProteinMPNN. Science 378, 49–56 (2022).

27. Jumper, J. et al. Highly accurate protein structure prediction with AlphaFold. Nature 596, 583–589 (2021).

28. Varadi, M. et al. AlphaFold Protein Structure Database: massively expanding the structural coverage of protein-sequence space with high-accuracy models. Nucleic Acids Res. 50, D439–D444 (2022).

29. Kempen, M. van et al. Foldseek: fast and accurate protein structure search. 2022.02.07.479398 Preprint at 10.1101/2022.02.07.479398 (2022).

30. Altschul, S. F. et al. Gapped BLAST and PSI-BLAST: a new generation of protein database search programs. Nucleic Acids Res. 25, 3389–3402 (1997).

31. Kipnis, Y. et al. Design and optimization of enzymatic activity in a de novo β-barrel scaffold. Protein Sci. 31, e4405 (2022).

32. An, L. et al. De novo design of diverse small molecule binders and sensors using Shape Complementary Pseudocycles. 2023.12.20.572602 Preprint at 10.1101/2023.12.20.572602 (2023).

33. Shimizu, K. et al. De novo design of a nanopore for single-molecule detection that incorporates a β-hairpin peptide. Nat. Nanotechnol. 17, 67–75 (2022).

34. MacDonald, J. T. et al. Synthetic beta-solenoid proteins with the fragment-free computational design of a beta-hairpin extension. Proc. Natl. Acad. Sci. U. S. A. 113, 10346–10351 (2016).

35. Marcos, E. et al. De novo design of a non-local β-sheet protein with high stability and accuracy. Nat. Struct. Mol. Biol. 25, 1028–1034 (2018).

36. Voet, A. R. D. et al. Computational design of a self-assembling symmetrical β-propeller protein. Proc. Natl. Acad. Sci. U. S. A. 111, 15102–15107 (2014).

37. Roney, J. P. & Ovchinnikov, S. State-of-the-Art Estimation of Protein Model Accuracy Using AlphaFold. Phys. Rev. Lett. 129, 238101 (2022).

38. Rettie, S. A. et al. Cyclic peptide structure prediction and design using AlphaFold. BioRxiv Prepr. Serv. Biol. 2023.02.25.529956 (2023) doi:10.1101/2023.02.25.529956.

39. Bommasani, R., et al. On the Opportunities and Risks of Foundation Models. arXiv.org https://arxiv.org/abs/2108.07258v3 (2021).

40. Naveed, H., Xu, Y., Jackups, R. Jr. & Liang, J. Predicting Three-Dimensional Structures of Transmembrane Domains of β-Barrel Membrane Proteins. J. Am. Chem. Soc. 134, 1775–1781 (2012).

41. Chennamsetty, N., Voynov, V., Kayser, V., Helk, B. & Trout, B. L. Design of therapeutic proteins with enhanced stability. Proc. Natl. Acad. Sci. 106, 11937–11942 (2009).

42. Dang, B. et al. SNAC-tag for sequence-specific chemical protein cleavage. Nat. Methods 16, 319–322 (2019).

43. Pace, C. N., Vajdos, F., Fee, L., Grimsley, G. & Gray, T. How to measure and predict the molar absorption coefficient of a protein. Protein Sci. 4, 2411–2423 (1995).

44. Kabsch, W. XDS. Acta Crystallogr. D Biol. Crystallogr. 66, 125–132 (2010).

45. Winn, M. D. et al. Overview of the CCP4 suite and current developments. Acta Crystallogr. D Biol. Crystallogr. 67, 235–242 (2011).

46. McCoy, A. J. et al. Phaser crystallographic software. J. Appl. Crystallogr. 40, 658–674 (2007).

47. Adams, P. D. et al. PHENIX: a comprehensive Python-based system for macromolecular structure solution. Acta Crystallogr. D Biol. Crystallogr. 66, 213–221 (2010).

48. Emsley, P. & Cowtan, K. Coot: model-building tools for molecular graphics. Acta Crystallogr. D Biol. Crystallogr. 60, 2126–2132 (2004).

49. Williams, C. J. et al. MolProbity: More and better reference data for improved all-atom structure validation. Protein Sci. Publ. Protein Soc. 27, 293–315 (2018).

50. Baek, M. et al. Accurate prediction of protein structures and interactions using a three-track neural network. Science 373, 871–876 (2021).

