## Supplementary Information for "Parametrically guided design of beta barrels and transmembrane nanopores using deep learning"

**This PDF file includes:**

Supplementary text
Figures S1 to S6
Tables S1 to S3
SI References

**Other supplementary materials for this manuscript include the following:**

RFjoint2 code is available to download for reviewers to assess:

http://files.ipd.uw.edu/pub/RFjoint2/da39a3ee5e6b4b0d3255bfef95601890afd80709/proteininpainting.tar.gz

http://files.ipd.uw.edu/pub/RFjoint2/da39a3ee5e6b4b0d3255bfef95601890afd80709/InpaintingApril22.pt

Code will be prepared for public release in the near future, and before publication of this manuscript.

RFdiffusion code is available at:

https://github.com/RosettaCommons/RFdiffusion

Utility scripts to generate input barrel backbones, RFjoint2 and RFdiffusion refinement commands, and to determine the extent of barrel closure are available at:

https://github.com/davidekim/parametric_barrels

Analysis scripts for processing ion conductance data are available on Github and archived in Zenodo.

https://github.com/sagardipm/denovoPores
DOI: 10.5281/zenodo.10939541

### Supplementary text

#### RFjoint2

We explored whether we could use RoseTTAFold based inpainting[1] for the design of beta barrels starting from global geometric descriptions. We trained an improved version of RFjoint; RFjoint2, which outperforms the original RFjoint model at motif scaffolding (Figure S1) and has not been described elsewhere. Several key improvements underpin RFjoint2 and are described in the methods section below. Firstly, RFjoint2 trains from a later version of RoseTTAFold, which was the first RoseTTAFold version to explicitly model sidechains (the original RF and RFjoint models only predicted backbone atoms), parameterized by torsion angles[2]. Secondly, in contrast to RFjoint, where sequence was decoded in “one-shot” by taking the most probable amino acid at each position, sequence in RFjoint2 is decoded iteratively over many “recycles”.

On the benchmark set of motif-scaffolding problems introduced in the RFdiffusion manuscript[4], RFjoint2 sequences significantly improve over RFjoint sequences (higher success rates in 14 cases, and one tie) (Table S1). With ProteinMPNN sequence re-design of the designed scaffold, RFjoint2 “solves” 21 of the benchmarking problems (vs 23 in RFdiffusion and 17 in RFjoint), with higher in silico success rates than RFjoint in 15 of these cases. While RFjoint2 still generally performs worse than RFdiffusion, RFjoint2 outperforms RFdiffusion on 6 of the benchmarking problems (Table S1).

### Supplementary methods

#### RFjoint2 Training

RFjoint2 was trained from an unpublished developmental version of RoseTTAFold2[7] (RF2). For clarity, we name this network RF-intermediate. This network is not a contribution of this work. For context, we provide a brief overview of the training of this network below.

##### RoseTTAFold Structure Prediction Training

The RF architecture used for RFjoint2 differs significantly from the original RF architecture[2]. Notably, unlike the original RF model, RF-intermediate explicitly models amino acid sidechains, using the parameterization in AF2 and RF2[6,7]. Unlike RF2 and AF2, RF-intermediate was trained only on the PDB (without an AF2 distillation set). Following AF2[6], the frame-aligned point error (FAPE) loss is the primary structural loss used during training.

Hyperparameters are detailed below:

- Training set: PDB set only (RoseTTAFold training set), 21120 examples per epoch
- Optimizer: AdamW[8] optimizer with default parameters in PyTorch[9]
- Batch size: 64 examples per batch
- Learning rate: 0.0005
- Linear warm-up for 16000 optimization steps, followed by;
- Linear decay for 200000 optimization steps

- Crop size: 300 amino acids
- Number of seed MSA sequences: 128
- Number of extra MSA sequences: 2048
- 300 epochs of training

The losses used were the following, and are fully described in the RF2 manuscript[7].

- Distogram loss ($L_{dist}$); 1.0
- FAPE loss ($L_{FAPE}$); 10.0 (80:20 backbone only vs full-atom)
- LDDT loss ($L_{lddt}$); 0.1
- MSA prediction loss ($L_{MLM}$); 3.0

##### RFjoint2 Training Parameters

RFjoint2 was fine-tuned directly from the RF-intermediate weights, and in an otherwise identical manner to the original RFjoint model[1], using the same training tasks (structure prediction, fixed-backbone sequence prediction and joint sequence-structure inpainting) and masking strategy. RFjoint2 was trained using the same losses as RF-intermediate, except for the addition of a clash loss ($L_{clash}$), described elsewhere[7], with weight 1.0.

Training hyperparameters are detailed below:

- Three training tasks (as in RFjoint[1]); Structure prediction (as in RF), fixed-backbone sequence design, joint sequence structure “inpainting”. Sampled with equal probability.
- During fixed-backbone sequence design training, 90-100% of residues were masked.
- During inpainting training, 10-35 contiguous residues were masked, flanked by 3-6 residues either side, where a loss was not applied (see RFjoint[1]).
- Training set: PDB set only (RoseTTAFold training set), 21120 examples per epoch
- Optimizer: AdamW[8] optimizer with default parameters in PyTorch[9]
- Batch size: 512 examples per batch
- Learning rate: 0.0005
- Linear warm-up for 16000 optimization steps, followed by;
- Linear decay for 200000 optimization steps
- Crop size: 260 amino acids
- 50 epochs of training

#### RFjoint2 Inference

While the training of RFjoint2 very closely matched the training of RFjoint, the inference strategy deviated significantly. The details of the changes are described below.

##### Recycling

RF, following AF2, uses “recycling” during structure prediction to iteratively refine a prediction (by re-running prediction with the latent prediction from the previous recycle). In RFjoint, recycling was also used, typically running 5 recycles during a prediction. However, in RFjoint2 we recycle for significantly longer. By applying the $L_{lddt}$ loss during RFjoint2 training, we handily maintain a “confidence” parameter that we can use to monitor the progress of an RFjoint2 trajectory. Because this loss penalizes the model’s assessment of how far the “inpainted” structure is from the true structure, the RFjoint2 pLDDT parameter correlates with structure quality. Through recycling, the pLDDT typically increases, and hence, we recycle until the pLDDT has plateaued (no improvement for 10 recycles). This often leads to in excess of 50 recycles. At this point, we begin autoregressive sequence design (see below).

##### Autoregressive sequence design

In RFjoint, the amino acid sequence was generated “one-shot”, by taking the most probable amino acid predicted at each position. Inspired by the iterative decoding in ProteinMPNN[3], in RFjoint2, after the pLDDT has plateaued (see above), we iteratively “decode” sequence, by sampling from the softmaxed-logit probabilities at a random position at each recycle. This position then gets fixed and input as “true” sequence to the next recycle. Because sequence design is a locally-defined problem[3], we simultaneously decode multiple positions at a time (through a greedy search permitting decoding of residues > 15 Å to be simultaneously decoded). This leads to a significant inference speed up. For all experiments, the softmax temperature for sampling was 0.1, following the default in ProteinMPNN. The random decoding order also creates stochasticity in the outputs of RFjoint2, in comparison to the deterministic inference used in RFjoint. This order-agnostic autoregressive sequence design strategy nearly matches the quality of sequences generated with fixed-backbone sequence design using ProteinMPNN[3], but we found it most expedient to generate 8 ProteinMPNN sequences per RFjoint2-inpainted backbone[4,5], and choose the sequence with the closest recapitulation of the design model according to AlphaFold2 (AF2)[6] structure prediction.

#### SI References


1. Wang, J. *et al.* Scaffolding protein functional sites using deep learning. *Science* **377**, 387–394 (2022).
2. Baek, M. *et al.* Accurate prediction of protein structures and interactions using a three-track neural network. *Science* **373**, 871–876 (2021).
3. Dauparas, J. *et al.* Robust deep learning-based protein sequence design using ProteinMPNN. *Science* **378**, 49–56 (2022).
4. Watson, J. L. *et al.* De novo design of protein structure and function with RFdiffusion. *Nature* **620**, 1089–1100 (2023).
5. Wu, K. E. *et al.* Protein structure generation via folding diffusion. Preprint at

https://doi.org/10.48550/arXiv.2209.15611 (2022).

6. Jumper, J. *et al.* Highly accurate protein structure prediction with AlphaFold. *Nature* **596**, 583–589 (2021).
7. Baek, M. *et al.* Efficient and accurate prediction of protein structure using RoseTTAFold2. 2023.05.24.542179 Preprint at https://doi.org/10.1101/2023.05.24.542179 (2023).
8. Loshchilov, I. & Hutter, F. Decoupled Weight Decay Regularization. *arXiv.org* https://arxiv.org/abs/1711.05101v3 (2017).
9. Paszke, A. *et al.* PyTorch: An Imperative Style, High-Performance Deep Learning Library. *arXiv.org* https://arxiv.org/abs/1912.01703v1 (2019).
10. Yim, J. *et al.* SE(3) diffusion model with application to protein backbone generation. *arXiv.org* https://arxiv.org/abs/2302.02277v2 (2023).
11. Leaver-Fay, A. *et al.* ROSETTA3: an object-oriented software suite for the simulation and design of macromolecules. *Methods Enzymol.* **487**, 545–574 (2011).

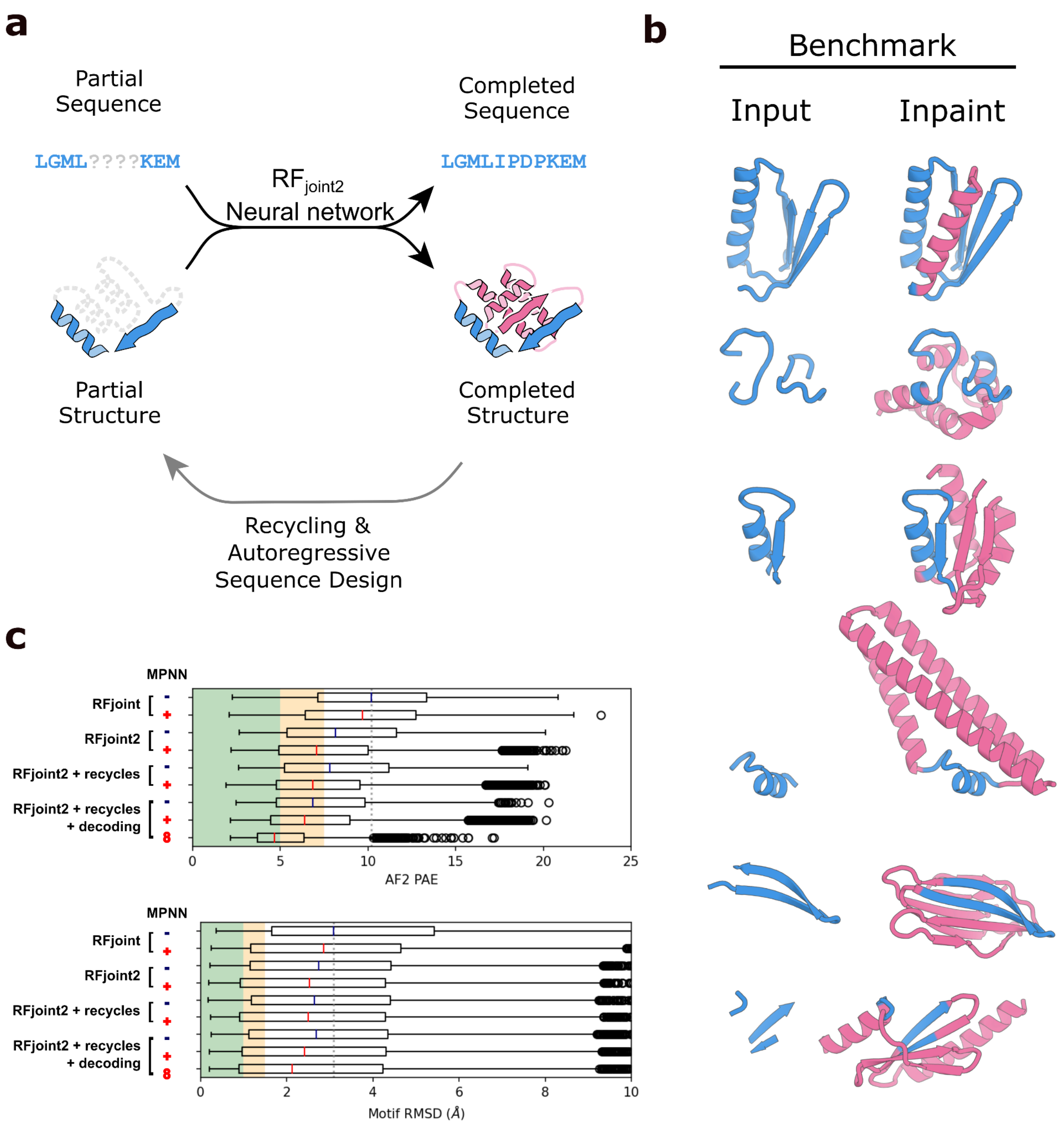


#### **Figure S1**. Overview of RFjoint2 inpainting

**a**) Overview of RFjoint2. RFjoint2 takes as input the sequence and structure of a “motif” (blue). It then predicts the sequence and structure of the “scaffold” around the motif (pink). Predictions are made over several “recycles”, and the sequence is iteratively decoded in an order-agnostic manner. **b**) A few examples from the RFdiffusion motif-scaffolding benchmark set, where RFjoint2 significantly outperforms RFjoint (examples are *2KL8, 1PRW, 5TPN, 6E6R_long, 5IUS, 5YUI*). Full benchmarking details are listed in Table S1. **c**) Ablation experiment showing the effect of different features in RFjoint2, as evaluated by AF2 confidence (pAE, top) and recapitulation of the motif (bottom). RFjoint2 outperforms RFjoint, with and without

ProteinMPNN sequence redesign of the de novo scaffold. Permitting indefinite recycling in RFjoint2 (until pLDDT plateaus; see Methods) subtly improves RFjoint2 outputs. Furthermore, subsequent order-agnostic autoregressive sequence design improves both the sequences (blue) and backbones (with ProteinMPNN sequences; red) designed by RFjoint2. Sampling 8 sequences in ProteinMPNN and taking the best (in line with other work[4,5,10]), unsurprisingly, leads to improved sequences.

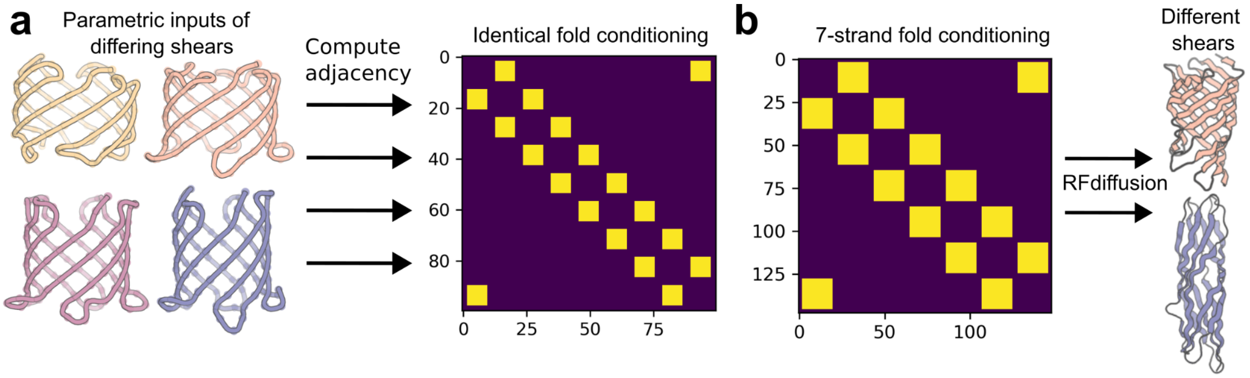


**Figure S2**. RFdiffusion fold conditioning cannot perform parametric beta barrel design.

**a**) Using the published code for RFdiffusion to compute “block adjacency” (fold conditioning) matrices for four parametric beta barrel inputs (left) with identical parameters other than their strand shear yields the same block adjacency matrix (right). Thus suggesting the shear parameter is not encoded by the block adjacency featurization. **b**) When generating 150 outputs with RFdiffusion using a block adjacency matrix consistent with a seven-strand beta barrel topology (left), RFdiffusion generated barrels with inconsistent shear (right).

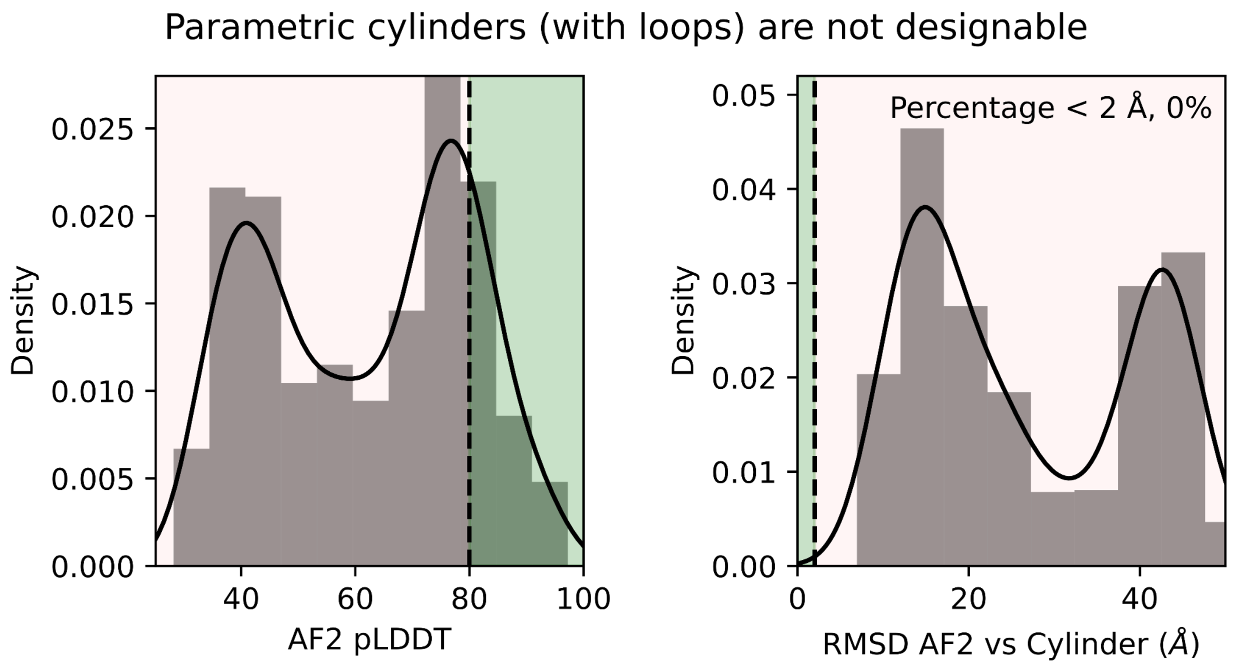


**Figure S3**. Parametric cylinders are not designable.

Without RFdiffusion/RFjoint2 refinement, the parametric backbone cylinders (with loops added) are not designable. Left: AF2 pLDDT is low (mostly below 80; the threshold for success; green area), and none of the AF2 predictions are within 2 Å of the cylinder (right, the threshold for in silico success, green area). $N$ = 928 cylinders with unique n, S and I parameters, spanning the same parameter space as used in Figure 1c and d in the manuscript. For each backbone, 8 ProteinMPNN sequences were sampled, and the "best" sequence plotted (by RMSD between the AF2 prediction and design).

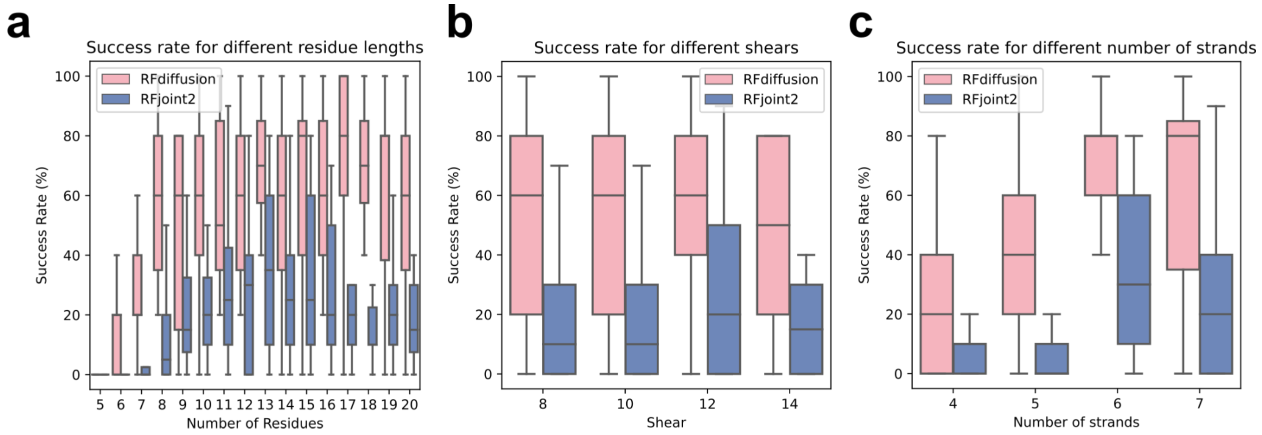


**Figure S4**. RFdiffusion outperforms RFjoint2

**a-c**) Comparison of in silico success rates between RFjoint2 and RFdiffusion, across the parameters tested; number of residues per strand (**a**), barrel shear (**b**) and the number of strands (**c**). Success is defined as AF2 pLDDT > 80, RMSD between design and AF2 prediction < 2 Å, and both the FastRelaxed[11] design and FastRelaxed AF2 prediction having hydrogen-bonding patterns consistent with barrel formation (Methods). RFdiffusion comprehensively outperforms RFjoint2.

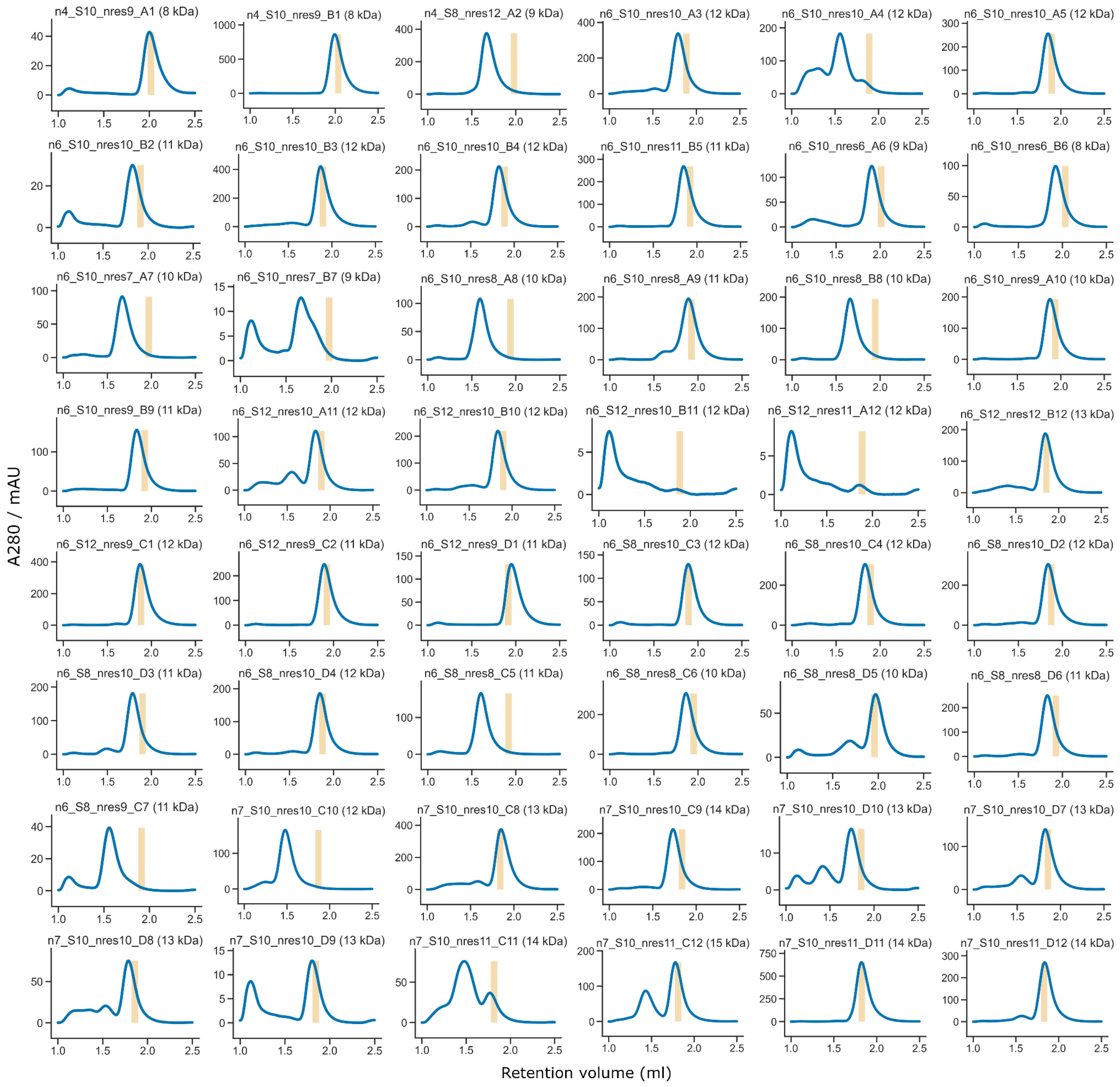
n4_S10_nres9_A1 (8 kDa)
n4_S10_nres9_B1 (8 kDa)
n4_S8_nres12_A2 (9 kDa)
n6_S10_nres10_A3 (12 kDa)
n6_S10_nres10_A4 (12 kDa)
n6_S10_nres10_A5 (12 kDa)
n6_S10_nres10_B2 (11 kDa)
n6_S10_nres10_B3 (12 kDa)
n6_S10_nres10_B4 (12 kDa)
n6_S10_nres11_B5 (11 kDa)
n6_S10_nres6_A6 (9 kDa)
n6_S10_nres6_B6 (8 kDa)
n6_S10_nres7_A7 (10 kDa)
n6_S10_nres7_B7 (9 kDa)
n6_S10_nres8_A8 (10 kDa)
n6_S10_nres8_A9 (11 kDa)
n6_S10_nres8_B8 (10 kDa)
n6_S10_nres9_A10 (10 kDa)
n6_S10_nres9_B9 (11 kDa)
n6_S12_nres10_A11 (12 kDa)
n6_S12_nres10_B10 (12 kDa)
n6_S12_nres10_B11 (12 kDa)
n6_S12_nres11_A12 (12 kDa)
n6_S12_nres12_B12 (13 kDa)
n6_S12_nres9_C1 (12 kDa)
n6_S12_nres9_C2 (11 kDa)
n6_S12_nres9_D1 (11 kDa)
n6_S8_nres10_C3 (12 kDa)
n6_S8_nres10_C4 (12 kDa)
n6_S8_nres10_D2 (12 kDa)
n6_S8_nres10_D3 (11 kDa)
n6_S8_nres10_D4 (12 kDa)
n6_S8_nres8_C5 (11 kDa)
n6_S8_nres8_C6 (10 kDa)
n6_S8_nres8_D5 (10 kDa)
n6_S8_nres8_D6 (11 kDa)
n6_S8_nres9_C7 (11 kDa)
n7_S10_nres10_C10 (12 kDa)
n7_S10_nres10_C8 (13 kDa)
n7_S10_nres10_C9 (14 kDa)
n7_S10_nres10_D10 (13 kDa)
n7_S10_nres10_D7 (13 kDa)
n7_S10_nres10_D8 (13 kDa)
n7_S10_nres10_D9 (13 kDa)
n7_S10_nres11_C11 (14 kDa)
n7_S10_nres11_C12 (15 kDa)
n7_S10_nres11_D11 (14 kDa)
n7_S10_nres11_D12 (14 kDa)
A280 / mAU
Retention volume (ml)

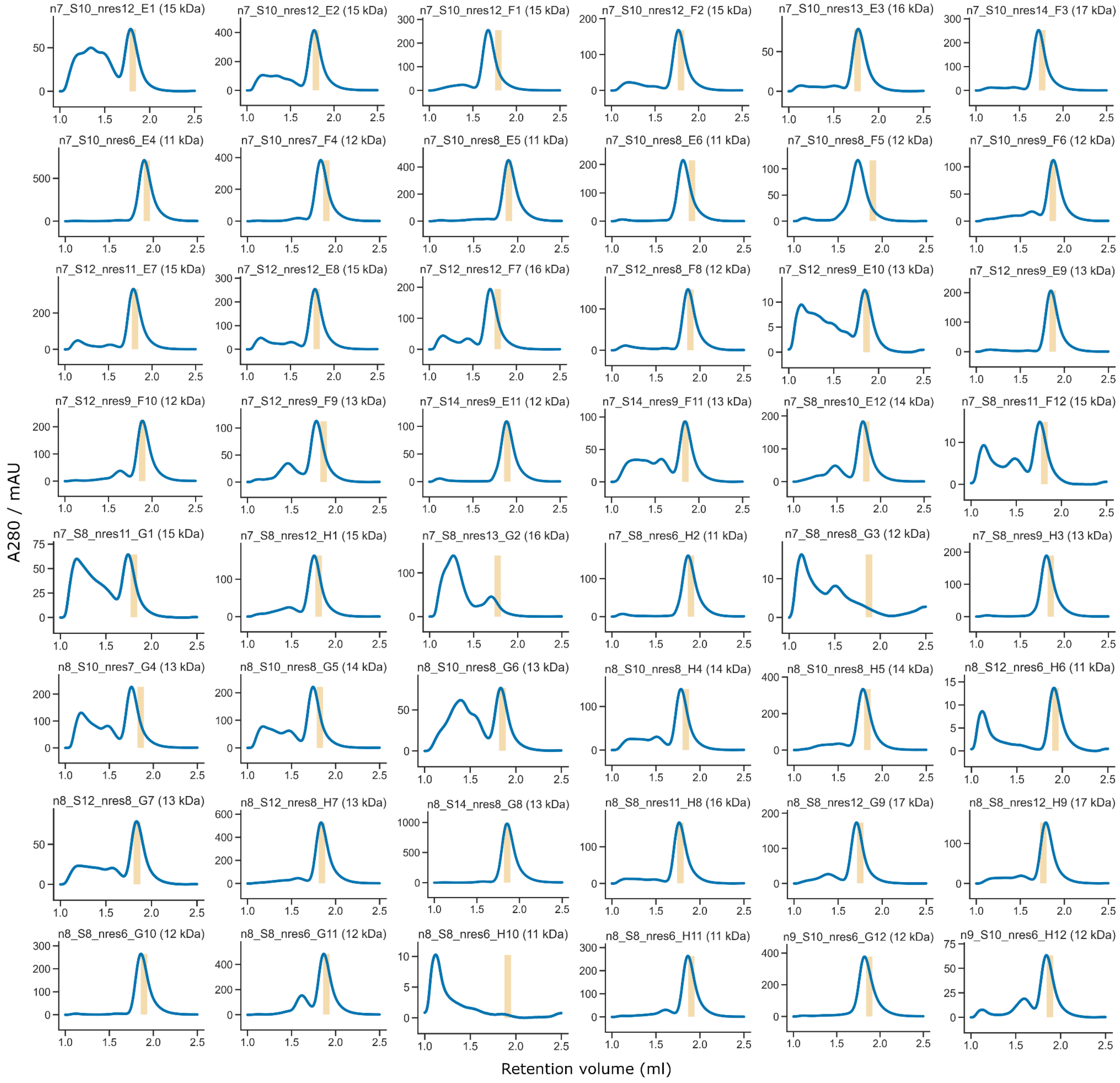


**Figure S5**. Size exclusion chromatography traces with estimated retention volumes (vertical bars) for monomeric designs.

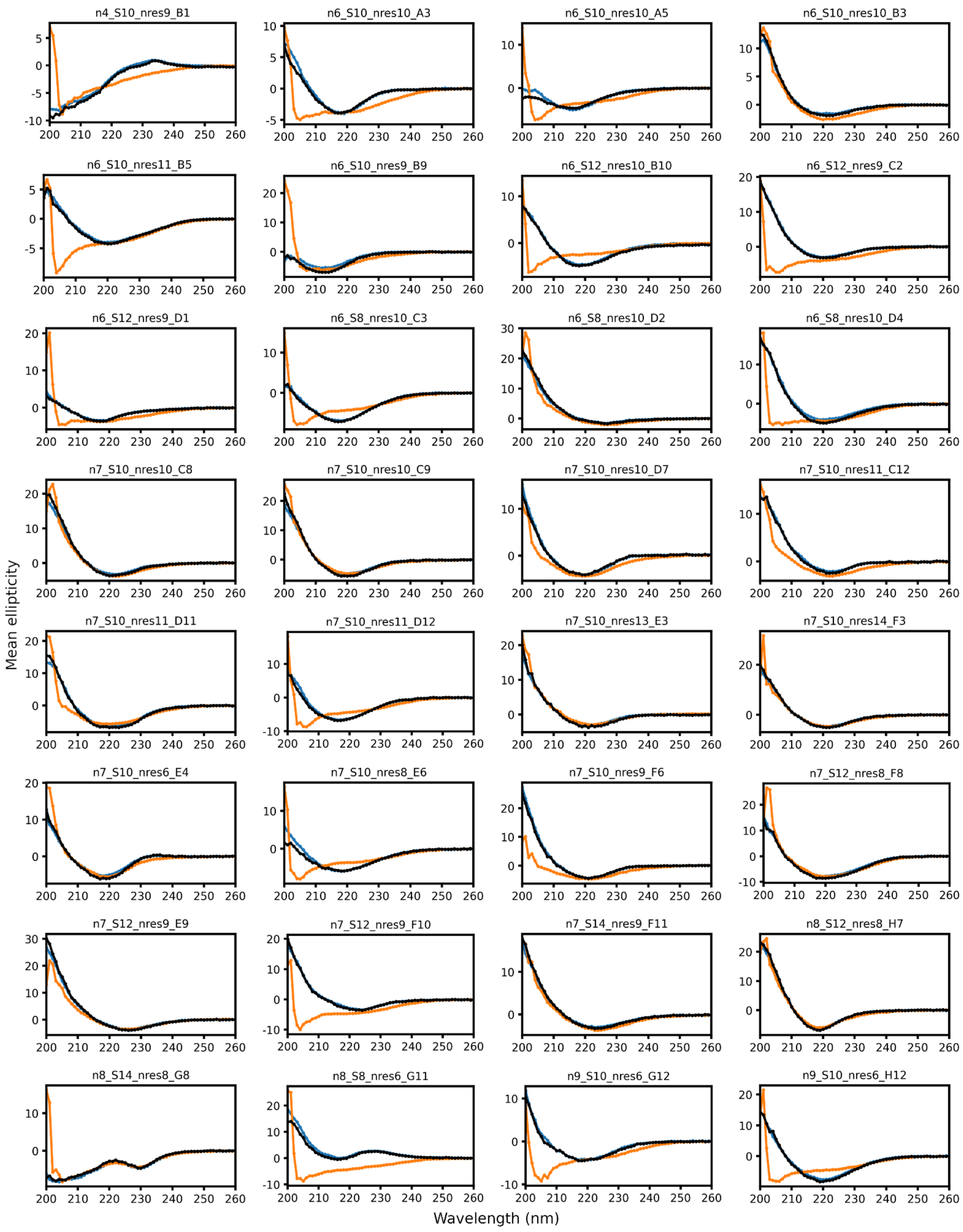


**Figure S6**. Far-UV circular dichroism spectra of designs at 25 °C (blue), 95 °C (red), and back to 25 °C from 95 °C (black).

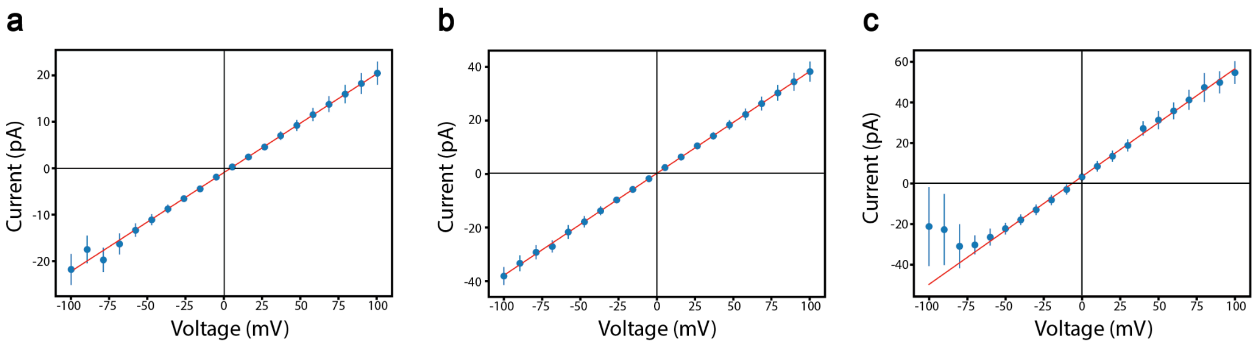


**Figure S7.** Current vs Voltage curves for the lowest current jumps corresponding to single pore insertions of **a.** TMB12_1,**b.** TMB14_1 and **c.** TMB16_1. All measurements were carried out in a symmetric (same on cis and trans side of the bilayers) 500mM potassium chloride solution.

#### **Table S1.** Benchmarking results for RFjoint2

Performance of RFjoint2 on the motif-scaffolding benchmark described in Watson et al.[4] The benchmark seeks to build de novo scaffolds around structural motifs. In RFjoint and RFjoint2, joint sequence-structure design is possible. In RFdiffusion (and RFjoint/RFjoint2 + ProteinMPNN[3]), 8 sequences of the scaffold region (but not the motif) are designed with ProteinMPNN. 100 samples are designed with each method, and the number of in silico successful (AF2 RMSD vs design model < 2 Å ; AF2 motif RMSD vs native motif < 1 Å ; AF2 pAE < 5) is plotted. Full benchmarking details are described in Watson et al.[4] Without ProteinMPNN, RFjoint2 "solves" 15/25 problems (compared to 6/25 in RFjoint), and outperforms RFjoint in 14 cases (with one tie). With ProteinMPNN, RFjoint2 "solves" 21 problems in total (compared to 23 in RFdiffusion and 17 in RFjoint) , and outperforms RFjoint in 15 of these cases. While RFdiffusion still generally outperforms RFjoint2, RFjoint2 outperforms RFdiffusion in 6 cases.

| **Problem Name** | **RFdiffusion (noise=0)** | **RFdiffusion (noise=1)** | **RFjoint** | **RFjoint + Protein-MPNN** | **RFjoint2** | **RFjoint2 + Protein-MPNN** |
|---|---|---|---|---|---|---|
| 1BCF | **100** | 98 | 65 | **100** | 92 | 99 |
| 6E6R_med | **89** | 67 | 0 | 27 | 10 | 38 |
| 2KL8 | 88 | 96 | 71 | 95 | **100** | **100** |
| 6E6R_long | **86** | 63 | 0 | 4 | 24 | 52 |
| 6EXZ_long | **76** | 51 | 0 | 0 | 2 | 8 |
| 1YCR | **74** | 58 | 12 | 57 | 12 | 33 |
| 6VW1 | **69** | 66 | 0 | 24 | 35 | 63 |
| 5TPN | **61** | 59 | 0 | 3 | 28 | 57 |
| 6EXZ_med | **49** | 33 | 0 | 3 | 5 | 14 |
| 4ZYP | 40 | 31 | 1 | 21 | **53** | 49 |
| 6E6R_short | **39** | 29 | 0 | 23 | 7 | 19 |
| 5TRV_long | **37** | 30 | 0 | 0 | 0 | 0 |
| 3IXT | 35 | 16 | 21 | 62 | 77 | **99** |
| 5TRV_med | **24** | 20 | 0 | 3 | 0 | 1 |
| 7MRX_85 | **11** | 6 | 0 | 0 | 0 | 8 |
| 7MRX_128 | **9** | 4 | 0 | 0 | 0 | 4 |
| 1PRW | 8 | 9 | 0 | 22 | 19 | **35** |
| 5TRV_short | 4 | **7** | 0 | 2 | 0 | 1 |
| 7MRX_60 | **2** | 0 | 0 | 0 | 0 | 1 |
| 6EXZ_short | 2 | 4 | 1 | **27** | 2 | 4 |
| 5IUS | 2 | 0 | 0 | 1 | 0 | **12** |
| 5YUI | 0 | 0 | 0 | **1** | 0 | 0 |
| 5WN9 | 0 | 1 | 0 | 0 | 5 | **46** |
| 4JHW | 0 | 0 | 0 | 0 | 0 | 0 |
| 1QJG | 0 | **2** | 0 | 0 | 0 | 0 |

**Table S2**. DNA and amino acid sequences of experimentally characterized designs.

| Design | DNA | AA |
|---|---|---|
| n4_S10_nres9_A1 | CGTCTTAGACGCGACTCCCTTACTTGATTAGCCGCGTCGTTGAAAACatactacggtctcaaggaATGCGCGGCTATCGCGGCGATTTTGTGGATGAAGATGGCCGCGTTATTCCGGGTACCTATTATGGCGGCGAACGTCCGGTACCGGGTGAATGGGTGGAAGTTGTGGCGGAAGACGGCAAAACCATTCGCGCGCGTCTGGTGGGCGAACTGGAAGAATTTGAAGTTCCGggttcccgagaccgtaatgcCGTTAGTTGACATAGTCGCGTGATATGAAACTTCTATGTCGGGATCA | MSGMRGYRGDFVDEDGRVIPGTYYGGERPVPGEWVEVVAEDGKTIRARLVGELEEFEVPGSGSHHWGSTHHHHHH |
| n4_S10_nres9_B1 | AGACGAGACGTCTTGCAAATTATCATCTTACGTACAATCTAAAGTCAGCatactacggtctcaaggaGAAACCGGCTGGAGCGGCAAATGGGTGCTGGATGATGGCTCTGTGGTTGAAGCGACCTATTGGAACAACGGCGTGAAACCGAAGAAAGGCGAGAAATATGAAGTCGTGCTGCCGGATGGCACCGTGGCGACCGGTGAACCAGTTGGTGAACTGACCAAAGTGACCggttcccgagaccgtaatgcTGTCGACATTGACCGCTTTAATTTTCGCGCCGTACATACCTGAACCGG | MSGETGWSGKWVLDDGSVVEATYWNNGVKPKKGEKYEVVLPDGTVATGEPVGELTKVTGSGSHHWGSTHHHHHH |
| n4_S8_nres12_A2 | TACACCTGACGCTATGGACAACGTATatactacggtctcaaggaGCGAAATATACCGGCAAAGGTAAAGGCACCATGACCAACGCGGATGGCAAAACCGTGGATGTGGAATTCACCGAATTTGAAGGTACCGGTACCCTGGAAGATGGCGTTCTGACCCTGACCAGCTGGAAAGGCAAAGGCGTTGCGGACGGCAAACCGTTCGAAGGCAGCGGCTCTGGTACCGTTACCGAAGGCGAAGTGAAAGCGGTGGAAggttcccgagaccgtaatgcCGGGGGTGAGGCTTGGGAGTGGCCTG | MSGAKYTGKGKGTMTNADGKTVDVEFTEFEGTGTLEDGVLTLTSWKGKGVADGKPFEGSGSGTVTEGEVKAVEGSGSHHWGSTHHHHHH |
| n6_S10_nres10_B2 | atactacggtctcaaggaGCGCCGAAAGTGATGAGCGGCACTCTGACCGTTACTGGCGGTACTGGCATTAGCGGTGGCACCGTGGATGTGACCGGCGTGGTTGATGATGGCGTGTTCACCGGTACGGGCACCTTTACGGGTACGGTTAACGGTAAAGCGGCGAGCGGTCCGGTGAACGTGGAAGCGGAAGTCGATGAAAACGGCGAAGTTACCGGTGGTACCCTGAGCGGTACCGTGACGGGCGGTTATGTGATTAACGGCGCGGTTAAACTGACCGGCACCCTGACCGAAGTGCCGGGCACCggttcccgagaccgtaatgc | MSGAPKVMSGTLTVTGGTGISGGTVDVTGVVDDGVFTGTGTFTGTVNGKAASGPVNVEAEVDENGEVTGGTLSGTVTGGYVINGAVKLTGTLTEVPGTGSGSHHWGSTHHHHHH |

| | | |
|---|---|---|
| n6_S10_nres10_A3 | atactacggtctcaaggaGCGAAACTGTATAAAGGCAAACTGAAAGTCACCGGCACCATCGGCGATACCAAAGTAGAAGGTGAAGTCGAAGCGTGGGGCGTGATTGATGGCGACACCACCGAATTTACCCTGAATGTTAAAGGCGAAGTGGAAGGTGGCACCATTGAAGGCGGCACCGTTAAAGCAACCGGTGAACTGACCCCGGAAGGCCTGAACGCGACCGGTACGTTTACGGGTACCTTTAACGGCGAAGAGAAAACGGGCGAAGTGACCCTGGAAGGTAAACTGGAAGAAGTTGAACCGggttcccgagaccgtaatgc | MSGAKLYKGKLKVTGTIGDTKVEGEVEAWGVIDGDTTEFTLNVKGEVEGGTIEGGTVKATGELTPEGLNATGTFTGTFNGEEKTGEVTLEGKLEEVEPGSGSHHWGSTHHHHHH |
| n6_S10_nres10_B3 | atactacggtctcaaggaAAACTGAAAGTGGGTGAAGGCACCGGCGATGGCGAAGTGGGCGGCGTTAAAGTGAAAGATGTCAAAGTTCTGGTGGTGGGCAAAATTGAAGATGGCAAATTCGAAGGTGAAGCGACCATTACCGGCCTGCCGAACGGCAAAACCGCGAAAGGCAAAGCGGAAGGTAAAGTCGAAGGCAACAAAATTAAAGGCACGGTGACCGATATTGAACTGGATGGTAAGAAAGTTGGCGGTAAAATCGATTTTGAAGGCGAATACTGGGACATTGAAGAAggttcccgagaccgtaatgc | MSGKLKVGEGTGDGEVGGVKVKDVKVLVVGKIEDGKFEGEATITGLPNGKTAKGKAEGKVEGNKIKGTVTDIELDGKKVGGKIDFEGEYWDIEEGSGSHHWGSTHHHHHH |
| n6_S10_nres10_A4 | atactacggtctcaaggaGAAAGCCTGAAATTTGAGGGCGAACTGGAACTGAAAGGTAACGTCGGTGGCAAAGAAGTTAAAGGTACGGTGAAGTTTGAAGGCGAACGTAAAGGCAACTACGTTGAGGGTGAATTCGAAGGCGAGCTGGAAGGCGGCATTAAAAGCGGCAAACTGGAAGGTGAAGGGGAGTTCGAGGGCGATTATTTCAAAGGCACCCTGACCGGCAACTTTGATGGTAAAGAAGGTACCGTCGAAGCGAAAGGCGAATTAAAATTCGAGGAACCGggttcccgagaccgtaatgc | MSGESLKFEGELELKGNVGGKEVKGTVKFEGERKGNYVEGEFEGELEGGIKSGKLEGEGEFEGDYFKGTLTGNFDGKEGTVEAKGELKFEEPGSGSHHWGSTHHHHHH |
| n6_S10_nres10_B4 | atactacggtctcaaggaATGAAACTGTATAAAGCGCTGATTGAAGATCTGAAAGGTGAAGTCGGTGGCAAGAAAGTGGAAGGCGGCAAAGCGGAACTGATTGGCAACCTGGAAGATGGCAAATTTAAAGGTGAGGGCGAAGCCGAAGTGACCGTTGAAGGTAAGAAAGTCAAAGGCCTGGCGGTTGGCGAAGGCAAAGTCGAAGGCGAAAAGATTAAAGGCGAACTGAAATTTGAAAGCGATGACGGCGAGAAAGGCGTGCTGAAAGGGGAAGGCAAGGTCGAAGAAATTGAAggttcccgagaccgtaatgc | MSGMKLYKALIEDLKGEVGGKKVEGGKAELIGNLEDGKFKGEGEAEVTVEGKKVKGLAVGEGKVEGEKIKGELKFESDDGEKGVLKGEGKVEEIEGSGSHHWGSTHHHHHH |
| n6_S10_nres10_A5 | atactacggtctcaaggaGAAGAAATTTATAAAGTTAAAATCACCGATCTCAAAGGTGATGTGGGCGGCAAAGTGGAAGGCGGTAACGCGGTGCTGGAAGGTGTGGTTAAAGATGGCATTTTCAGTGGTGAAGGTGAAGTCGAAGTGACCGTGAACGGCAAGAAGAAAACCGGTAAAGCGAAAGGCGAAGGCAAGGTGGATGGCGATGAATTTAGCGGCGAAATTGATGTTGAAGATGATGA | MSGEEIYKVKITDLKGDVGGKVEGGNAVLEGVVKDGIFSGEGEVEVTVNGKKKTGKAKGEGKVDGDEFSGEIDVEDDDGNKGKVKGKGKVFKL |

| | | |
|---|---|---|
| | CGGCAACAAAGGGAAAGTGAAAGGCAAAGGTAA AGTGTTTAAACTGGATggttcccgagaccgtaatgc | DGSGSHHWGSTH HHHHH |
| n6_S10_nres11_B5 | atactacggtctcaaggaGCACCAGTTGCGGTGAGCGG CACTGCGACCTTTACTAGCGGTACCCTGGGTGG CAAAGCGGTTGCAGGCGGCACCTTAACCGATATT ACGGGTACCGTGACCGGCACGACTTTTACGGGC ACCGCGACCTATACTGCCACCGATGGCACCACC GGTACCATTACCGGTTTAACCGCGGCCGTTGCGG ATGGTGCAATTACCGGCACAGCGACTGGTACCGA TACCGCAACCGGCCAGGCGGTTACGTTAACCGG TGTGAGTGGTACTCTGACCGCGGTTCCGGCGCC Gggttcccgagaccgtaatgc | MSGAPVAVSGTATF TSGTLGGKAVAGG TLTDITGTVTGTTFT GTATYTATDGTTGTI TGLTAAVADGAITGT ATGTDTATGQAVTL TGVSGTLTAVPAPG SGSHHWGSTHHH HHH |
| n6_S10_nres6_A6 | TCGACGTCAAGTATAATCACCGCGAGGTCGGACC Tatactacggtctcaaggagcgctgaccatcaccggcaa AGTGACCATTAACGGCAAGGATGCGGGCACCGT GACCGGCGAAGGCGTTGAAGGCGAAGAATTTAC GGGCAAAGATGAAAACGGCGTGGAATTTAAAGG CGTGTATAAAGATGGCGTGCTGACCGGTACCCTG AATGGTGAGAAATTTAGCGGTCCGGCCACCGCG ACCGCGggttcccgagaccgtaatgcAGTCATAAGCGTTG GGGACTGTTTCGCGCCTTCAA | MSGALTITGKVTIN GKDAGTVTGEGVE GEEFTGKDENGVE FKGVYKDGVLTGTL NGEKFSGPATATAG SGSHHWGSTHHH HHH |
| n6_S10_nres6_B6 | GCCCTACACCCCCTTACTTTGGAGTCGGGCAGA GatactacggtctcaaggaGCAACTGTGACCGGTACTGG TGGTACTCTGGGTGGTGCGCCGATTGCGGGCTT TAGCGGTACCTTGACCGATGGCGTGTTTACCGGC ACTGTGGATGGCAAACCGGCAACGGGTACTGTG GTGGACGGTGTTGCGAACTTTACCGTTGATGGTG TGGCGGGTACCGCGCCGGTGACTTCTATTACCCC AGCGCCGggttcccgagaccgtaatgcTGTAAGTCAATTC GTATCGACGCATAATCTACT | MSGATVTGTGGTL GGAPIAGFSGTLTD GVFTGTVDGKPAT GTVVDGVANFTVD GVAGTAPVTSITPA PGSGSHHWGSTH HHHHH |
| n6_S10_nres7_A7 | CTTCACTGTCGAAGTATTTACAGTCatactacggtctca aggaGGCAAAGTGACCATTAAAGCGGAAGATGGC ACCGTTATTACCGGTGAAGCGAAAGCGATTGAAG AAAACGGCGTTACCCTGATCGAAGGCGAATTCGA AGGTGGTAAAGCGAGCGGCAAAGCCGAAAACCT GAAGAATGGCAAAACCGGCACGGGCACCATCAC CGTGAACGGTAAAGATTATACCGGCGAAATTTACT ATGAAGAAATTGAAggttcccgagaccgtaatgcGGGTTT AGTTCAACCCCCGTAGCA | MSGGKVTIKAEDG TVITGEAKAIEENG VTLIEGEFEGGKAS GKAENLKNGKTGT GTITVNGKDYTGEI YYEEIEGSGSHHW GSTHHHHHH |
| n6_S10_nres7_B7 | CCATGAAAAGGCGGTAGCGAatactacggtctcaaggaG GCAAAGTGGTTGCAACCTATACCGGCCCGCCGA GTGGTGAAGGTACCGTGACCGGTGAAATTGTGG GCGATGGCTTTGTTGGCACTGATGCGGCAACCG GTGCAACTGTGCGCGGCCCGTTAGTTCCAGTTG GCGAAACCGGCGAAGGCGAATACGAAGATGATG | MSGGKVVATYTGP PSGEGTVTGEIVGD GFVGTDAATGATV RGPLVPVGETGEG EYEDDGVWVSGFT YTPTEIEPAVAGSG |

| | | |
|---|---|---|
| | GCGTTTGGGTGAGCGGCTTTACGTATACCCCGACCGAAATTGAACCGGCCGTGGCGggttcccgagaccgtaatgcTTGCACTAAGTGCCGGGGGC | SHHWGSTHHHHHH |
| n6_S10_nres8_A8 | CTTGAGTTTGTCTCatactacggtctcaaggaATGCGCGTGTTAGAAGTGGATAGAGGCGAAGTGGGCGGCGAAGAAGTTGAAGGCGGCCGCGTAGAAGGTGATGTCGAAGAGGGTGAAACTGGTCCAGCGCGCGGTGAAGTTGGCGGTGTTCCAGCCGAAGGCGAATTATATGTTGAAGATGGCCGAGGTCGTGTGGAAGGCGAGGCGGATGTTGATGGTGAACGTGTGCCGTTTCGTGCGGAAGGCCGTGTCGTTGAACCGCCGggttcccgagaccgtaatgcGTGGCATCTTATCG | MSGMRVLEVDRGEVGGEEVEGGRVEGDVEEGETGPARGEVGGVPAEGELYVEDGRGRVEGEADVDGERVPFRAEGRVVEPPGSGSHHWGSTHHHHHH |
| n6_S10_nres8_B8 | AGGCCATGGGATTGCCAGAatactacggtctcaaggaACCGTGTATGAATTTGAAGTTGAAGGTGGCACCAACGGCCTGACCGGTGGCAAATTTGAGGGCGAACTGGAAGGCGATAAAGTTAAACTGAAAGGTAAAGACGATAACGGCGTTGAAGTCGAAGGTGAAGGCACCCTGAAAGATGGCAAAGCGAAAGTGGAAGGCAAAGACGAAAACGGCAAACCGGTTGAATTTACCGCGAAATTTGTGAAAGAGCTGGAAAAGggttcccgagaccgtaatgcAAGGCTGTTCTATATTCA | MSGTVYEFEVEGGTNGLTGGKFEGELEGDKVKLKGKDDNGVEVEGEGTLKDGKAKVEGKDENGKPVEFTAKFVKELEKGSGSHHWGSTHHHHHH |
| n6_S10_nres8_A9 | AACAGCTCatactacggtctcaaggaGGCATGGATCTGGTGGAAGGTGAAGTCGAAGGCACCGTGGCGGGCAAACCGGCAAAAGGCGAATTTAAAGGTGAACTGAAAGACGGCAAAGTCAAGGGCGAAGTTAAAGTTGAAGTGGATGGCAAAGAAGTGAAAGGCCTGGCGGAAGGTAAACTGTATGAAGGCAACTACGTTAAATTAGAAGGCAAAACCGATGATGGCACCGAAGGCCTGAAAGCCGAGGGCAAAGTTAAGGTGATTAAACCGggttcccgagaccgtaatgcTAACCGAC | MSGGMDLVEGEVEGTVAGKPAKGEFKGELKDGKVKGEVKVEVDGKEVKGLAEGKLYEGNYVKLEGKTDDGTEGLKAEGKVKVIKPGSGSHHWGSTHHHHHH |
| n6_S10_nres9_B9 | GGCTTCGatactacggtctcaaggaGCGACCAAAGTGACCATTGAAGGTACCCTGGAAACCGATGACGGCTATGTGGGCGAAGTGAGCGGCGAAGGCGTGTTATTAGAAGGCGGCAAAGGCGTTTTCGATCTGACCGATACCGAAACCGGCGAGAAATTTACCGTCGAACTGAAGAACGCGAAACTGGGCGAGGGCACCTTCGAAGTTACCAAAATTATTCGCGATGGCAAAGATATTGGCGGCGGCACGTTTACCGGCACCATTACCGAAGTGggttcccgagaccgtaatgcGTAAGT | MSGATKVTIEGTLETDDGYVGEVSGEGVLLEGGKGVFDLTDTETGEKFTVELKNAKLGEGTFEVTKIIRDGKDIGGGTFTGTITEVGSGSHHWGSTHHHHHH |
| n6_S10_nres9_A10 | AACGCACACTatactacggtctcaaggaAAATATCTGCTGGAAGGCGAATTTACCGATGCCGATGGTACCAAAGGCACCATTAAAGCGGAAGGTGAAGTTAAAGATGGCGTGTTTACCGGCGACGGCACCGCGACCGTGGATGGCAAGAAATATAAAGTCACCGGCGTCAAAGGCGAAGTGGACGGCGATAAATTTAAAGGCAAAACCG | MSGKYLLEGEFTDADGTKGTIKAEGEVKDGVFTGDGTATVDGKKYKVTGVKGEVDGDKFKGKTGDFSDGTAGGAPVEGT |

| | | |
|---|---|---|
| | GTGATTTTAGCGATGGCACTGCGGGCGGTGCGC CGGTTGAAGGTACCGTGAAGAAAGTGGAAggttccc gagaccgtaatgcCCTTGAGAT | VKKVEGSGSHHW GSTHHHHHH |
| n6_S12_nres10_B10 | atactacggtctcaaggaGTGGAGAAATTTAAAGTCGTG GGCAAAGTGATTAGCGGTACCGTGGGTGGCGAA GCGGTGGAAGGTGGTACCATTGAAGGCGAAGCC ACCAAAGAAGGCGACGAAACCACCATTACCGGC AGCTTTACGGGCACCGTTGGCGGTGAAAACGTG AGCGGCAGCAACCTGACGTTTAAAGGCAAACTG GATGGCAACAAATTTACCGGTACCCTGACCGGCA CCATTGGCGGCAAAACCTATACCGGCGGCGAACT GACCTTTGAAGGTGAACTGATTGAACCGggttcccga gaccgtaatgc | MSGVEKFKVVGKVI SGTVGGEAVEGGTI EGEATKEGDETTIT GSFTGTVGGENVS GSNLTFKGKLDGN KFTGTLTGTIGGKT YTGGELTFEGELIE PGSGSHHWGSTH HHHHH |
| n6_S12_nres10_A11 | atactacggtctcaaggaATGAAAACCTTCAGCGGCACC TTAACCTTTACCGGCACCCTGGGCGGTCAGGCG TTAACCGGTGCAACCGCAGAAGTTTCTGGCGTG CTGGAAGGCGATACCCTGACCGGTGAAGGCACT CTGACCGGTACCATTGGCGGCAAAGCGGTTAGC GGCCCGTTTACCTTTACGGGCAAATATACCCCGG GCCAGGATTTTCAGGGCACGCTGACCGTGCAGG TGGGCGATAAAACCGTGAGCGGCGATGTGACCT TTAGCGGTGAACTGGAATTTACCGATCTGggttcccg agaccgtaatgc | MSGMKTFSGTLTFT GTLGGQALTGATAE VSGVLEGDTLTGE GTLTGTIGGKAVSG PFTFTGKYTPGQD FQGTLTVQVGDKT VSGDVTFSGELEFT DLGSGSHHWGSTH HHHHH |
| n6_S12_nres10_B11 | atactacggtctcaaggaCGCACCCGCGTGGTGGCGCG CGTGCGTCTGACCATCGATGGTAAAGAACATCGC GGCGAACTGACCTTTGATGGTGAAGTGACCGAA GGCGAAGGCTTTCGCGGCACCTTTAGCGGCACC ATTGATGGCAAACCGGCGAGTGGTGAACTGGAA GGCCGTGGCGAAATTACCGAAGAACGCTTCCGC GGTGAAGCGACCGGCACGATTAACGGCCAGAAA ATTGACGGCGCGCCGATTGATGCGGAAATTGTGG AACGCGAAGAAATTCCGCCGggttcccgagaccgtaatg c | MSGRTRVVARVRL TIDGKEHRGELTFD GEVTEGEGFRGTF SGTIDGKPASGELE GRGEITEERFRGEA TGTINGQKIDGAPID AEIVEREEIPPGSG SHHWGSTHHHHH H |
| n6_S12_nres11_A12 | atactacggtctcaaggaGCGCGCCGTGTTCGCGCGAC TGGCACCGTTGAAGTTGATTTGCCAGGTGGCCG TCGTGCGACCATCGAATTTGATGGTGAAGGCGAA CTGGAAGGTGATACCCTGGTGGTGCCGTTCACC AGCGGCACCATTGATGGCAAACCGTTTACCGGC GAAGGCACCTTAACCATTCGCGGTGATCCTGAAG CGCCGGGTGCACCGGTTACTGCAACTTTCACCG GTGATGCGGGTACCTTTGTGGATTATCCAGGTCG CGTGGTTGCGGCGCGTGTGACTCCGTTACCGGC Gggttcccgagaccgtaatgc | MSGARRVRATGTV EVDLPGGRRATIEF DGEGELEGDTLVV PFTSGTIDGKPFTG EGTLTIRGDPEAPG APVTATFTGDAGTF VDYPGRVVAARVT PLPAGSGSHHWGS THHHHHH |
| n6_S12_nres12_B12 | atactacggtctcaaggaATGAAAGGCTTTGTCGCCGAA GGCGAAATCACCAACGGCAGCTTTAACGGCAAA | MSGMKGFVAEGEI TNGSFNGKPVKGK |

| | | |
|---|---|---|
| | CCGGTGAAAGGCAAATTTAAAATGAAAGTGATTG GCGAACTGGGCAAAGGGAAAAGCGCGCCGGCG GAATTTGATATTATTAACGAAAACGGCGAAGTTGT GGGCCGCTTTAAAGGCACCATTTATTTTGGCGATA TTGATGGCAACAAATTCACCGGCAAATTCGATGC GGAAGGTACCCTGGGCGATGAGAAAGGTAAATTT AGCGGCGAATTCGAAGCGGAAATTACCGAAACCT TTGAACTGggttcccgagaccgtaatgc | FKMKVIGELGKGKS APAEFDIINENGEVV GRFKGTIYFGDIDG NKFTGKFDAEGTL GDEKGKFSGEFEA EITETFELGSGSHH WGSTHHHHHH |
| n6_S12_nres9_C1 | atactacggtctcaaggaGGCAACTATGTGAAATTCAAAG TGTATGATGAAAACGGCAACCTGCTGGGCGAAAT TTACATCGAAACCGATGAGAACTTCGGCAAGAAA GGCTATAAAAGCAAAATTAGCGGCGTGATTAATGG CAAACCGATTGATGAAGCGGAAGTGACCTTTCTG ACCGATATCGAAAATGGTAAAGCGAAAGTGATTAT CAGCGGTGTGATTGGCGGCGAGAAAATTGAAAA CAAAGTGGTCAACATCGAATTTATCGAGAACGTG GACGAACCGggttcccgagaccgtaatgc | MSGGNYVKFKVYD ENGNLLGEIYIETDE NFGKKGYKSKISGV INGKPIDEAEVTFLT DIENGKAKVIISGVI GGEKIENKVVNIEFI ENVDEPGSGSHH WGSTHHHHHH |
| n6_S12_nres9_D1 | TTTAGatactacggtctcaaggaATGACTGTGGAAGTGG CGACTGTTACCGGCGAAGGTGGTGCGGTTGCAC CGGGTTCTACTTTGACCGGCACTCTGGAACCGG GTCAAACCGTGGATTTTAGCGGCGTGTTTCGTGG CCGTCCGGCGACCGGTCGTTTAGAACTGCTGAC TCGCCCTGAAGTTGGCAAATGGTTTGATGCGCGT GGCACCATTGGCGGCGTTGAAGTTCCGCGTCTG CGTCTGTTCGTGGAAGAAATTCGCACCGTCGAAG TGggttcccgagaccgtaatgcGGTTC | MSGMTVEVATVTG EGGAVAPGSTLTGT LEPGQTVDFSGVF RGRPATGRLELLTR PEVGKWFDARGTI GGVEVPRLRLFVE EIRTVEVGSGSHH WGSTHHHHHH |
| n6_S12_nres9_C2 | GatactacggtctcaaggaAAGAAATATAAAGTTACCGCG AGCGGCACGGCGAACGGCCAGGCGTTTGATGAA GTTGTGTTTACCGTGCCGAGCCTGAAGAAAGGT GCAGTAGCGCCGGTTACCGGTACCGTTGGCGGT GTGGCAATTGCGTCTGGTACCGTGACCTTTCTGG ATGATGTGGAAGTGGGCAAAACGTTTACCTTTAC CGGCACCCTGAACGGCGTTGCGATTACGGGTGG CACCGCAACCGTGAAAAGCATCGAAGAAATTGAA GCGGAAggttcccgagaccgtaatgc | MSGKKYKVTASGT ANGQAFDEVVFTV PSLKKGAVAPVTGT VGGVAIASGTVTFL DDVEVGKTFTFTGT LNGVAITGGTATVK SIEEIEAEGSGSHH WGSTHHHHHH |
| n6_S8_nres10_D2 (BBn6 in main text) | atactacggtctcaaggaAAGAAAGTGAAATTCGTGGCA GAAGGCACCTTTAAAGCGACCAACAAACCGATTG AAGGCCTGGTTAAAGCCGAAGGCGAGGCGGAAC TGGGCAAAGATTTTGAAGCAGAAGTGACCGGCG AACTGGATGGCAAACCGTTTAAAGGCAAAGCGAA ATTTCCGCCGTACAAAGGTAAAGGCTATACCTATC GCGGCGAAGCGGAAGGCGAAATCGATGGCGAAC CGATTAAAGGTATTGTGGATGCGACCGTGTTCAA AGAAGAGGAAGTTCCGGAAGGTGAAggttcccgagac cgtaatgc | MSGKKVKFVAEGT FKATNKPIEGLVKA EGEAELGKDFEAE VTGELDGKPFKGK AKFPPYKGKGYTY RGEAEGEIDGEPIK GIVDATVFKEEEVP EGEGSGSHHWGS THHHHHH |

| | | |
|---|---|---|
| n6_S8_nres10_C3 | atactacggtctcaaggaCGCGTGCTGGTTAGCGGTGAAGGGGAATTCGAAGGCACCCTGGGCGGCAAACCGGCCAAAGGTACCGTTCGTGTGGAAGATGCCGAGGTTGAAGGTGATCGCGCACGTGGCCCGGCAACAGTTACCGCAGATGGCAAAACCGGCGAAGTGGAAGCGGATCTGGAATTACGCGGCGATGAACTGCGCGGCCGTGTTCGTGGCGAAATTGAGGGTGAACCGGTAGAAGGCGAAGCGCGTGTGCGTGTTACCGAACGTCGCGAACTGCCGCCGggttcccgagaccgtaatgc | MSGRVLVSGEGEFEGTLGGKPAKGTVRVEDAEVEGDRARGPATVTADGKTGEVEADLELRGDELRGRVRGEIEGEPVEGEARVRVTERRELPPGSGSHHWGSTHHHHHH |
| n6_S8_nres10_D3 | atactacggtctcaaggaAAAGAAATTATCGTTAACGGCAAAGGCACCGCCGAAGGCAAACTGGGTGGCAAAGAAGTGACCGGAGAAGCGGTGCTGGAAAACGGCCGCCTGGAAGGCGATAAAATTACCGGCAAAATTCTGGTGGAAATGGATGGCCAGAAAGGCGAAGTCGAAGCGGAACTGACCCTGGAAGGTGATAAAGCGACCGGCCCGGTTAAAGGTGAAGTGGGCGGCAAACCGTTTGAAGGCGAGGCGACCATTGAAGATCTGGAACTGGAAGAAGTGGAAggttcccgagaccgtaatgc | MSGKEIIVNGKGTAEGKLGGKEVTGEAVLENGRLEGDKITGKILVEMDGQKGEVEAELTLEGDKATGPVKGEVGGKPFEGEATIEDLELEEVEGSGSHHWGSTHHHHHH |
| n6_S8_nres10_C4 | atactacggtctcaaggaGAAGAAAAGCTGTTTAAATTTGAAGGTACCTTTACCGGCGACGGCGGCAAACTGAAAGGTAAAGTTAAAGGCGAAGGCTATGCGAAAGAAGGCGATACCAGCTTCGAAGGTGAAGCGAAAATTTATCCGGATGATGGCGGCCCGGTGATTAGCGGCAAAGCCAAAGTTGAGGGCGAAATTAAAGATGGCTTTAAAGGTGGCACCGGCACCTTTGAAAACGAAGACGGCGTGAAAGGCACCATTACCGATTTAAAAGGCAACCTGGAAGATATTGAAggttcccgagaccgtaatgc | MSGEEKLFKFEGTFTGDGGKLKGKVKGEGYAKEGDTSFEGEAKIYPDDGGPVISGKAKVEGEIKDGFKGGTGTFENEDGVKGTITDLKGNLEDIEGSGSHHWGSTHHHHHH |
| n6_S8_nres10_D4 | atactacggtctcaaggaGGCCGCCTGCGTGGCGAGGGCGAATTCGAAGGTGAAATCGGCGGCGAAAAGGCGAGCGGTCCGGTTACCGTTGAAGGTGAAGCGGAACTGCGCGATGGTGAATTACGCGCGCGTGTTGAAATTAAAGGCGAAGCGAACGGCGGTGAGAAAATTGAAGGAGAAGGCGTTTTACGCGGTCGCCTGGAAGGTGATACCGTTCGCGGTGAACTGGAAGGCGAACTGAACGGCAAACCGTTTCGCGGCGAAGTGGTGTTACGCGATGCGCGCTTTGAAGAACTGCCGGAAggttcccgagaccgtaatgc | MSGGRLRGEGEFEGEIGGEKASGPVTVEGEAELRDGELRARVEIKGEANGGEKIEGEGVLRGRLEGDTVRGELEGELNGKPFRGEVVLRDARFEELPEGSGSHHWGSTHHHHHH |
| n6_S8_nres8_C5 | GGGTGTGatactacggtctcaaggaGAGAAACGCGTTTTCCGTGGCGAGGGTGAAGGCGAACTGGAAGGAGAACGCGTCGGCTTTCGTGCAGAATTAGCGCTGCCAGGTGTTGGTCAGGTGGGCCGCGGCGAAGTTGAAGATGAAGAAGGCCGTCGCGGTGAAGCAGAAGGCGTGCTGTTAGATGAGGATGGCAGCCGTATTCTGGTGCGTGGTGAAGTGGGCGGTGTGCCGTTTGAAGGTGAACTGTTTGGCACCGAAGATATTGGTAC | MSGEKRVFRGEGEGELEGERVGFRAELALPGVGQVGRGEVEDEEGRRGEAEGVLLDEDGSRILVRGEVGGVPFEGELFGTEDIGTVGSGSHHWGSTHHHHHH |

| | | |
|---|---|---|
| | CGTGggttcccgagaccgtaatgcACATAC | |
| n6_S8_nres8_D5 | CCGGACGTTGTATACGatactacggtctcaaggaATTGTTGGCACCGGCACCCTGACCACCAACCTGGGCACCCATACCGGCAATATTACCGTGACCGGTACCGATAAGAACAACATGACCATTAGCGGTACCCTGGATGGCCAGGATTTCTCTGGCACGGGCACCGTGAACGATGATCTGAGCAGCGGCACCTATAGCGGCACGCTGAACGGCCAGTCTGTTAGTGGCACCTTTAAACTGACCAACATTAACAAAATTCCGGGCggttcccgagaccgtaatgcGTGCGTTCAGAAGAC | MSGIVGTGTLTTNLGTHTGNITVTGTDKNNMTISGTLDGQDFSGTGTVNDDLSSGTYSGTLNGQSVSGTFKLTNINKIPGGSGSHHWGSTHHHHHH |
| n6_S8_nres8_C6 | CTAGACACTTTAGCTGCatactacggtctcaaggaATGGCGGTGTTTAAAGGCACCGTGGAACTGGATGGTGGTGAAGTGGGCACCTTTGAAGCGGAAGGCAAACCGGGCGAAACCGTGCCGGTCAAAGGCGAAGTGGATGGCGAACCGATTGAAGGCGAGGGCGTTATTAGCGAAGATGGCAAAAGCGGTACCTTCAAAGGTGAATATCGCGGTAAACCGGTGACCGGCAAACTGAAAGTGACCGAAGTTGAAGTGCCGCCGggttcccgagaccgtaatgcTTTTAAGTTGTGAAAAA | MSGMAVFKGTVELDGGEVGTFEAEGKPGETVPVKGEVDGEPIEGEGVISEDGKSGTFKGEYRGKPVTGKLKVTEVEVPPGSGSHHWGSTHHHHHH |
| n6_S8_nres8_D6 | CGCAATCGatactacggtctcaaggaGAAACCCTGCTGCAGCTGACCGGTGAAGGCGATTTCGATGGCGTGAAATTTACCTTTGTGACCGAACCGTTTGCGCCGGATGCGACCACCGTGAACTTTACGGGCACCACGGAAGATGGCACCGTCTTTACCGGCACCGCGACCTTAAATCCGGATCGTACCGGTGGCACTGGTGAAGCGACCATTAACGGCGAGAAAGTGACCGGCGAATTTAAAGTTACCGAAGTGCGCGAAGTTCCGGCGCCGggttcccgagaccgtaatgcTTTCGCTG | MSGETLLQLTGEGDFDGVKFTFVTEPFAPDATTVNFTGTTEDGTVFTGTATLNPDRTGGTGEATINGEKVTGEFKVTEVREVPAPGSGSHHWGSTHHHHHH |
| n6_S8_nres9_C7 | atactacggtctcaaggaGCGGTGGTGGAAGATTTTAAAGCGGAAGGCGAACTGGACGGCAAACCGGTGACCGCGGATTTTGGCACGGTGGATTATGAAGAAGGCAAAGGCGTTAGCGGTGAAGGCACCGTGACCATTGATGGCAAGAAAGTGCCGGGCAGCTTTGATGGTGATTTTGATGCGGAAACCCAGAAAGCCAAAGGTACCATGAAAGCGACCCTGGATGGTAAAGAACTGAGCGGCGAAGCCACCGGCAGCGGCAAAGTTGTGGAACTGGAAGGCggttcccgagaccgtaatgc | MSGAVVEDFKAEGELDGKPVTADFGTVDYEEGKGVSGEGTVTIDGKKVPGSFDGDFDAETQKAKGTMKATLDGKELSGEATGSGKVVELEGGSGSHHWGSTHHHHHH |
| n7_S10_nres10_D7 | atactacggtctcaaggaGCGACCGCGACCCTTACCCTGACCGTGACCGATGAAAACGGCGAAACCCTGACGCTGCAGCTGGATCTGACGGATTTTGATGCGGATGGTGTGGCAACCCCGACCTTAACCGCAGTGAGTGGTGGTCGTACCCCAGGTAGCGCAAGCGGCAAAGTGACCATTAAAGATGGCAAACTGACCTTTGATCTGACCCTGAGCTATGAACGCGATGGCGTGCGTTATGAAGGCCGCCTGAAAGGCGAAGGCACCTACGATG | MSGATATLTLTVTDENGETLTLQLDLTDFDADGVATPTLTAVSGGRTPGSASGKVTIKDGKLTFDLTLSYERDGVRYEGRLKGEGTYDAATGTATATATEGVLRRVEAGS |

| | | |
|---|---|---|
| | CAGCAACCGGTACTGCGACTGCAACCGCCACCGAAGGTGTTCTGCGTCGCGTGGAAGCGggttcccgagaccgtaatgc | GSHHWGSTHHHHHH |
| n7_S10_nres10_C8 | atactacggtctcaaggaGCGATTGATGCGGAAGGCACCTTTAAAGGTACGGTCGATGGTAAAGAAGTTACCGACGGCAAAGTGAAAGTGACCGGCCTGGTGCTCGAAACCGGCTATGAAGGCAAAGCGAAAGTTAAAATTCAGGTGGATGGCAAATGGCTGGGCGAAGGTACCCTGAGCTATAAAGTCACCGAATACAAAGACGGTAAGTTCAAATTCGACTTCACCATCACCAACCTGGATAACGGCAAATATAAAGGCAGCGGCACCGCGACCGGCGAAGTGGAAGGCGATAAAGCCAAAATTGATGTGAAAGGCACCATCGAAAGCACCGAAggttcccgagaccgtaatgc | MSGAIDAEGTFKGTVDGKEVTDGKVKVTGLVLETGYEGKAKVKIQVDGKWLGEGTLSYKVTEYKDGKFKFDFTITNLDNGKYKGSGTATGEVEGDKAKIDVKGTIESTEGSGSHHWGSTHHHHHH |
| n7_S10_nres10_D8 | atactacggtctcaaggaGCGAAGAAATTTACCGGCACCTTTAAAGCGACCGCGACGATTGATGGTAAGAAAGTGACCGGCGAAGGTGAAGTGGAAGGCGAAGTTAAAGATGGCGAAGTCACCTTCAAAATTACTAGCGGCGAACTGGGCGGTAAACCGATTGAAGGCGGTGGCGGCAAAGGCAAAGTGGATGAAAACGGCAACATTACCATTGATGAAGTTACGGCGACCATTGGTGGCGTTCCGGCGGAGAAAGGTACCGGGGAAGGTAAACTGTTTGATGATGGCACCGTGGAACTGAAACTGAAATTAGGCGCGGTGGATACCAGCGGCATTAAAggttcccgagaccgtaatgc | MSGAKKFTGTFKATATIDGKKVTGEGEVEGEVKDGEVTFKITSGELGGKPIEGGGGKGKVDENGNITIDEVTATIGGVPAEKGTGEGKLFDDGTVELKLKLGAVDTSGIKGSGSHHWGSTHHHHHH |
| n7_S10_nres10_C9 | atactacggtctcaaggaATGCGCGTGGAAGGCGAACTGGTTGTGATTGATGAAGATGGCACCCGCTATGAAGGCCCGGCGGTTCTGGATCTGGATGAGGATGGCAAAGGTACCCTGAAAGCGAAACTGTATCGCGATGGTAAACTGGTGCTGGAATTCGAAGGCGAGATCGAACTGGGCGAAGACGGCAAAACCTTTACCGTTGAAGCCGAAGGCACCCTGGAAGGTAAAGAAGTTCGCTTTCGCGGCGAAGGTGAAATCGAAATTGAAGGCGATAAATTTAAAGCGGAACTGCGCGCGGAATTTGAAGAAGTGGAACCGCCGCCGggttcccgagaccgtaatgc | MSGMRVEGELVVIDEDGTRYEGPAVLDLDEDGKGTLKAKLYRDGKLVLEFEGEIELGEDGKTFTVEAEGTLEGKEVRFRGEGEIEIEGDKFKAELRAEFEEVEPPPGSGSHHWGSTHHHHHH |
| n7_S10_nres10_D9 | atactacggtctcaaggaGAACCGTTTCGCTTTACCACCAGTGTGGATGGTACGCTGGATGGCACCGCCTTTACCAGCGGCACCGTGGATCTGAGCGTGTCTGATGTGGGCGATGGCAAAGCCACCATCACCTTCGATAGTGGCACCTTTGCGGGTAAAGCGTTTACCGGCGGTAGCGGCGAAGCGACCCTGACGAAAATTGGCGACGGCGTGTATAGCGCGAAAGGCACGTTTAGCGGTGTGGTGATTGATGGCGTGGAATATACCGGTACCTTTGAAGGCACCATTACCCTGGATAGCAACGGCA | MSGEPFRFTTSVDGTLDGTAFTSGTVDLSVSDVGDGKATITFDSGTFAGKAFTGGSGEATLTKIGDGVYSAKGTFSGVVIDGVEYTGTFEGTITLDSNGNVKSVKLTNITLTPAAGSGSHHWG |

| | | |
|---|---|---|
| | ACGTGAAAAGCGTGAAACTGACCAACATCACCCTGACCCCGGCGGCGggttcccgagaccgtaatgc | STHHHHHH |
| n7_S10_nres10_C10 | atactacggtctcaaggaATGCACATTACCGGCACCTTCACCTTTGATATCAACGTGAACGGCAAGAAATATAGCGGCAGCGGCGAATTTGTGACCGAAGAAGACGGCGTTACGTTTACCTTTAAAGGCACCACGGAAGATGGTAGCGTTATTGAAGGTACCGGCCGCCTGACCCGTCAGCCGGATGGCAGCATTACCTTAACGTTAACCGATATTAAAATTGATGGCAAAGCGGTTGGTGGTAGCGCGACTGGTCCGGCGACCTTACAGCCAGATGGTCATATTACCGCGACGGATATCACCGGCGAAATTACCGATggttcccgagaccgtaatgc | MSGMHITGTFTFDINVNGKKYSGSGEFVTEEDGVTFTFKGTTEDGSVIEGTGRLTRQPDGSITLTLTDIKIDGKAVGGSATGPATLQPDGHITATDITGEITDGSGSHHWGSTHHHHHH |
| n7_S10_nres10_D10 | atactacggtctcaaggaGGTAAAGCGAAATTAGTGATTAACATTAAAGGTGGCGTGGGCGGCGAAGTTGAAGAACTGGTGATCGAAGGCGAAATCGATCCGGAAACCGGCGAATTTGAAGGCGAAGGCACCGCGAAATATAAAGACGGCACCACCGTGAAAGGCAAAGCGGAAGGCAAACTGACCTTTGAAGATGGTAAACTGGTTAGCTTTGAACTGGAAGGTGAACTGGATGATGGCAGCACCTTTAAATTGACGGGCACCGGCACGGGTACCTTACCAGCGGTGGGTGAAACCGGTACTATTGATGCAACCGCGACGGGCGAACTGACCCCGGCGCCGggttcccgagaccgtaatgc | MSGGKAKLVINIKGGVGGEVEELVIEGEIDPETGEFEGEGTAKYKDGTTVKGKAEGKLTFEDGKLVSFELEGELDDGSTFKLTGTGTGTLPAVGETGTIDATATGELTPAPGSGSHHWGSTHHHHHH |
| n7_S10_nres11_C11 | atactacggtctcaaggaGCGAAAACCGAACACGTGAAATTTACCTTTAAATTCACCTTTACCCTGGATGGCGAAGAGAAAACCGGCACCGCGGAAGTGGATGCGGATCTGCCGACCGAAGTGGGTGGTAAAGGGAAAGGCAAAGGCGTTTTAAAAGACGAAAACGGTGAAAAGATTGGCGATTTCGAGGTGGACTTTGAACTGATCGGCGGCGATGGCGTGAAAGTGTATAAATTTGAATTCGAAGGCGAACTGAACGATGGTGCGAAAGGTAAAGGCAAGGGCACCGTGAAAACCAAACTGAAAGACGGCGAAAGCGGCGAAGTCGAAGCGGATGTGGAAGCCGATTTTAAGAAACATggttcccgagaccgtaatgc | MSGAKTEHVKFTFKFTFTLDGEEKTGTAEVDADLPTEVGGKGKGKGVLKDENGEKIGDFEVDFELIGGDGVKVYKFEFEGELNDGAKGKGKGTVKTKLKDGESGEVEADVEADFKKHGSGSHHWGSTHHHHHH |
| n7_S10_nres11_D11 | atactacggtctcaaggaAGCAGCCGCGATGTGACCGTGAACTATACCATTAAAGGCACCATTCTGGGCAAACCGTTTGATGCGGGCAAAGTTACCGCGCCGGGCACCTTAAGCGATGGCGTGCTGACCATTAACAGCAAATATGTGGGTAAAGCGGGCGGTAAACCGGTGACCGATGTTACCTTTACCCTGGATCTGGGCTCTGATGAAGTGGGCTATAGCGGTAGCGGTACCGGCACCGCGGAAGGTACCTACGATGGCAAGAAAGTGGCGGGTAACGTGACCTGGAAAGGCAAAATCGTTGCGCCGACCCTGATTGAACTGGATATTACCATTGAAGATTGGAAAGAAGTGCCGGAAGCGggttcccgagaccgtaat | MSGSSRDVTVNYTIKGTILGKPFDAGKVTAPGTLSDGVLTINSKYVGKAGGKPVTDVTFTLDLGSDEVGYSGSGTGTAEGTYDGKKVAGNVTWKGKIVAPTLIELDITIEDWKEVPEAGSGSHHWGSTHHHHHH |

| | | |
|---|---|---|
| | gc | |
| n7_S10_nres11_C12 | atactacggtctcaaggaACCAAAACCGGCACCGGCGAAGGCGAATTCGAAATTAACGTGGATGGCGAGAAAGTGAAGGGCAAATTCGAGTTTGAAATTACCATCAATGACGATAACGGCAACGGTACCGTGAAAGGTAACCTGTATGATGATGAAGGCAAGAAAATTGGCAGCTTTGAAGGCCCGGTGAAAGTTACCGCGCAGGCGGATGGCACCTACGATGTGAAATTTACCGATCTGAAAATCGATATCAACGGCGAATATAGCGGCAAAGGTGAAGGCGAAGGTAAAACCAAAGAAGTGAACGGCCAGCTGGGCAAAATTGAAGGTGCGAAACTGAAAGGCGAAGCGAAGAAAATCCCGGGCGTGggttcccgagaccgtaatgc | MSGTKTGTGEGEFEINVDGEKVKGKFEFEITINDDNGNGTVKGNLYDDEGKKIGSFEGPVKVTAQADGTYDVKFTDLKIDINGEYSGKGEGEGKTKEVNGQLGKIEGAKLKGEAKKIPGVGSGSHHWGSTHHHHHH |
| n7_S10_nres11_D12 | atactacggtctcaaggaGCGAAAACCCTGAAACTGGACGTGGATGTGGCGAAAGCGACCTACAACGGCAAAGAGTACGACAAAATGAAAGTTAAACTGACCATCGATAACCCGCCGACCAACGGCCAGAGCTTTACCGCGACCGGTGAACTGTATGATGCGAACGGTAACAAAATTGGCAACATTACCTTTAAAGGTACCGGCGATTTTACCAATGGCAAAACCGGCAGCGCCGAAGGTGAAGTGACCGACACCGATACAGGTGCGAAAGGCACCTTAAAACTGACCGATGTTAAAGTGAGCGGCGATACCGCCACGGCGACCGCGGATGAACTGGTGCTGAACAAACCAGGCAGCggttcccgagaccgtaatgc | MSGAKTLKLDVDVAKATYNGKEYDKMKVKLTIDNPPTNGQSFTATGELYDANGNKIGNITFKGTGDFTNGKTGSAEGEVTDTDTGAKGTLKLTDVKVSGDTATATADELVLNKPGSGSGSHHWGSTHHHHHH |
| n7_S10_nres12_E1 | atactacggtctcaaggaATGACCTATAAAGTGAAACTGGATGTGAAATTTAAAAGCGAACTGGGCACCGGCAAATTTACCGCGGTGGTGGATCTGGATTTTGATGAAAGCACCGGTAACGGCAGCGGTACCATTACCGATGTGAGCGGCGATGTGGGTGGCGTGAAAATTAAAGGCGGCAAAGGCACCATTAAATTCGAAGGCGGTGAAAAGAAAGACGGCGTCTACTATGTGAAGAACATCAAAGTCAAATACACCGATGATAACGGCAACGGCTTTGAAGGTGAAGGCGAAGGGGAAATTAAACTGAACGATGATGGCACCGCGGAAGGTAAACTGGATATTGAAGGCAAAGTGATTCTGCCGAAACCGggttcccgagaccgtaatgc | MSGMTYKVKLDVKFKSELGTGKFTAVVDLDFDESTGNGSGTITDVSGDVGGVKIKGGKGTIKFEGGEKKDGVYYVKNIKVKYTDDNGNGFEGEGEGEIKLNDDGTAEGKLDIEGKVILPKPGSGSHHWGSTHHHHHH |
| n7_S10_nres12_F1 | atactacggtctcaaggaATGACCAAAGTGCTTAAAGTGTATATGAAAGGCACCGACGAAGAAGGCCACGAGTTTGAAGTTGAAATGGAAATTGAAAACCCGAAAATCGATGAGGATGGCCGCTTTAGCGGCAAAGGCCCGGTGACCGTGACCATTAACGGCAAGAAATTTGAAGGCGAAGGGGAAGTCGAAGGCCGCATCGGCGAAGATGGCGTTACCGTGGAAGCGGAACTGCGTGCAGAAGGCACGGACGAAGACGGCACCAAATTCGAGGTTGAAGCGCGCGGTGAAGGCAAAGTGGAAGAA | MSGMTKVLKVYMKGTDEEGHEFEVEMEIENPKIDEDGRFSGKGPVTVTINGKKFEGEGEVEGRIGEDGVTVEAELRAEGTDEDGTKFEVEARGEGKVEEDENGVWNADLEATEAKVRIEP |

| | | |
|---|---|---|
| | GATGAAAACGGCGTCTGGAACGCGGATCTGGAAGCGACCGAAGCCAAAGTTCGCATTGAACCGCCGAAACCGggttcccgagaccgtaatgc | PKPGSGSHHWGSTHHHHHH |
| n7_S10_nres12_E2 | atactacggtctcaaggaATGAAAATGGATATCGATGTGACCATTAACCTGACCGATGAAAACGGCAAAGTTACCACCCTGAAAGCCAAAACCTATGAAGGCGAAGTGGATGTGGAGAAAGGTACCGTTAAATTTAAAGCGAAAATCTACGAAGGCACCGGCGGCAAATGGGAAGGTGAGTTCGAAGCGGAAGGCAAACTGACCAAACTGGCGGATAATAAATATCTGGTGCATTTCGAATTCGAAGGTGAATTTACCGATCCGAATGGCAACAAATATGAATTCAAGGGTGAAGGGGAAGGCGTTATCGAACAGAAAGGCGATACCTGGACCGGCGAACTGAAAGCAGAGGGCGAAGCGAAGAAACTGGAAggttcccgagaccgtaatgc | MSGMKMDIDVTINLTDENGKVTTLKAKTYEGEVDVEKGTVKFKAKIYEGTGGKWEGEFEAEGKLTKLADNKYLVHFEFEGEFTDPNGNKYEFKGEGEGVIEQKGDTWTGELKAEGEAKKLEGSGSHHWGSTHHHHHH |
| n7_S10_nres12_F2 | atactacggtctcaaggaGCGAAAATTACCGGTACCCTGAACCTGACCGGCACCAGCAAAGATGGCACCAAATTTAGCATCAACGGCAACATTACCGACTTTACCTATGATAAAGAGACGGGCAAAATTAGCTTTTCGGGCACCATTACTGGTGGTACCGTGGGTGGTGTGACCATTACCGGCGGCACCTTTAGCGGCGAAGGCACTGCGAAACTGTTAGAAGATGGGAAAATCACCGATATTGATGTGGATGTGACGATTAAAACCGATAACGGCTATGTGCTGAAAGGCAAATTCACCGGTGAAGGCTATTACGATAAAGAAACCGGCCATCTGACCCTGAACTTAAGCGGCAGCGGCTTTGAACTGACCAAACCGGATCCGAACCTGggttcccgagaccgtaatgc | MSGAKITGTLNLTGTSKDGTKFSINGNITDFTYDKETGKISFSGTITGGTVGGVTITGGTFSGEGTAKLLEDGKITDIDVDVTIKTDNGYVLKGKFTGEGYYDKETGHLTLNLSGSGFELTKPDPNLGSGSHHWGSTHHHHHH |
| n7_S10_nres13_E3 | atactacggtctcaaggaATGGTGACCGGCACCTTTGAAATGAACAACATTAAAGGCACCCTGGATGGCAAACCGTTTGAAGCGGAATATGCGAAAGGTACCCTGACCGATATCAAAATCGATTTTGAAAAGGGCATCATTACCGGCAAAGCGAACTTTACCATTCACTTTAAAGATGGCAGCAAACTGACCGTTACCGATGCGGATTTTAAAGCGACCGCGACCATTGATGAAGATGGTACCATTACCAACGGTAAAATTGAAGGCGAAGGCGAGGGCACCGATCAGAACGGCAAGAAATACAAAGTCAAATTTGAAGGTGACATCGTCGACGGCAAATATGACAAAGAGAAGAAACACCTGACCCTGAAAATTGACAACGTGAAAATGGAAATTACCAGCATTAAAggttcccgagaccgtaatgc | MSGMVTGTFEMNNIKGTLDGKPFEAEYAKGTLTDIKIDFEKGIITGKANFTIHFKDGSKLTVTDADFKATATIDEDGTITNGKIEGEGEGTDQNGKKYKVKFEGDIVDGKYDKEKKHLTLKIDNVKMEITSIKGSGSHHWGSTHHHHHH |
| n7_S10_nres14_F3 | atactacggtctcaaggaGCGGAGAAAACCAAAACCGATGTTGATATTGATGTGACCGGCACCGCGGATGGCAAACCGTTTAAACTGAAAGTGAAACAGGAAAACGCGCCGACCACCGTGACCAAGAACGATAACGGCGTGTATACCTTTGAAATTGATGTGGATAACGTTACCG | MSGAEKTKTDVDIDVTGTADGKPFKLKVKQENAPTTVTKNDNGVYTFEIDVDNVTGEIGGKKVEGGKA |

| | | |
|---|---|---|
| | GCGAAATTGGCGGCAAGAAAGTGGAAGGCGGCAAAGCGAAAATTAAAGGCAAATTCAAACTGACCGATCCGGAAGGTAAGAACGCGAAATTTGAGTTTGAAATCGAAATCGACGTGGAAGTGGATGGCGAGAAATATGGCGGCGTGGCGAAAGGTGAAGGCGAAGCCAAGACCAAAATGGAAGGTGATAAACTGGTGCTGACCGAACTGAAGGGTGAAGCCGAAGGCGAGCTGAAGAAAATTGAAGAAggttcccgagaccgtaatgc | KIKGKFKLTDPEGKNAKFEFEIEIDVEVDGEKYGGVAKGEGEAKTKMEGDKLVLTELKGEAEGELKKIEEGSGSHHWGSTHHHHHH |
| n7_S10_nres6_E4 | AGGCTTGCCGAATatactacggtctcaaggaATGAAAACCCCGCTGAAAGTGACCTATAAAGGCAAAGAATATGACGGCGAACTGAGCGAAGTGGTGGATGGTAAAGCGAAAGTTACCGTTACCATTGATGGCAAAACCTATACCGGCACCGTGGAATGGGACGGCAACAAATTTGAAGGTAAACTGGATGAAACCGGCGCGAAAATTAAAGGCGAGCTGAAAGATGGCGTCGTTTATGCGGAAATTGATCCGGATGGCATTCCGGAAggttcccgagaccgtaatgcAAATTTCTGCTT | MSGMKTPLKVTYKGKEYDGELSEVVDGKAKVTVTIDGKTYTGTVEWDGNKFEGKLDETGAKIKGELKDGVVYAEIDPDGIPEGSGSHHWGSTHHHHHH |
| n7_S10_nres7_F4 | atactacggtctcaaggaGGTGAAGCGTTTAAAGGCCTGGCGAACGGCACCATTGATGGCGTGCCGGTGACCAACTTACCGGTTGAAGGCAGCTTCGGCGATGGTACCATTACCATTTATGATCAGGATGGTAAGAAACTGGGCGAAGCGACCGTGAGCGGCAAAGATAACAAATTCAAGGGCGAAATTACCTATGACGGCAAGAAATACGAAGTTACCGGCGTGGTGAAGAAAACCGGCGAAGGCAAAGCGGAAGTGACCTTCGAAGCCAAAGAAGTGCCGGAAGAAggttcccgagaccgtaatgc | MSGGEAFKGLANGTIDGVPVTNLPVEGSFGDGTITIYDQDGKKLGEATVSGKDNKFKGEITYDGKKYEVTGVVKKTGEGKAEVTFEAKEVPEEGSGSHHWGSTHHHHHH |
| n7_S10_nres8_E5 | atactacggtctcaaggaGCGCAGGCGTTTACCGTGACCGGCACTGTTGATGGTCAGACCTTTGATAGCGGCAGCGCGACGGGCACCTGGAGCGGTAACGAAGGTCAGTTAACCAGTGGTACCCTGGGTGGTGTGCCGTTCACCGGTGGCACGCTGAAAGGTACCGATATTGGCAACGGCAAATTTACCTTTACCATTACCGGCCTGACCATTAACGGCGTGCTGTATAGTGGCACCGGTACCGCGACCTTAAACAGCGATGGCGTGTGGACCCTGAACCTGAATGTGACCACCGTGGCGGCGggttcccgagaccgtaatgc | MSGAQAFTVTGTVDGQTFDSGSATGTWSGNEGQLTSGTLGGVPFTGGTLKGTDIGNGKFTFTITGLTINGVLYSGTGTATLNSDGVWTLNLNVTTVAAGSGSHHWGSTHHHHHH |
| n7_S10_nres8_F5 | atactacggtctcaaggaGCGGAGAAACCGACCGTCGAAATTATTGGCACCGATGGCACCAAACTGACCATGGTGGTGACCGAAGGCGAACTGATTGGCGGCCGCGCGAAAGGTGAACTGTATCAGGATGGTAAGAAAGTGGGCGAAGGTACCCACGAAGTGAACGAAGATGCGACCACCGGCAGTGCGGAAGGCACCATTAACGGCAAACCGTTTCGCCATGAAGGTGAAGTGGTTGAAATTAGCCCGGATGGCAACAAAGTGGTGCTGGAAGGCCCGCTGGAAATTAAAggttcccgagaccgtaatgc | MSGAEKPTVEIIGTDGTKLTMVVTEGELIGGRAKGELYQDGKKVGEGTHEVNEDATTGSAEGTINGKPFRHEGEVVEISPDGNKVVLEGPLEIKGSGSHHWGSTHHHHHH |

| | | |
|---|---|---|
| n7_S10_nres8_E6 | atactacggtctcaaggaGCGCCGATTGCGGCGACCTTCACCGGCACCATTGACGGCAAAACCTTTACCAGCGGTACACTGACCGTTACCCGTGTGGGCACCACCGGTATTACCGGCGATGGCACGATTGATGGCGAGAAAGTGACCGGCTTTGTGTTTACCGGTGATGTGACGGGTGCGGGTACTCATAGCGGCACCGCGAGCATTGAAGAGAAAGGCTATACCGGTCCGGCCACCATTACCATTAGCGCGGATGGTACCACCGCGGATATTACGGCGGATCTGACCGCCGATAGCGCGggttcccgagaccgtaatgc | MSGAPIAATFTGTIDGKTFTSGTLTVTRVGTTGITGDGTIDGEKVTGFVFTGDVTGAGTHSGTASIEEKGYTGPATITISADGTTADITADLTADSAGSGSHHWGSTHHHHHH |
| n7_S10_nres9_F6 | atactacggtctcaaggaACCATTACCGGCGATTTTGAACTGACCGCGGAAGGCAAAACCTATAAAGGTACCTTTACCATCAACGGCAGCGTGGAAGAAGGCGCGGAATTTGAAGGCACCCTGAAAGGCAACGACGGCACCATTAGCAGCGGCAAATTTACCGCCAAAGTGGGCAAAGATGGCAAAGCGAAAGTGAAATTCACCGATATTACCATTAACGGCGAGAAATATGCGGGCGAAGGCGAAATGCAGATTACCAACGAAAACGGTAAAACCACCGTGAAAATTAAAGGCGATCTGAAGAAAGTGGAAggttcccgagaccgtaatgc | MSGTITGDFELTAEGKTYKGTFTINGSVEEGAEFEGTLKGNDGTISSGKFTAKVGKDGKAKVKFTDITINGEKYAGEGEMQITNENGKTTVKIKGDLKKVEGSGSHHWGSTHHHHHH |
| n7_S12_nres11_E7 | atactacggtctcaaggaAAGAAAATTCCGGAAATTATTGAAATTGATTTTACCATTAACGGCAAGAAATATCATATGACCATTGAACTGACCAAAGTGGAAGATCTGGGCGATGGCAAATATACCTTTACCGGCAAAATTAAAGATCTGACCGGCCATGGTGATGGCGAAGCGGAAGGCGAATTTGAATATGATGAAGAAGCGGGCGAAATTACCAAATTTAGCATTACCGGTGAAGTGGACGGCAAACTGTTTGATATTGAATTTAAACCGTATGATACCAAAGAATTTACGAAAGATGGTAAATATGGCCTGCGCTTTAAAGCGGATGCGAAAGGTGAATTTTATGATCCGggttcccgagaccgtaatgc | MSGKKIPEIIEIDFTINGKKYHMTIELTKVEDLGDGKYTFTGKIKDLTGHGDGEAEGEFEYDEEAGEITKFSITGEVDGKLFDIEFKPYDTKEFTKDGKYGLRFKADAKGEFYDPGSGSHHWGSTHHHHHH |
| n7_S12_nres12_F7 | atactacggtctcaaggaATGGAAGTGAAAACCAGCAAAGCCAAACTGAACATTACCGGTGAACTGGATGGCAAAGAATTTAAACTGGAAGCCGAAGGCACCGTGAAATACATCCTGGAAGAAGAAAACGGCAAAGCGCATCTGAAATTCGAAATGGATTTTGAAGGCGAACTGAATGGTAAACCGGCGACCGGCACCCTGAAAGGCGATCTGGAAGGCGAAGTTCTGGGCAACGGCCTGTATGAATTAGAAGGCCCGTTTACCCTGACCGTGGATGGTGATGTGTATGAAGGTGAAGCGAAAGGCATTGTCAAATTTGATGAAAATGGCTATCTGACCGAAATGGATCTCGAAGTGACCGGCGCGAAACTGGTGAAGAAAGCGCCGGGCAGCggttcccgagaccgtaatgc | MSGMEVKTSKAKLNITGELDGKEFKLEAEGTVKYILEEENGKAHLKFEMDFEGELNGKPATGTLKGDLEGEVLGNGLYELEGPFTLTVDGDVYEGEAKGIVKFDENGYLTEMDLEVTGAKLVKKAPGSGSGSHHWGSTHHHHHH |
| n7_S12_nres12_E8 | atactacggtctcaaggaACCAAAACGGTGCCGGTGAACCTGACCCTGAAAGGCGATGGCAAAACGCTGAAA | MSGTKTVPVNLTLKGDGKTLKLSGEFD |

| | | |
|---|---|---|
| | CTGAGCGGCGAATTTGATGTGGAACTGCGCCAGGAAGGTGATGAAACCGTGCTGCGTGGTAAAGGTCCGGTGACCGGTACCTGGGAAGGCAAACCGGTTACCGGTGAAATGGAATTTGAAGCGCGCGGCAAACTGGATGGCGATAAAGGCGACTTCGATCTTGAAGGCCGCATGAAAATTGACGGCAAATTGTATAAATTTCGTGGCAAAGCGACCGTTACCCTGCATCGCGATGAAAACGGCATTACCGGCATTGATATTGATGCGACCGATGTGAAAATCGAAGAAATTGAAGAAggttcccgagaccgtaatgc | VELRQEGDETVLRGKGPVTGTWEGKPVTGEMEFEARGKLDGDKGDFDLEGRMKIDGKLYKFRGKATVTLHRDENGITGIDIDATDVKIEEIEEGSGSHHWGSTHHHHHH |
| n7_S12_nres8_F8 | atactacggtctcaaggaGCGGCGAAAACCTTTAACGTGACCGGCACGTTTGATGGCGTGCCGATTGATCTGACCACCCTGGAATATACGATTGAAAACGGTAAACTGACCGTGAAAGGCAAACTGAACATTGGCGGCGTGGCGAAAGATTTTAAAGCGGAAGGCACCATCGAAGGCAACACCGCGACCCTGACGGGCACCTTAGATGGTGTTCCGGTTACCGCAACCCTGACCAACCTGGATATTAAAGATGATGAAATTCTGGGCACCGCCACCATTACCATGAAAACCGTGACCggttcccgagaccgtaatgc | MSGAAKTFNVTGTFDGVPIDLTTLEYTIENGKLTVKGKLNIGGVAKDFKAEGTIEGNTATLTGTLDGVPVTATLTNLDIKDDEILGTATITMKTVTGSGSHHWGSTHHHHHH |
| n7_S12_nres9_E9 | atactacggtctcaaggaACCACCTTTAAAGGTCTGACCATTACCGGCACCCTGGATGGCGAACCGTTTACCCTGACCACCGATAAACCGCTGGATGTGGCGGGCAACAACAAACCGTTCGAAATTAGCGGCGATTTTACGCTGAACGGCAAACCGGTGGATGCGACGCTGACCGGTACCATTAACCTGACTGAAGGTAGCGCGAACACCATTCGCGGCACCATTGATGGCCGCCCGTTTGAATTGCGTGGTGGCACCGTGAGCATTGAAGATGGCGTGCTGCGCATCGATGGCATGGAAGGCCGCTTTCTGGATggttcccgagaccgtaatgc | MSGTTFKGLTITGTLDGEPFTLTTDKPLDVAGNNKPFEISGDFTLNGKPVDATLTGTINLTEGSANTIRGTIDGRPFELRGGTVSIEDGVLRIDGMEGRFLDGSGSHHWGSTHHHHHH |
| n7_S12_nres9_F9 | atactacggtctcaaggaATGCAGACCACCATTAAAATTATTGGCGGCGAGCTGGATGGCGAACCAGTGGAAGGCGGTGAACTGGTGATTACCGAACGCGATGGCACCGAATTTACCGGCGAAGGCGAACTGGACGGCTTAGAACGTGGTCCGTTTCGTGGCCGTCATGATAAACTGGGTGAAGTGGGCGCGCGTGGTACCGCGGAAGGTGTGTTCGGTGGTTATCGTGTTCGCTTTGAATATGAAATTGTGGATGTGGCGGATGGTGTGATCACCGTTGAAATTACCGGTGGCACCCGCGAACCGGTGCCGGCGggttcccgagaccgtaatgc | MSGMQTTIKIIGGELDGEPVEGGELVITERDGTEFTGEGELDGLERGPFRGRHDKLGEVGARGTAEGVFGGYRVRFEYEIVDVADGVITVEITGGTREPVPAGSGSHHWGSTHHHHHH |
| n7_S12_nres9_E10 | atactacggtctcaaggaAGCAACACCGTCACCGGCAACCTGACCCTGACCGTGGATGGTGAAACCATGACCGGCACCGTGACCATGGATGCAGCGAATGCGGAAAAGGCGGGCGCAACTTTTACCTTCACGGGCGTGGTGTATCGCGGCAAACCGGTTAGTGGCACTGGC | MSGSNTVTGNLTLTVDGETMTGTVTMDAANAEKAGATFTFTGVVYRGKPVSGTGTLNFNAGEKGSWT |

| | | |
|---|---|---|
| | ACCCTGAACTTTAACGCGGGCGAAAAGGGCAGCTGGACCGTTACCTTTGAAGGCTATGAAGTTAAAGGCGAAGGCTTTATGAGCAAAGTGGATAGCAACAACAAATTTGATCTGGATCTGAAAGGTGGCGAACGCGTGCCGCTGCCGggttcccgagaccgtaatgc | VTFEGYEVKGEGFMSKVDSNNKFDLDLKGGERVPLPGSGSHHWGSTHHHHHH |
| n7_S12_nres9_F10 | atactacggtctcaaggaGCGAACGCGACCGGCACCTTTAAAGCGGATCTGGATGGCACCGTGGTTACCGGTACCTTCACCGTTACCAACCAGACTGCGGGCGGCAATGGCATCACGGGCACCCTGTATAACATTACGGGCCATGCGGATGGTACGTTTACCATTACCAATCTGACCAACGGCAAAGGTACCGTGACCCTGACCATTGATGGCGTGACCTATACCGGCAACGTGACCGTGGATAGCACCATCACCGCGGGCTCTACGGGTACTCTGACCTTTACCGATCTGAAACGCGTGGCGGCGGCGCCGggttcccgagaccgtaatgc | MSGANATGTFKADLDGTVVTGTFTVTNQTAGGNGITGTLYNITGHADGTFTITNLTNGKGTVTLTIDGVTYTGNVTVDSTITAGSTGTLTFTDLKRVAAAPGSGSHHWGSTHHHHHH |
| n7_S14_nres9_E11 | atactacggtctcaaggaGCGACCCAGACCATTGTGGTGACCACCCCGGATGGCGAAACCCATACCCTGCAGACCAACCTGGCTCTGAAACCGGGTGATACTGGTGGCACCGGTACTTTCTCTGGCACTGTGGCGGGTCGTGCGATTAGCGGTGGTACCCTGACCTTTGAACGTCTGGAACCGGGCGCGCGTTTTACCTTAACCGATGATGCGGGCAACGTGATTACCGGCCGTGTTGTGCGCGTTGAAGAAGGTGGCGCGATTCATCTGCAGGCGGATGCGGTGACGTTTGTGGATggttcccgagaccgtaatgc | MSGATQTIVVTTPDGETHTLQTNLALKPGDTGGTGTFSGTVAGRAISGGTLTFERLEPGARFTLTDDAGNVITGRVVRVEEGGAIHLQADAVTFVDGSGSHHWGSTHHHHHH |
| n7_S14_nres9_F11 | atactacggtctcaaggaATGGAGAAATTTAGCGGCGAAATTCGCATTCTGGATGAAAACGGCGAAGTTGTCTTTGAAGGCCGCTTCGAAGGTGAACAAGTTGGCGAAAACGAACAGGTTCTGCGCGCCGAAGGCGAATTTCGTGGCGAACCGTTTGCGTTTGAAGCGCGCGTGGAAGGCAAACTGGAACCGGGCAAAGAATTCGAAGCGGAAGGTACCTTAGGCGGTCGTCGCGTTGCGTTACGCCTGCGTGTGTTAGAATTTGGCGATGGCAAATTAGTGGTGCAGCTGCTGGAATTCCGTGAAGTGGCGGCGggttcccgagaccgtaatgc | MSGMEKFSGEIRILDENGEVVFEGRFEGEQVGENEQVLRAEGEFRGEPFAFEARVEGKLEPGKEFEAEGTLGGRRVALRLRVLEFGDGKLVVQLLEFREVAAGSGSHHWGSTHHHHHH |
| n7_S8_nres10_E12 | atactacggtctcaaggaAAAGTTAAAGCGACCGGCGTGATTCCGGTGCGTGGTGAACTGAGCGGCGTTCCATTTGAAGGTGAAATGAAAGTGGATTTCGATTTTGAAGCGGGCAAAAGCGGCGAATTTGATTTTACCCTGACCCTGGATATTGATGGTAAAACCCATGTGGCGGTGGGCAAAGGCAAATACGATGCGATTAACGAGGGCGAAACCGGTAAAGGCGAGTTCGGCTTTTATGATAAAGATACGGGCGTTACCGGCGTTGGCGAAGTGAAATTTACCGTGAAAGATGGCAAAATCACCGGCGAAGCGCGCGCGGAAGGCGAAATTCCGAAGAAAg | MSGKVKATGVIPVRGELSGVPFEGEMKVDFDFEAGKSGEFDFTLTLDIDGKTHVAVGKGKYDAINEGETGKGEFGFYDKDTGVTGVGEVKFTVKDGKITGEARAEGEIPKKGSGSHHWGSTHHHHHH |

| | | |
|---|---|---|
| | gttcccgagaccgtaatgc | |
| n7_S8_nres11_F12 | atactacggtctcaaggaATGACCGTGACCGTTGAAGCGGAAGTGACCGGCAACATTGGCGGCGAGAAATTTAAAGGCGAGGCGAAAATCACCCTGAAAATTACCGAAAAGGATGGCGTTCTGACCTTTGAAGGGGAAGGCGAAGTTACCCTGCGCTTTGAAAACGGCCTGGTTCTGAAAGGCAAAATTATTGATCTGAAGGGTGTGATTAAAGAAGACGGCAGCTTCAAAGGCAAAGGTAAATTCGAAGGTGAAAACGATGGCATTAAAGCGAAGGGCGAAGCGTATATTGAAGGCAAGCTGGTTGATGGCAAACTGGTGGATGTGAAAGTGAAAATCGATGGCGAACTGGAGAAAGTGGAAGAAggttcccgagaccgtaatgc | MSGMTVTVEAEVTGNIGGEKFKGEAKITLKITEKDGVLTFEGEGEVTLRFENGLVLKGKIIDLKGVIKEDGSFKGKGKFEGENDGIKAKGEAYIEGKLVDGKLVDVKVKIDGELEKVEEGSGSHHWGSTHHHHHH |
| n7_S8_nres11_G1 | atactacggtctcaaggaATGAAAGTTGAACTGGATGCGGAAGCGGATCTGAAAGACGAAAATGGCAACCTGGTGAGCAAATTTAAGTTTAAACTGGAAGTGGAAATTGATGAGAACGGTGAAGTGACCGGCGAAGCGGAATTTAAAGATCAGGGCGGCATCAAAGAAATGAAGATGGAAGTGAAAGGCAAATATGATAAAGAAACCGGCAAAATTACCGGGGAAGTCGAAGGTGAATTTGTGGATGAAAACGGCAAGAAACATAAATTCAAAGGCGAATTTGAAGGCAAAGTGGATCTGGAGAAAGGTAAAGGCACCATGAAAATGGACCTGACCCTGCTGCCGGAAGAGAAAggttcccgagaccgtaatgc | MSGMKVELDAEADLKDENGNLVSKFKFKLEVEIDENGEVTGEAEFKDQGGIKEMKMEVKGKYDKETGKITGEVEGEFVDENGKKHKFKGEFEGKVDLEKGKGTMKMDLTLLPEEKGSGSHHWGSTHHHHHH |
| n7_S8_nres12_H1 | atactacggtctcaaggaATGAAACTGAAATTGAAAGCGATTGCGGATTTTGATCTGGTGGATGAAGATGGCAAGAAAATTGGTACCATGAAAGCCGAATTCGAAGCGGATGTGCTGGAAGGCGGTAAACTGGAGAAAGCGCCGTTTACCTTTCGCGGCGAAATTAACGGTCAGGATGTTAGCGGTACCGGTGAAGCGCAGGGCCGCGTGGAATTAGAAGAAGGTGGCAAAGTTAAAGCGGAAGTGGATTTTAAAATCACCGAAAACAACCTGGGCTTTCAGGGTGAAGGCACCGCGCGTGGCGAAGGCAAATTGGAAGATGGTGTCGCGACCGTGGATATGGAAGGCACCATTACCGGCAAACTGCCGGAAggttcccgagaccgtaatgc | MSGMKLKLKAIADFDLVDEDGKKIGTMKAEFEADVLEGGKLEKAPFTFRGEINGQDVSGTGEAQGRVELEEGGKVKAEVDFKITENNLGFQGEGTARGEGKLEDGVATVDMEGTITGKLPEGSGSHHWGSTHHHHHH |
| n7_S8_nres13_G2 | atactacggtctcaaggaAGCACCAAATATACCGGCAAATTTGAAGCGACCGTGACCGGTACCATTGATGGTAAACCGTTTAAAGCGAAACTGAGCGCGGAATATGAAGCAGAAGTGGATAACCCGGAAGACGGCAAACCGTTCACCTTTCGTGGCCGCTTTCGCGGCGAACTGACCTTTGATGATGGCACCGTGGTGCCGGTGGAAGGCGAATTCGAAGGCAAAGCGGAAGGTCCGCTGAGCGATTTCACCGGCGAAGTGAAAGTGTATATCAAAATGAAAATCGATGGCAAACTGTACACCGGCAG | MSGSTKYTGKFEATVTGTIDGKPFKAKLSAEYEAEVDNPEDGKPFTFRGRFRGELTFDDGTVVPVEGEFEGKAEGPLSDFTGEVKVYIKMKIDGKLYTGSGTTKFTGKIDTEKKVLDLDLGTI |

| | | |
|---|---|---|
| | CGGCACCACCAAATTTACCGGTAAAATTGATACCGAAAAGAAAGTGCTGGATCTGGATCTGGGCACCATTACCGTGGATTTTACCGAAGCCGAAGAAggttcccgagaccgtaatgc | TVDFTEAEEGSGSHHWGSTHHHHHH |
| n7_S8_nres6_H2 | ACTGCCCatactacggtctcaaggaACCGAAGAACGCGTTCGCGGTGAACTGGATGGCGTTCCGTTCGAAGGTGTTCTGCGCGGTCCAGAAGAAGGTCGCGTCCGCATTGAAGGCGAAGTGGATGGTCAGCCATTCGAAGCCGAGGGCGAACGCGAAGGCGATCGTATTCGCCGCTTTCGTGGTGAATTAGATGGCCGCGAATTTGAAGGTGAAGGTCGTCGCGTTGCGCCGGGTTTATGGGAAGCGCGTGCAGAAACCTTGCCGGTGCTGGAAggttcccgagaccgtaatgcTACCGG | MSGTEERVRGELDGVPFEGVLRGPEEGRVRIEGEVDGQPFEAEGEREGDRIRRFRGELDGREFEGEGRRVAPGLWEARAETLPVLEGSGSHHWGSTHHHHHH |
| n7_S8_nres8_G3 | atactacggtctcaaggaATGATGAAAGTTAAAGTCGATGTGGAAGGCACCCTGGGCGGCAAATATAAAGTGAAAGCGGTGTTAGAAGGCAACGACGATGGCAGTAAAGCGCGTGGCGAGGGCGAAATCGAAGGCGGCACCAAAATTGAAGTTGAGGGCACCATTAAAGATGGTAAAGTTACCGACATCAAAGCCGAAGCGGAACTGCCGGATGGCAACAAAATTCTGGGTGAGGGTGAAGGCGAATACGATGAAAACGGCAAAGGCAAAGCGACCCTGGAAGGTGAAGTTATTAAGAAAGAAAAGggttcccgagaccgtaatgc | MSGMMKVKVDVEGTLGGKYKVKAVLEGNDDGSKARGEGEIEGGTKIEVEGTIKDGKVTDIKAEAELPDGNKILGEGEGEYDENGKGKATLEGEVIKKEKGSGSHHWGSTHHHHHH |
| n7_S8_nres9_H3 | atactacggtctcaaggaATGCCGAAAACTGTTCGCGTGCGTGTGCGCGGCGAAGTGGATGGCGTGGAAATTGAAGGGGAAGTGGTTCTGGAAGGCGAAGAAATTGACGGCGTCTTTGTGGGCGAAGGTGAGGGAGAAGGCGAAACCGCAGATGGTCGCCGTTTCCGTGGTCGTTTGCGCGCGACCTTACCACGTGGTGAAGTGGGTGAAGGTGAACTGGAAGCGGAATTCGAAGATGGCACCCGCTTTCGCGGCCGCGTTCGTGGTCAGGTTGATGGTGATGAAGGCGAGGGCGAAATCGTTGGCGGTGAATTTATTCCGCCGAAAggttcccgagaccgtaatgc | MSGMPKTVRVRVRGEVDGVEIEGEVVLEGEEIDGVFVGEGEGEGETADGRRFRGRLRATLPRGEVGEGELEAEFEDGTRFRGRVRGQVDGDEGEGEIVGGEFIPPKGSGSHHWGSTHHHHHH |
| n8_S10_nres7_G4 | atactacggtctcaaggaATGAAGAAATACAAATTCACCATTACCCTGAACGATGGCACCAAATATACCGGCGAAATGGAAGTGAACGAAGATGGCGTGGTTCGCTTTGAAGGTGAGGGCGAAGACGGTACCAAAGTGGTTGGCGAAGGCGAAGTGGATCTGGAAACCGGTGAAGGCACCGCGGAACTGGAAGCGGATGGCGGCAAAACCAAATGGAAAGCGACCTTTAACCTGAAGAAGGGCACCGGCGAGATCTATGATGAAAACGGCAACAAAGTTGGCACCGTGAAAATTGAAGAAGTTGAAGTCGAAGAAggttcccgagaccgtaatgc | MSGMKKYKFTITLNDGTKYTGEMEVNEDGVVRFEGEGEDGTKVVGEGEVDLETGEGTAELEADGGKTKWKATFNLKKGTGEIYDENGNKVGTVKIEEVEVEEGSGSHHWGSTHHHHHH |
| n8_S10_nres8_H4 | atactacggtctcaaggaGAAAAGAAAACCCTGAAAGGC | MSGEKKTLKGTFK |

| | | |
|---|---|---|
| | ACCTTTAAAAGCAGCGACGGCAAACTGAGCGGCGAAATTACCCTGGAAGGCGATTTTCAACCGGGCGGTAAAGCGACCTTTGAAGCGAAAGGCGAACTGGAAGGTCAGCCGGTTACCGTGAAAGGTGAGGCGAAAGTGGGTGAAGGCGGTCGCCTGTCTGGCGAAGGTGAAGCAGATGTTGGTGGCAAGAAATATCCGGCGACCCTGGATGTGGGTGCGGATGGCAAAGGTACGCTGAAAATTGGCGAGGGCGGCAACGTTTATACCGCGGATGTGACGCTGGATGAAAAGATTGAACTGAAACCGCCGGGCGAAggttcccgagaccgtaatgc | SSDGKLSGEITLEGDFQPGGKATFEAKGELEGQPVTVKGEAKVGEGGRLSGEGEADVGGKKYPATLDVGADGKGTLKIGEGGNVYTADVTLDEKIELKPPGEGSGSHHWGSTHHHHHH |
| n8_S10_nres8_G5 | atactacggtctcaaggaATGTATTATAAAGGTACCTTCGAGGGTGAAATCGATGGCGTGAAATTTAAAGGCGAATTCGTTGTGGACCTGGACAACAACTATTTTCAGATGAAAGCGGTGGACGAAGATGGCAAACTGCTGAAAGCGAAAGGCGAGATCGATCTGGAAAACGGCACCTTTAAATTTGAAGCGGAATATGATGGTGGCAAAGGCGAAGGGGAAGGCGAGGGCGATCTGACCAAAGTTGGCGGCAAATTCAAATTCGAAGGTGATATCGGCGGTAAGAAAGTGAAAGGAGAAGGCGAAATTACCGAAGAAATTGACGAGTTTGAAGAACTGTTTggttcccgagaccgtaatgc | MSGMYYKGTFEGEIDGVKFKGEFVVDLDNNYFQMKAVDEDGKLLKAKGEIDLENGTFKFEAEYDGGKGEGEGEGDLTKVGGKFKFEGDIGGKKVKGEGEITEEIDEFEELFGSGSHHWGSTHHHHHH |
| n8_S10_nres8_H5 | atactacggtctcaaggaGCGAAAATTGTGAAAGGTGAGTTCGAATTGGAAAACGGCACCAAAGGCAAGTTCGAGGGTGAAGTGAAAGACGGCAAAGGCGAATTCGAGGGCGAAATCGATAAAGATGGTGTTAAAGGTGTGTTTAAGGGCGAGGTGAAACTGGATGAAGATGGCAAAATTTGGGGTGAATTTGAAGGTGAACTGGGTGGCAAAGATTTCAAAGGTAAATTCGAAGGCAACGTGAAAGGCGATGGTAAAGGCACCATCGAACTGGACATCGATGGCGAGAAAATTAAAGGCGAAATGTACGTTAAAGTGGAAGATGCGCCGAAACCGGTGCCGggttcccgagaccgtaatgc | MSGAKIVKGEFELENGTKGKFEGEVKDGKGEFEGEIDKDGVKGVFKGEVKLDEDGKIWGEFEGELGGKDFKGKFEGNVKGDGKGTIELDIDGEKIKGEMYVKVEDAPKPVPGSGSHHWGSTHHHHHH |
| n8_S10_nres8_G6 | atactacggtctcaaggaAAGAAAATCACCGGCGGCAAATTTGAAGGCGAAGTGGGTGGCAAGAAAGTGGTGGTCGATATTACCGATGTGAAAGATCTGGGCGATGGCAAATATGAAGCGACGTTTAAAGCGACCGGCCTGGGCGGTGATGGTAAACTGACCTTTAAACTGTTTGAAGATGGCAGCATCAAAGGTGAATTCACCGTGGATGCGAACGGTAAGAAAGGCACCTTCGAAGGTAAAATCGATGAAAACGGCGAAGGTGAAGTGGAACTGGATTTTGATGGCCAGAAAGGTAAAGGCAAAATTACCCTGACCCTGGAAGAAggttcccgagaccgtaatgc | MSGKKITGGKFEGEVGGKKVVVDITDVKDLGDGKYEATFKATGLGGDGKLTFKLFEDGSIKGEFTVDANGKKGTFEGKIDENGEGEVELDFDGQKGKGKITLTLEEGSGSHHWGSTHHHHHH |
| n8_S12_nres6_H6 | atactacggtctcaaggaACCACCGTGAAAGTGACCGGCACGATTGGCGGTCAGGCGGTGGATCTGACCGGTACTGAAATTGCGCCGGGCGTGTATCAGGTGGAA | MSGTTVKVTGTIGGQAVDLTGTEIAPGVYQVEGTIGGKAFS |

| | | |
|---|---|---|
| | GGCACCATCGGTGGTAAAGCGTTTAGCGGCACCCTGACCGCGGCGAATGGTACCTGGAGCCTGAAAGGTACTCTGGATGGCAAACCGTATGATGCGGGCAGCGGTACACTGAGTGCTGGCGGTGCGTTTTCTGGCACTGATGCAACTACCGGTGCGGCAGTTTCTGGTACCATTACCGCGACGGCGGCGggttcccgagaccgtaatgc | GTLTAANGTWSLKGTLDGKPYDAGSGTLSAGGAFSGTDATTGAAVSGTITATAAGSGSHHWGSTHHHHHH |
| n8_S12_nres8_G7 | atactacggtctcaaggaAAGAAATACACCTTTAAAGCGGAAATTGATATCAACGGCGAGAAACATGTGCTGACCGGCGAACTGGAAAAGGAAGGCAACACCTATAGCGGCAAAATTACCGATGGTACCATCTTTGGCAAACCGTTTACCGGTACCTTTGAAAACGTGGAAATTGGCGGTCCGGGTAAATGGTTCCCGGGTACCCTGGGTGGTAAACCGATGCGCTTTAAAATCGCGATCGATGAACTGGATGAAACCGGCACCATTCATTTTGAATTGGAAAACGGCCTGAAAGGCGAAGGCACGATTACCATCACCGAAGTGCCGCCGggttcccgagaccgtaatgc | MSGKKYTFKAEIDINGEKHVLTGELEKEGNTYSGKITDGTIFGKPFTGTFENVEIGGPGKWFPGTLGGKPMRFKIAIDELDETGTIHFELENGLKGEGTITITEVPPGSGSHHWGSTHHHHHH |
| n8_S12_nres8_H7 | atactacggtctcaaggaACCCGCTACTATGTGAAATTTGAATTTGAAGCGGATGGTAAGAAATTCGTGGTCGAAGGTGATGCGGGCAAACTGGGCGAATGGGGCCCAGCGACTATTACCGTCGATGGCAAAGCGTATCCGGGCGAAATTAAAATTACCGCGATTGATGGCGATAAAGTGGTGGTGGAATTCAAAGGCGAGATCGACGGCAAACCGATTGAAGGCAAAGGTACCTTTACCGTGAAAGATGGTAAAGGTGAAGGTAAAGTGGAAATGGAACTGGATGGCAAGAAACTGGAAGGCGAAGGCACCATGGAACTGAAAGAAGTGggttcccgagaccgtaatgc | MSGTRYYVKFEFEADGKKFVVEGDAGKLGEWGPATITVDGKAYPGEIKITAIDGDKVVVEFKGEIDGKPIEGKGTFTVKDGKGEGKVEMELDGKKLEGEGTMELKEVGSGSHHWGSTHHHHHH |
| n8_S14_nres8_G8 | atactacggtctcaaggaACCAAAACCTTTGAAGTGTATGATGCGAACGGCAACAAAATTGATACCTTTACCGTGGTGTATCGCGATGGCAAAACCATTGTTTGGGTGGATAGCAACGGCAATAAAATTGTGTGGGAACGCGTGGGCGATGATAAGAACCGCATTACCGCGACCGATGAAAACGGTAAAGTTACCAACTGGACCATTCTGGAATTTGAAAATGGCGAATTACTGGTGGTGAAGAACGAAGAAACCGGCGAAGTGATTACCTTCAAAGGCGATACCGGCTGGAGCGAAGAAGGCTTTggttcccgagaccgtaatgc | MSGTKTFEVYDANGNKIDTFTVVYRDGKTIVWVDSNGNKIVWERVGDDKNRITATDENGKVTNWTILEFENGELLVVKNEETGEVITFKGDTGWSEEGFGSGSHHWGSTHHHHHH |
| n8_S8_nres11_H8 | atactacggtctcaaggaAGCAAAATCCGCATCTACAAGGGCAAGTTTAAATTCAAGAACAAGAAAACCGGCGAAACGGGTGAAGGCGAGGCGGAAGCGGAAATCGATGAAAGCAACGATTACAAAGGCACCTTCAAATTTGAAGCGAAGAACGATGATACCGGCGAGAAATTTAAAGGGGAAGGGAAAGTTAAACTGGATCCGGAAACCGGTGAAGTGACCCTGACCGATCTGGAAGTG | MSGSKIRIYKGKFKFKNKKTGETGEGEAEAEIDESNDYKGTFKFEAKNDDTGEKFKGEGKVKLDPETGEVTLTDLEVENLETGVKGKGEGKGGK |

| | | |
|---|---|---|
| | GAAAACCTGGAAACAGGCGTTAAAGGCAAGGGC<br>GAAGGCAAAGGCGGCAAAATTAAAGGCGAAGAG<br>GGCTTCGATTTCAAGTTTGAAATCAAACTGGATAA<br>CGGTGAAACCCTGGTGGCCGAAGGTAAACTGAA<br>ACTGGTGGAAGTGCGCGAAggttcccgagaccgtaatgc | IKGEEGFDFKFEIKL<br>DNGETLVAEGKLKL<br>VEVREGSGSHHW<br>GSTHHHHHH |
| n8_S8_nres12_G9 | atactacggtctcaaggaATGCATCTGAAAGGCGAGGGC<br>AAAGTGAAACTGGACGGCGATGGTGATGAATTTA<br>AAGGTGAAGGTAAAGCGTATGTGGATAGCGATCT<br>GGATGAAAACGGCAACGGCTTCAAACTGAAATTC<br>GAAGGGGAAGTTAAATTCAAAGATGGCACCGTTA<br>ACAAATTCGTGGGCGAAACCAACGATGGCGTGC<br>TTAAAATTGAAGATGGTAAAGGCAAACTGCAGCT<br>GCCGGTGAAATTTCGCTTTCTGGATAAAGGCGGC<br>GTGGGTGAAGCGAAATTTGATGGCGGTGAAGTG<br>ACCGTGGAAGACGGCAAATTTAAACTGGATGCCG<br>AAGCGGATTTTGAAGCCGAACTGGATGATGGTAC<br>CAAATTCAGCGGCAAAATGAAATTGGATGTGGAA<br>GGCGAAGTCGAAGAAGTGGAAggttcccgagaccgtaat<br>gc | MSGMHLKGEGKVK<br>LDGDGDEFKGEGK<br>AYVDSDLDENGNG<br>FKLKFEGEVKFKDG<br>TVNKFVGETNDGV<br>LKIEDGKGKLQLPV<br>KFRFLDKGGVGEA<br>KFDGGEVTVEDGK<br>FKLDAEADFEAELD<br>DGTKFSGKMKLDV<br>EGEVEEVEGSGSH<br>HWGSTHHHHHH |
| n8_S8_nres12_H9 | atactacggtctcaaggaGCGAACAAAGTGCTGGAAGCC<br>GAAGGCACCGTGAAATTTGAAGGCGATCTGAAC<br>GGCCAGCATTTTAGCGGCGAGGGCAAAGCGAAA<br>GCGATTTACGATGAAGGCAACAAAGAACTGAAAG<br>TAGAATTCGAAGCTGAAGGTAAACTGAATGGCCA<br>GGATATTGGTATGAAAGGCGAAGGTACCATCAAA<br>GGTGTGGATCTGAGCAAGGGCTATTTTAAAGGCA<br>AAATGGACATGAAAGTGGATATTACCGGCCTGGG<br>CGGTAAAGGTGAAGTGACCGAAGCAGAATTAAAA<br>GATAACGGCGGCAAAGGTTGGGTGATTGAAGCG<br>GAAGGTGAAGCGCAGACCCCGCTGGGCAACTTC<br>AAGTTCAAAATCGAGCTGGACCTGGACAAGTTTA<br>AAGAGATTAAAggttcccgagaccgtaatgc | MSGANKVLEAEGT<br>VKFEGDLNGQHFS<br>GEGKAKAIYDEGN<br>KELKVEFEAEGKLN<br>GQDIGMKGEGTIK<br>GVDLSKGYFKGKM<br>DMKVDITGLGGKG<br>EVTEAELKDNGGK<br>GWVIEAEGEAQTP<br>LGNFKFKIELDLDK<br>FKEIKGSGSHHWG<br>STHHHHHH |
| n8_S8_nres6_G10 | atactacggtctcaaggaAGCAAGAAAGTGCTGTTCGAA<br>GGCGAAGTGGATGGTGTGCCGTTTACCGGCGAA<br>TTTGATGTTGTTCCGGATGGCGCGCCAGCGACC<br>GCCACCATTAACCTGAATGGCAAAGATCATGTGTT<br>CACCGGCACCTTTAGCAACAACACCTTTGTGGGC<br>ACCGATGCGGCGACCGGCTGGAAACTGACCGTT<br>AAAATTGATGGCGATACCGCGACCCTGACCCTGG<br>ATAAGAACGGCAAGAAATATCAGGCCGAAGGCAA<br>AGTGGTTGAACCGCCGggttcccgagaccgtaatgc | MSGSKKVLFEGEV<br>DGVPFTGEFDVVP<br>DGAPATATINLNGK<br>DHVFTGTFSNNTFV<br>GTDAATGWKLTVKI<br>DGDTATLTLDKNGK<br>KYQAEGKVVEPPG<br>SGSHHWGSTHHH<br>HHH |
| n8_S8_nres6_H10 | CTACTatactacggtctcaaggaGCGAAGAAAGTGGAATT<br>TAGCCTGACCGATGCGAACGGCAATACCATTCAG<br>GGCACCGCGGAAGAAGGCAAAGAACTGCCGGTT<br>TATGATCAGAATGGCAACCTGGTGGGTACCGCGT | MSGAKKVEFSLTDA<br>NGNTIQGTAEEGKE<br>LPVYDQNGNLVGT<br>AFIKDGELYFKDNN |

| | | |
|---|---|---|
| | TTATTAAAGATGGCGAGCTGTATTTCAAGGATAACAACGGTAACGTGTATAAAATCGAAGTGGACGGCGATAAAGCGACCGCGACCATTAACGGCGTGGAATATAGCGGCACCGTGAAAGAAATTGAAGCGCCGggttcccgagaccgtaatgcGTCAT | GNVYKIEVDGDKATATINGVEYSGTVKEIEAPGSGSHHWGSTHHHHHH |
| n8_S8_nres6_G11 | atactacggtctcaaggaACCTATACCGTGACCGGCGAATGGAGCGATGGTAAGAAAACCAGCGGCACCCTGACCCTGAGCCCGGGCAACAAATTTACGCTGACCTTTGATGATGGTCTGGTTCTGACCGGTAATTGGCCGGCGGCAGGTCAGCCGTTTAGCGGCAAAGATCAGAACGGTAACCCGTTTACCGGTACCAGCGCGGCGGCGGGTCAGGAATTTACCATTACCATCGATGATAACGGCAAGAAAACCGTGTTTAAAGGCCTGCTGACCGCGGATCCGGATAGCggttcccgagaccgtaatgc | MSGTYTVTGEWSDGKKTSGTLTLSPGNKFTLTFDDGLVLTGNWPAAGQPFSGKDQNGNPFTGTSAAAGQEFTITIDDNGKKTVFKGLLTADPDSGSGSHHWGSTHHHHHH |
| n8_S8_nres6_H11 | AatactacggtctcaaggaGCGATTGTGGGCGAAGCGAAGAACAAAGACGGCAACACCGCGAAATTTCGCATTGAAGATGGCAAAGTCAAAGGCGAAGTGACCGTGAACGGTAAAACCTATGAATTTGAAGCGGAACTGATTGATGGCACCAAATTCGAAGGCAAAATTAAAGGCGCGGACGGCACCGTGAGCGGCGAAATCGAAGATGGTAAAGTTAAAGTGGAAGTGGATCTGAACGGCAAAACGGTGAAACTGGAAGGCACCTACGAGAAAGTTGAAggttcccgagaccgtaatgc | MSGAIVGEAKNKDGNTAKFRIEDGKVKGEVTVNGKTYEFEAELIDGTKFEGKIKGADGTVSGEIEDGKVKVEVDLNGKTVKLEGTYEKVEGSGSHHWGSTHHHHHH |
| n9_S10_nres6_G12 | atactacggtctcaaggaAAACCGACCGTGACCTTTAAAGCGGATGATGGCACCACCGGTAAAGGCGAAGTGGAGATTGGCGATAAAATTGAATTTAAAGGTGAACTGAGCGATGGCAATACCTTCAAAGGCGTGATCGATCCGAAAGCGGGCACCTTTACCGCGCAGAACACCACTACCGGTCTGACCGCGAAAGGCAAAGTCGAAGGCAACACGTTTAGCGGCACGTTTGATGACGGCACCAAATTCAAGGGCGAAATCAAAGAAGATGGCAACGTTTATATTTATGCGCAGACCGCGCCGAAAggttcccgagaccgtaatgc | MSGKPTVTFKADDGTTGKGEVEIGDKIEFKGELSDGNTFKGVIDPKAGTFTAQNTTTGLTAKGKVEGNTFSGTFDDGTKFKGEIKEDGNVYIYAQTAPKGSGSHHWGSTHHHHHH |
| n9_S10_nres6_H12 | atactacggtctcaaggaACCATTACCTTTAAAGGCACCGTGCTGGGCAAAGAAATTACCGGCACTGTGACTGGCACTCCTGGCACCGGTACTGCGACCGCGGATTTAGGCGGTGAACATTTTACGGGTACCATTAGCGCGGGCCCGCTGACCGTTACCCTGACCAACGCCAAAGGTGATGTGATTACCATCAAAGTTGATGGCACCAAAGTGACCGTGAGCGGCGATGTGGATGGCAAGAAATTTGAAGGCGAAGGCACGTATACCCCGGGTAGCGATCATATTGATCTGGGCGTGGCGGAAGAAATCCCGGGCGTGggttcccgagaccgtaatgc | MSGTITFKGTVLGKEITGTVTGTPGTGTATADLGGEHFTGTISAGPLTVTLTNAKGDVITIKVDGTKVTVSGDVDGKKFEGEGTYTPGSDHIDLGVAEEIPGVGSGSHHWGSTHHHHHH |
| TMB_n12_S14_nres14_2 | atactacggtctcaaggaAGCAAAGAGAAAGAATACGGCGCGGAACTGGGCCTGGGCTGGCGCGACAACAAATA | MGSSKEKEYGAELGLGWRDNKYYIYAG |

| | | |
|---|---|---|
| | CTATATTTACGCGGGCACCTATAGCGAAGATGGTGA<br>AAAGCGCGACGAAGAAAAGAAGAAAGAAAGCCTG<br>GGCGTTGATCTGGGCCTGGGCCTGAAAGAAGAAC<br>GCAAATATTATGTGTATGTGGGCGCGTATGTGGAAT<br>TTGGCGAAAGCGATGATAAGGACAAAGAACTGTAC<br>GTCGGCAGCTATCTGAAACTGGGTCTGGATGATGA<br>ATGGGAAGCGAAACTGGGCCTGTATGCGGGCGGC<br>AAATACAAAGACAAAGATAAAGACTTCGAAATTTAC<br>GTGGGCGTGTATGGCGGTGCGGGTGCGGATAAAA<br>CCAAGAAACTGTATTACGAATTCGGCGTTTATACCG<br>GCCTGGGTTTTGATAGCAAGAAGGATAAAGAGAAA<br>GGCTATTTCGGCACCGGTCTGGGCGGCCAGTATAA<br>AGATAACCAGTACAAATATCGCGGCGATATTGAAGC<br>GGGCATTGGCTATGATGATTCCAAGAAAGATAGCG<br>CCTATGTGAAAATTTATGCGGGTTGGGATGATCGCA<br>ATAAATTCGAAGCGGGTCTGGGTAGCGGCTTTGCG<br>CAGAAGAAATATGATAAAGAATAATAGggttcccgagacc<br>gtaatgc | TYSEDGEKRDEEKK<br>KESLGVDLGLGLKE<br>ERKYYVYVGAYVEF<br>GESDDKDKELYVGS<br>YLKLGLDDEWEAKL<br>GLYAGGKYKDKDKD<br>FEIYVGVYGGAGAD<br>KTKKLYYEFGVYTG<br>LGFDSKKDKEKGYF<br>GTGLGGQYKDNQY<br>KYRGDIEAGIGYDD<br>SKKDSAYVKIYAGW<br>DDRNKFEAGLGSGF<br>AQKKYDKE |
| TMB_n12_S14_nres14_1 | atactacggtctcaaggaAGCTATGAGAAAGAACTGGGCG<br>TCGAACTGGGCGCGGGCCTGCGCGATAACAAATT<br>CTATGTGTATATTGGCGCGTATAGCGAAGATGGCGA<br>TAAAGATGATGAAGAGAAGAAGAAGAAAAGCTACG<br>GTTTTGACGTTGGCGGCGGCGCGAAAGAAGAACG<br>CAAATTTTATTTCTATCTGGGTGTGTACGCGGAATAT<br>GGCGAAAGCGATGACAAAGATGAAGAACTGTATAC<br>CGGCCTGTATAGCAAAGCGGGCTTTGATGACGAAT<br>ATGAACTGAAAATCGGCCTGTACATCGGTCTGAAA<br>TATAAAGACAAGGACAAAGACTTCGAATTCTACGCA<br>GGCCTGTATGCGGGTACCGGTTATGATAAAACCAA<br>GAAATGGTATCTGGAAGTGGGCGTTTACGCGGGTC<br>TGGGCTATGATCGTAAGAAAGATAAAGAAAAGGGC<br>TATATCGGCGTGGGCGGTGGCTATCAGGGCAAAG<br>ATAACCAGTATAAATATCGTCTGGATGCGGAAAGCG<br>GCGCGGGTTGGGATGATAGCAAGAAAGAAAGCGC<br>GTACGTTAAAATTTATCTGGGCCTGGATGATCGCAA<br>CAAATGGGAAGGCGGCTTAGGTGTGGGCACCGCG<br>AAGAAAGATTATAAGAAAGAATAATAGggttcccgagacc<br>gtaatgc | MGSSYEKELGVELG<br>AGLRDNKFYVYIGA<br>YSEDGDKDDEEKKK<br>KSYGFDVGGGAKE<br>ERKFYFYLGVYAEY<br>GESDDKDEELYTGL<br>YSKAGFDDEYELKI<br>GLYIGLKYKDKDKDF<br>EFYAGLYAGTGYDK<br>TKKWYLEVGVYAGL<br>GYDRKKDKEKGYIG<br>VGGGYQGKDNQYK<br>YRLDAESGAGWDD<br>SKKESAYVKIYLGLD<br>DRNKWEGGLGVGT<br>AKKDYKKE |
| TMB_n12_S14_nres14_3 | atactacggtctcaaggaCGTGAGAAAGAATTTGGCGCGG<br>GCCTGAAACTGAAAAGCAAAGAAGATAACAAATAT<br>GATGCGGGCAGCGAAGTGTATTTTGGCCATCGCG<br>ATGAGAAAGATAAACGCTATCTGTATGGCGGCGTG<br>GAATTTGAATATAGCGATGACAAGAAAGATTATGAA<br>ATTAAAGCGTATGTGGGTTATGGCTATAAAGAAAAC<br>AAACAGGAATATGGCCTGGATTTCTATTTTGGTTTT<br>GGTCTGCGTGATGATAAATTTACCTATGATGTGGGC<br>TTTGGCATTCAGAGCGGCTCCGAAGAAGATAAATA | MGSREKEFGAGLKL<br>KSKEDNKYDAGSEV<br>YFGHRDEKDKRYLY<br>GGVEFEYSDDKKDY<br>EIKAYVGYGYKENK<br>QEYGLDFYFGFGLR<br>DDKFTYDVGFGIQS<br>GSEEDKYKYGWGA<br>KLGSKYYDKKLSVY |

| | | |
|---|---|---|
| | TAAATATGGTTGGGGCGCGAAACTGGGTAGCAAATATTACGATAAGAAACTGAGCGTGTATGTTGGCCTGTATGCGGAACTGGGCTATGAAGAAAATAAATACAAATATCGCTATGGCACCGGTACCGGCCTGGAAGCGAAAGATGATTGGTTTGATCTGTATATTGAAGTGGGCCTGGGCCTGGGTTATCAGGATGATAAGAACCAGGAATGGTACGCGGGCGGCAAACTGCGTGCGGGCGCGGAAGATAATAAATATAAAGGCTATACCGAAGTGAACATTGGCGGTGGCCATAAAGATAAAGAAGATTATTAATAGggttcccgagaccgtaatgc | VGLYAELGYEENKYKYRYGTGTGLEAKDDWFDLYIEVGLGLGYQDDKNQEWYAGGKLRAGAEDNKYKGYTEVNIGGGHKDKEDY |
| TMB_n12_S14_nres14_4 | atactacggtctcaaggaCGCAAACAGGATAAACTGGGCACCCAGGTGGGCTTCGGCGCGCGCGATAAGAAATGGGAAGTGTATCTGGGCTTTTGGCTGGAACAGGGTAAAGAGAAGGATGACAAGAAAGATCGTGATAAATTTGGCATTGAACTGGGTGCAGGCGCGGATAGCAAGAATAAATATCGTATTGAATTGGGCCTGTGGATTGAATATGGCAAAGAAGATGATAGCGAGAAAGAACTGTATGCGGGCACCTATCTGAAAAGCGGTTGGAAAGATGAATACGAAGTGGAATTCGGCGTTTACGCGGGCGTGCGCTACCGCGATGATAAAGATGACTATAAAGTTGAAGCGGGTCTGTATAGCGGCGGTGGCTTCGATAAAGATAACAAATGGTATTTCAAACTGGGTAGCTATACCGGCGCGGGCTTTGAAAAGAAGAAAGACAAAGAAGAAGTGTACGCCGGTGGCGGCTTTGGCGGCGAACTGGAAAAGGACGAATATAAACTGAAAATTTATTTTAAAGTGGGTGCGGGCTACAAAGACGATAAGAAGGAACGCCTGTATATTGAAGCCTATGTGGGCTATGATTCTAAGAACAAATATGAATTAGGCGCCGGCCTGGGTTTAGAAAAGGCGGATGATTATAAAGAATAATAGggttcccgagaccgtaatgc | MGSRKQDKLGTQVGFGARDKKWEVYLGFWLEQGKEKDDKKDRDKFGIELGAGADSKNKYRIELGLWIEYGKEDDSEKELYAGTYLKSGWKDEYEVEFGVYAGVRYRDDKDDYKVEAGLYSGGGFDKDNKWYFKLGSYTGAGFEKKKDKEEVYAGGGFGGELEKDEYKLKIYFKVGAGYKDDKKERLYIEAYVGYDSKNKYELGAGLGLEKADDYKE |
| TMB_n12_S16_nres14_1 | atactacggtctcaaggaGATAAAGGCTGGCGCGCCGAATTCGAATTTGAATGGTATCGCGATAACCGCTATAAAGCGGGCGTTGCGCTGGGCGTGAAACTGGGCAAGAAAGACGAAGATAAGAAAGGCTATTACGCTGAAGCGGGTGTGAAATTAGGCTTCAAAGATAACAAACTGTACAGCGGCCTGTATGCGCAGGTGGGCTACGAAAAGAGCGAAGACGAGAAACTGAAATATCGTCTGTATAGCTACTTTGGCTTCGGCTGGGATGATGATAAAGGTTATATTTATGCGGGCACCTATCTGGGCCTGCAGAACGAAGACAAGAAGGATAAGAAATATTATTACGGCGATACCGGCCTGGATGTGAAGCTGGATAAAGACAACAAATATGATGGCGGCGTTAAAACCGGCATTGGCGTGGGCTTTAAATATCATATTGACGACAAAGACAGCGAATACGAACTGGGTTTTCGCCTGGGTCTGGATAAAGATGGCGAATTTAAAAGCGGCATTCGCTTTGGCGCGCGCTATGAAGATAAAAGCGAGTATAAATATGTGTATCTGGACGCCCAGGTGGATTATTTTGAAGATCGCAAAGC | MGSDKGWRAEFEFEWYRDNRYKAGVALGVKLGKKDEDKKGYYAEAGVKLGFKDNKLYSGLYAQVGYEKSEDEKLKYRLYSYFGFGWDDDKGYIYAGTYLGLQNEDKKDKKYYYGDTGLDVKLDKDNKYDGGVKTGIGVGFKYHIDDKDSEYELGFRLGLDKDGEFKSGIRFGARYEDKSEYKYVYLDAQVDYFEDRKAEAIRIKLGAGYDEKKKD |

| | | |
|---|---|---|
| | GGAAGCGATTCGCATTAAACTGGGCGCGGGCTATGATGAAAAGAAGAAAGATTAATAGggttcccgagaccgtaatgc | |
| TMB_n12_S16_nres14_2 | atactacggtctcaaggaGATGAAGGCTGGGAAGTGGCGGTGGATGTGGAATATAAAGATGATCAGAAACTGAAAGCCGGCTTTCGCATTGGCAGCGAATTTGGCAAAGAGGATGAAAAGAAGAAAGGCTACTATTTTAAAGTCGGCCTGAAAAGCGGCCTGGAAGATAACAAATGGTATGCGGGCGTGCAGAGCACCGGCGGCTATAAAAGCGCGGAAGACGAAAAGAAAGAATTTGAAATTCAGTTTTATCTGGGCGGCGGTGCGAAAGATAAAAGCTATTATTTCTACCTGGGCATTGAAGGCGGCTGGCGCGAAAAGGATGATAAAGATAAGAAGAAATACGAAGCGAAAGTGGGCTTAGAAGCCAAATTTGATAGCAGCAACAAATATCGCGGTGGCTTTTATGCAGGCCTGGGTCTGGGCCTGCAGAAACGCGATAGCGAACGTGATAGCGATTATGAAACCGGCATTAAAGCGGGTCTGAAATCCGATGGCAAACTGGAAGCCGGTATTTATACCGGCTTAAAATATGAGAAGAAGAAAGAATATGAATATCTGTATGCGAAAACCGAAGTGAAATATGCGGATAAACGCAAAGGCGAAAGCGTGCGCGTGGAACTGGGTTTAGGCTACAAGAAGAAGAAGAAAGATTAATAGggttcccgagaccgtaatgc | MGSDEGWEVAVDVEYKDDQKLKAGFRIGSEFGKEDEKKKGYYFKVGLKSGLEDNKWYAGVQSTGGYKSAEDEKKEFEIQFYLGGGAKDKSYYFYLGIEGGWREKDDKDKKKYEAKVGLEAKFDSSNKYRGGFYAGLGLGLQKRDSERDSDYETGIKAGLKSDGKLEAGIYTGLKYEKKKEYEYLYAKTEVKYADKRKGESVRVELGLGYKKKKD |
| TMB_n12_S16_nres14_3 | atactacggtctcaaggaAAGAAATATGGCTACAAATTTGGCGTTTATACCCGCTGGTACGATGATAACGAATATGAACTGGGTAGCGGCTTTCATCTGGTGTATAAAGATGAAAGCGACAAAGGCTATAAACTGGGCGTGGGTGCGAAATACGATAAAGATAAAGACAAACTCGAAGCGGGCGGCGATTTTCAGTGGTACAAAGAAAAGAAAGACGACGAAGAGAAATGGCTGTACGTGTATGCGAAACTGGATGAAAAGGATAAGAAAGATCTGGAAATTGGTACCGGCGCGGGCTTTGGCAAAGAAGACAAGAACAACGAAAAGTATTACGTCGGCGTCTATGGCGGCCTGGGTTATGAAGATGATAAAGGTTATCTGTATGCCGGCCTGTATGTGGGCGCGCGCTGGGATGCGGATCGCAAATATAAATTTGGCACCGGCGTGTATAGCGGCGCGTACCTGTACGAAGATAAGAAATACAGCGCGGAAGTGGGTCTGTATATTAAAGTGGGCTGGTATGACAAAGATAAATTCGAAATCGGCAGCAAATTCGGTGCGGGTCTGGATCGTAAGAACCGCTTTTATATTGAATTTGGCCTGGAACTGGGCTATGGTGACAACGATCTGCTGCGCAAAAGCTAATAGggttcccgagaccgtaatgc | MGSKKYGYKFGVYTRWYDDNEYELGSGFHLVYKDESDKGYKLGVGAKYDKDKDKLEAGGDFQWYKEKKDDEEKWLYVYAKLDEKDKKDLEIGTGAGFGKEDKNNEKYYVGVYGGLGYEDDKGYLYAGLYVGARWDADRKYKFGTGVYSGAYLYEDKKYSAEVGLYIKVGWYDKDKFEIGSKFGAGLDRKNRFYIEFGLELGYGDNDLLRKS |
| TMB_n14_S16_nres12_1 | atactacggtctcaaggaAAAGAAGGCGAAATTTATAGCGGCGCGGATCTGGATAAAGATAAAAAATATGAAGAACTGCGCTTTGGCACCCGCTATGATGAAAGCCGCAGCAAAGATAAAGATGATGATTATCTGGGCGTGGATTTTGAGAGTCGCGCGAAAAATAAAGCGTATCGCGAAG | MGSKEGEIYSGADLDKDKKYEELRFGTRYDESRSKDKDDDYLGVDFESRAKNKAYREGGLGLKWGSKSK |

| | | |
|---|---|---|
| | GCGGCCTGGGCCTGAAATGGGGCTCGAAAAGCAAAGATGATAAGGACGAATTTGGCGTGCGCCTGGAAGCGGGCTGGGATGAAAAACGCAAATATTATGTGGGCGCCAAAGTGTATGCGGGCTATCGCTACGAGGATAATTTTAAAAGCGATTTCGAAGGGGAATTACGCTTGGAAGACGATTATAAACTGAAAGTGGAGGCCAAAATTGGTGTGAAATTGAAAGACGACAAAGGTGAATACGAAATTGGCGCGGGCTTCGGCGGCTACCGTGATGACGACAAAAAATTAAAATGGAAACTGGATATTTATGCAGGCATTAAATTTGACAAGGATAAATTTAGCAGCGGGGCGTGGGCGAGCGTGGAATATGATAGCAGCAATAAGAGTGATAAACTGAGCGCCGGCACCGAACTGGGGCTGCGCGATAAGGACAAAGAGCTGTATTTAGGCACCCAGCTGGACAAGGATCGCGATGACAAGGAAGAAAAAGTGCGCGTGTATGGCCGCGTTAAGAAACGCAAAAAAGATGATAAGGATGACAAATAGTAAggttcccgagaccgtaatgc | DDKDEFGVRLEAGWDEKRKYYVGAKVYAGYRYEDNFKSDFEGELRLEDDYKLKVEAKIGVKLKDDKGEYEIGAGFGGYRDDDKKLKWKLDIYAGIKFDKDKFSSGAWASVEYDSSNKSDKLSAGTELGLRDKDKELYLGTQLDKDRDDKEEKVRVYGRVKKRKKDDKDDK |
| TMB_n14_S16_nres12_2 | atactacggtctcaaggaAAAGAAGGCGAATTTTATGCGGGCATTGATTATGATAAAGATAAAAAACTGGAAGAATTTCGCAGCGGCCTGCGCTATGATGAAAGCAAAGACAAGGACAAAGATGATGATTATTGGGGCGTGGATGTGGAAGTGCGCGCGAAAAATAAATTCTATCGCGAAATTGGCTCGGGCGTGAAATTTGGCAAAAAAAGTAAGGATGATAAGGAGGAATATGGCGCGCGCGTGGAACTGGGCTATGACGAAAAACGCAAATGGTATGCAGGCGTCAAAAGCTATCTGGGGTATCGCTACGAAGATAATTTTAAAATTGATCTGGAGGTGGAAGCGCGTTACGAGGACGATCTGAAAATTAAAACCGAAGTGAAAACCGGCCTGAAATATAAAGACGACAAAGGTGAACTGGAAGGCGGCTTTGGCCTGGGCGTGTATCGTGATGACGATAAGAAAGCCAAATGGAAGTTGGACGTGTACGCCGGCGGCAAATATGACAAAGATAAACTGAGCTTAGGCACCTGGGCGAGCGCGGAATTTGATAGCAGCAATAAGCTGGATAAGTTTAGCTCTGGCGCCGAGCTGGGCTTACGCGATAAGGATAAGGAAGCGTACGCGGGTCTGCAGATTGACAAAGATCGCGATGACAAGGAAGAAAAGCTGCGCCTGTATTTTCGCTTTAAGAAACGTAAAAAAGACGATAAGGATGACAAATAGTAAggttcccgagaccgtaatgc | MGSKEGEFYAGIDYDKDKKLEEFRSGLRYDESKDKDKDDDYWGVDVEVRAKNKFYREIGSGVKFGKKSKDDKEEYGARVELGYDEKRKWYAGVKSYLGYRYEDNFKIDLEVEARYEDDLKIKTEVKTGLKYKDDKGELEGGFGLGVYRDDDKKAKWKLDVYAGGKYDKDKLSLGTWASAEFDSSNKLDKFSSGAELGLRDKDKEAYAGLQIDKDRDDKEEKLRLYFRFKKRKKDDKDDK |
| TMB_n14_S16_nres12_3 | atactacggtctcaaggaGAACGCTTTGAAGTGTGGGTGGGCCTGAGCCAGAGCAGCGATGATAAATATCGCGATGGCAAACTGGGCACCCGCTTGAGCGAAGAAGAGGATAAAAAAGATCAGAAACGCCAGGGCGGCATTGAGGTGAAAGCGAAATATAAAGATGATTATTATGAAGAAGCGGGCCTGGGTCTGTATTGGGGCGAAGCGGACAAAAGCAAAAAGAAATGGCTGGGGCTGAAAGTGGATGCGGGCTGGGATAAAGAAAATAAAGCGTATTTTG | MGSERFEVWVGLSQSSDDKYRDGKLGTRLSEEEDKKDQKRQGGIEVKAKYKDDYYEEAGLGLYWGEADKSKKKWLGLKVDAGWDKENKAYFGVDAEFGFQDDDYLRTE |

| | | |
|---|---|---|
| | GCGTGGACGCGGAATTTGGCTTTCAGGATGATGAT<br>TATCTGCGCACCGAATTTAAAATTAGCTTGAAATTC<br>AAGGATGATGCGTATACCCAGGCGGAAGTGCGCG<br>TGGGCATCGAATATAAGGACAAAGATGGCGATCTG<br>GAAGCAGGCAGCGGGATTGGTGTGCGCAAAGATA<br>AGGACCAGAAATATGAATGGTATAGCTATCTGTATG<br>CCGGCGGCCGCTATAAAAAGGATCGCTTTCGCAGT<br>GGCATTTATGCGCAGCTGGATTATGATAGCAGCTAT<br>AAGAGCGAAAAACTGCGCTTTGGCCTGGAGTTTG<br>GTTTGGATAATGATAAATATGAACTGGATGCTGGCG<br>CGGAAAGCCAGTATCAGCGCGATCGCAAGAAGGA<br>AGAAGTGGAAGTGTATCTGAAATTTCGCGAGGAAA<br>AAGACAAACAGGACGACGATAAATAGTAAggttcccga<br>gaccgtaatgc | FKISLKFKDDAYTQA<br>EVRVGIEYKDKDGD<br>LEAGSGIGVRKDKD<br>QKYEWYSYLYAGG<br>RYKKDRFRSGIYAQ<br>LDYDSSYKSEKLRF<br>GLEFGLDNDKYELD<br>AGAESQYQRDRKK<br>EEVEVYLKFREEKD<br>KQDDDK |
| TMB_n14_S16_nres12_4 | atactacggtctcaaggaGATGATCTGCGCCTGTGGGCGG<br>GCACCAATAAAGATAAGGATGATAAATATGAAGAAA<br>GCTATCTGGGCTTTGAATTTCGCAAAAGCGATGAT<br>GATAGCGATAAATATAAAAAATGGGGCGTGTATGGC<br>TACCTGAAATACAAGGATGATTGGTTTAAAGAAGCG<br>GGCGTGGGTGTGGAATTTGGCGATGAAAGCAAAG<br>ACAAATCGTATTGGCTGGGCAGCCGCATTGAACTG<br>GGCCTGGATGAGGAACGCAAATATTATTTAGGCGC<br>GAAAATTTATGGCGGCTATCGCAAGGACGACAATTA<br>TGAGAGCGAAGTGGAAGTTGAAGTGAAAGGCTATT<br>CGGATTATTACTTTGCGGCGGAGGTGAAATTGGGT<br>GCGAAACTGAAAGACGATAAAGGCGATCTGCAGTT<br>TGGCTTTGGCGCGGGGACCGAAGATGACAAGGAC<br>AAAAAATGGCGCTTTTATATTAAATTTGAAAGCGGC<br>CTGAAGGCGCGCCGCGATAAGTACGACTTGGGGC<br>TGTATGCGGATTTTAAAGCGGATAGCAAAAATGAAA<br>AAAAAGATGCGAAAATTGGCCTGAGCACCGGCTAT<br>AAACGCGATAAAAAGAAGGCCGAGGTGGGCGCGG<br>AGGTGGATCAGAGCAGCGACGACAAAGATGAGAA<br>ACTGCGCGTTGAATTGAAGATTGAAGATAAGGAAA<br>AGAAAGATAAAGACGATAAATAGTAAggttcccgagaccg<br>taatgc | MGSDDLRLWAGTN<br>KDKDDKYEESYLGF<br>EFRKSDDDSDKYKK<br>WGVYGYLKYKDDW<br>FKEAGVGVEFGDES<br>KDKSYWLGSRIELG<br>LDEERKYYLGAKIY<br>GGYRKDDNYESEV<br>EVEVKGYSDYYFAA<br>EVKLGAKLKDDKGD<br>LQFGFGAGTEDDKD<br>KKWRFYIKFESGLK<br>ARRDKYDLGLYADF<br>KADSKNEKKDAKIG<br>LSTGYKRDKKKAEV<br>GAEVDQSSDDKDE<br>KLRVELKIEDKEKKD<br>KDDK |
| TMB_n14_S16_nres12_5 | atactacggtctcaaggaAAAGATCTGCAGGTGTATGTGG<br>GCGGCGATTATGATGAAGATGAAAAATTTAAAAGCT<br>TTTATGCGGGCGCGAAATGGGAAGAAGAGGAGGA<br>TAAAAGCAAAGACAAAGAACGCTTTGGCGCAAAAG<br>TGGATGTGGAACTGAAAGATGATCTGTATCGCAGC<br>CTGGGCGCGGGCTTTGAAGGCGGCCGCAAAAAG<br>AAGGACGACGAATATGAACTGGGCGTGAAAATTGA<br>TGCCGGCTGGGATGAAAAACGCGACTATTATTTTG<br>GCGTGGAAACCGAATTAGGCCTGCAGGATAAGGAT<br>TACTTTTATATTAAATTTAAACTGCGCGGCGAATATA<br>AAGACAAATATTATCTGAAATTTGATGCGGAAGTGG | MGSKDLQVYVGGD<br>YDEDEKFKSFYAGA<br>KWEEEEDKSKDKE<br>RFGAKVDVELKDDL<br>YRSLGAGFEGGRK<br>KKDDEYELGVKIDA<br>GWDEKRDYYFGVE<br>TELGLQDKDYFYIKF<br>KLRGEYKDKYYLKF<br>DAEVGSKYKDKDG<br>RFEAGSGLGVRKDK |

| | | |
|---|---|---|
| | GCAGCAAATATAAGGATAAGGATGGCCGCTTTGAA GCGGGTAGCGGCCTGGGGGTGCGCAAAGACAAA GATGATGATGCGAAATACGAGATTTATCTGCGCTTG GGCACCGAAGCGAAAAAAGATCGCTATGATACCGG CCTGTATATTCAGAGCGATTATGATAGCAGCAATAA AAAAAAACGTCTGCGCGCCGGCGCGTATATTGGCT GGGATTACGATAAAGACCGTAGCCGCGCAGGCCT GGATGGCGAGTACGAAAGCGATAATAAGAAAAAGA AATTAAAAGTGTGGCTGGAGGTGGAAAAGAAAAAA AAAGATGATAACAAAAAAGATTAGTAAggttcccgagacc gtaatgc | DDDAKYEIYLRLGTE AKKDRYDTGLYIQS DYDSSNKKKRLRAG AYIGWDYDKDRSRA GLDGEYESDNKKKK LKVWLEVEKKKKDD NKKD |
| TMB_n16_S20_nres12_2 | atactacggtctcaaggaAGCAAATTTAAGCTGGATACCGA TCTGGAAATTAAATTCGATGATAGCGCGAAATGGAG CGCGGATGCGAAAGTGAGCGCGAAGAACAGCGAC GATCTGGATGCGAGCCTGAGCCCGCCGAAATATAG CGATGATAAATACGTGCTGAAGCTGAAAATTACCGT TCAGTGGCCGACCAGCGGCGCCAAAGGCGATGTG GATAGCAGCTATGATTCGGATGATAACAAAGCGAG CTATAGCATTGCGATTACCCTGCCGCTGTGGGATA GCAACGATGTTAAAGGCACCAGCAGCGGCAGCTAT TCTGATAGCAAATACGATTATGAAGTTACCGCGACC GGCAAAGATCCGAGCGGCGTGGATATTGATTTTAC CGGCCGCTTTAGTGATGATGACGATTTTGATTATAC CCTGAGCACCAAAGTGCAGTATCCGAAAGATAACC TGACCCCGACCTTTGATGCGACCCTGCAGGATGG CAGCCTGAAAGCGACCTTTCTGCTGAAACTGGATA CCCCGGATTGGGGTGGTCCGAAACTGAGCGTGAG CGTGCAGAGCGATCTGGACAGCAACGATAACTATA AACTGACCCTGACCGCGGATGTGAAATTTCCGGG CGCGCTGGTTAGCGTGTCCACCAAATATGATAGCA ACACCGATAAACTGAGCCTGGATGCCAAACTGACC GCGAGCAGCCTGGATTAATAGggttcccgagaccgtaatgc | MGSSKFKLDTDLEIK FDDSAKWSADAKV SAKNSDDLDASLSP PKYSDDKYVLKLKIT VQWPTSGAKGDVD SSYDSDDNKASYSI AITLPLWDSNDVKG TSSGSYSDSKYDYE VTATGKDPSGVDID FTGRFSDDDDFDYT LSTKVQYPKDNLTP TFDATLQDGSLKAT FLLKLDTPDWGGPK LSVSVQSDLDSNDN YKLTLTADVKFPGAL VSVSTKYDSNTDKL SLDAKLTASSLD |
| TMB_n16_S20_nres12_3 | atactacggtctcaaggaAACGAAAAGATTAAAGTCGATAG CGGCCTGAAACTGGATGATGGCGATGCGAAATGG TGGACCAAAGTGAAACTGAAAACCAAGAAAGATGA CGACTACGACGTGGAATCCAAAATTGATTTTGACG ATGGCAAATATCATCCGGATATTAGCGTGACCGTGT ATGATCCGACCGATCCGAACGATTATCTGACCGGC ACCGTGAGCTTAGATGATGATGATAACGATCCGCC GAAAGTGACCTTTAGCGCGAAAGCGAAATACGGC CCGTTTGATCTGGAATTTAGCGTCACCCTGTTTCT GGATACCAACAAGAAGAAGAAAGTTCTGCTGAACG TGAAAGGCGATGATAGCGACGGCAAAAGCGCGCT GCTGAAAGCGGATCTGGATGACGATACCAATACCT ATACCGCCGATCTGACCTTTTCGGGCAGCTTTCCG GGCTATAAATTTACCCTGACCAGCCAGTGGAACTAT GATAGCAGCTATAAAGCGAACCTGAACAGCCTGGC | MGSNEKIKVDSGLK LDDGDAKWWTKVK LKTKKDDDYDVESKI DFDDGKYHPDISVT VYDPTDPNDYLTGT VSLDDDDNDPPKVT FSAKAKYGPFDLEF SVTLFLDTNKKKKVL LNVKGDDSDGKSAL LKADLDDDTNTYTA DLTFSGSFPGYKFT LTSQWNYDSSYKAN LNSLAQLDFPGIKGT TKATWDSDTDDAKP DPPSLSLDYPDASA |

| | | |
|---|---|---|
| | GCAGCTGGATTTTCCGGGCATCAAAGGCACCACC<br>AAAGCGACCTGGGATAGCGATACCGATGATGCCAA<br>ACCGGACCCGCCGAGCCTGTCTTTAGATTATCCGG<br>ATGCGAGCGCGTTTATTAAAGTGAGCCTGGATAAA<br>GATAGCGATAGCTACAAATGGGAAGCGCCGGATCT<br>GAGCCTGGAAGTGGATTAATAGggttcccgagaccgtaatg<br>c | FIKVSLDKDSDSYK<br>WEAPDLSLEVD |
| TMB_n16_S20_nres12_1 | atactacggtctcaaggaAGCAGCGATCCGAAACTGAAAG<br>CGGGCCCGAAACTGAGCGACGGCAAATGGCGCTT<br>TTATACCAGCGCGACCGTGGATCTGAAAAGCGATG<br>ATAGCTATAAAAGCTCGGTGACCCTGGATCTGCGC<br>GATGGCAAATACGATATTAAACTGACCCTGAGCGT<br>GCTGGATCCGACTGATCCGACCGATAGCGCGGAA<br>CTGAGCGGTCAGTATGATCTGGATGATAACGATCC<br>GCCGAAAAGCACCATTAGCGCGAAATTTAAATTTG<br>GCCCGTTTGATGGCCAGCTGAGCCTGACCTTTTG<br>GCTGGATACCAACAGCCGCAAAAGCAGCGAACTG<br>AGCGTGAAAGTGGATGATTCGGATGGCCGTCGTG<br>CGCAGCTGAGCGCGCTGTTAGATGATAGCACCAAC<br>TCTTATACCTTTGTGGGCACCATCGATACCAAACTG<br>CCGGGCTGGGATGGCAGCACCACCATTAAAGCGG<br>ATTGGGATAGCAGCGCCAAATATCAGCTGAGCGTG<br>ACCGTGACCAGCGAATTTCCGGGCCTGAAAGCGA<br>GCGCAAGCGTGAGCTATGACAGCAACACCGACAG<br>CTATCAGCTGACCGGCCCGGATATCAAAGTTGATTA<br>TCCGAACGGCGATGCGAGCGCGTCGCTGACCTAT<br>GATAACACCAACGATAAATATGATGCGAAACCGCC<br>GGATGTGCGCGTTAAAGTGGATTAATAGggttcccgag<br>accgtaatgc | MGSSSDPKLKAGPK<br>LSDGKWRFYTSATV<br>DLKSDDSYKSSVTL<br>DLRDGKYDIKLTLSV<br>LDPTDPTDSAELSG<br>QYDLDDNDPPKSTI<br>SAKFKFGPFDGQLS<br>LTFWLDTNSRKSSE<br>LSVKVDDSDGRRA<br>QLSALLDDSTNSYT<br>FVGTIDTKLPGWDG<br>STTIKADWDSSAKY<br>QLSVTVTSEFPGLK<br>ASASVSYDSNTDSY<br>QLTGPDIKVDYPNG<br>DASASLTYDNTNDK<br>YDAKPPDVRVKVD |
| TMB_n16_S20_nres12_4 | atactacggtctcaaggaAGCCTGCCGGAAAACGCGGGC<br>ATTAGCTATAAATGGGATCCGGATGATTTTAGCTTTG<br>AAAGCAAACTGACCCTGAAAGACCCGGATAAAGAT<br>AGCCGCGAAGCGAGCCTGGATCTGAGCGTGAAAG<br>CCGATGATAACTTCAAAAGCGATTACGACCTGAAA<br>GTGAAACTGTCGCTGCCGGGCGCGGATGTGGATG<br>TGCTGGTGACCTATAACGATAGCAGCGATCTGTCG<br>TATAAATATGATGTTACCGTGCAGTTTCGCCTGCCG<br>ACCTTTGATGTGACCGCGAGCAGCAGCCTGGATAA<br>TGATGATCCGACCAGCAACAAATATAGCAGCACCG<br>TTAAACTGGCGAAAGATCCGAGCGCGGATATTGAT<br>GGCACCGTGACCTGGGATCCGACCTCCGCGAAAT<br>TTACCCTGACCCTGCGCCTGAAACTGGATACCAAC<br>GATCTGGATCTGAAAGCGGATTATACCTGGAGCAG<br>CGATGATCCGCCGAAATATACCACCAGCGTTAAAG<br>CAGATTTTAAATATCCGAGCCATAAAGGCGATATCG<br>TTGCGAAATACGATCTGAAGAACGGCGCGTATAAG<br>AAAGCGGACGGCACCACCAAACCGGATTATCCGG | MGSSLPENAGISYK<br>WDPDDFSFESKLTL<br>KDPDKDSREASLDL<br>SVKADDNFKSDYDL<br>KVKLSLPGADVDVL<br>VTYNDSSDLSYKYD<br>VTVQFRLPTFDVTA<br>SSSLDNDDPTSNKY<br>SSTVKLAKDPSADID<br>GTVTWDPTSAKFTL<br>TLRLKLDTNDLDLKA<br>DYTWSSDDPPKYTT<br>SVKADFKYPSHKGD<br>IVAKYDLKNGAYKKA<br>DGTTKPDYPGTSPS<br>ISWDLDSDTDKLSF<br>DASPPDGKLD |

| | | |
|---|---|---|
| | GCACCAGCCCGAGCATTAGCTGGGATCTGGATAG<br>CGATACCGATAAACTGAGCTTCGATGCGAGCCCGC<br>CGGATGGCAAACTGGATTAATAGggttcccgagaccgtaat<br>gc | |
| TMB_n16_S16_nres14_1 | atactacggtctcaaggaGATTATAAAGCGGATGCGCGTAC<br>CCGCGCGGCGCTGGAAAAAGATAAAGACAATAAAG<br>AAGAAAAACTGGAAATTCAGGTGGATTTTGAAAGC<br>AAATGGTGGGATCGCAAAGATGAAGAATTTCAGGT<br>TGAACTGAGCAGCAAATATGAAAAACGCAAAAAGG<br>ATTACAAGGACGATGATAAATACGATGTGGCGCTGA<br>AAGCGCGCGTGAAATTGGATAAGGATCAGTCGAGC<br>AGCGAAGTGGAATCGAAATATACCGATAAGGATAG<br>CAAAAAAGACTATGATGTGAGCCTGAAACTGGCGA<br>TTAAACGCAAATTTCTGGACAAGGATGATTACGAGA<br>GCGAAGGCAGCGCGGAAGGCCAGTTTAAATTTGA<br>TAAGGATAAACGCGAAAAAGATGCGAAACTGCGTG<br>TTAAGTTAGACTTTCGCTGGAAGAAGGACAAGTAC<br>GAAAGTAAATTGCGCATTCGCGCCGAATACGAGGA<br>AAAAAAGGAGAAAAAAGATAAGCGCAAACAGTTTG<br>AAGCGGAAGCGCAGGCGGATTATGATGAATCGGAT<br>GCAAAAGTGCAGATTAAAACCAATCTGGCGTTGGA<br>TGAGGAGGAAGACAAAGATAAAAAGAAAAAGAAAG<br>AATATCGCCTGGAAAGCAGCCTGGCGGTGCAGTT<br>CAAATTTAGCCAGCTGGATGGCCGCATTCAGTTTA<br>GCGCGAGCTTTAAAACGAAAAAGGACGATCGTAAG<br>GAGGACGAGAGTGAATTCCAGGCGAGTTTTGAAG<br>TGTATTATAATAAGGATAAAGTGCGCTCGAAAGTGG<br>ATACCAAAAAAAAAGAGAAGGATGTGGATGAAGAT<br>AAGAAGAAACGTGAAAGCTGGCGCTTGGAGTTGG<br>AACTGCAGTTGGAGGATAAGAAAAAAGACGATTAG<br>TAAggttcccgagaccgtaatgc | MGSDYKADARTRA<br>ALEKDKDNKEEKLEI<br>QVDFESKWWDRKD<br>EEFQVELSSKYEKR<br>KKDYKDDDKYDVAL<br>KARVKLDKDQSSSE<br>VESKYTDKDSKKDY<br>DVSLKLAIKRKFLDK<br>DDYESEGSAEGQF<br>KFDKDKREKDAKLR<br>VKLDFRWKKDKYES<br>KLRIRAEYEEKKEKK<br>DKRKQFEAEAQADY<br>DESDAKVQIKTNLAL<br>DEEEDKDKKKKKEY<br>RLESSLAVQFKFSQ<br>LDGRIQFSASFKTKK<br>DDRKEDESEFQASF<br>EVYYNKDKVRSKVD<br>TKKKEKDVDEDKKK<br>RESWRLELELQLED<br>KKKDD |

**Table S3**. Crystallography data collection and refinement statistics.

| | **n6_S8_nres10_D2** |
|---|---|
| **Resolution range** | 38.87 - 2.03 (2.08 - 2.03) |
| **Space group** | P 32 2 1 |
| **Unit cell** | 74.19 74.19 97.635 90 90 120 |
| **Unique reflections** | 20611 (1442) |
| **Multiplicity** | 13.5 (13.0) |
| **Completeness (%)** | 99.95 (99.72) |
| **Mean I/sigma(I)** | 14.27 (1.01) |
| **Wilson B-factor** | 50.84 |
| **R-merge** | 0.1028 (2.213) |
| **R-pim** | 0.02934 (0.6344) |
| **CC1/2** | 0.998 (0.644) |
| **Reflections used in refinement** | 20611 (1442) |
| **Reflections used for R-free** | 2003 (138) |
| **R-work** | 0.2197 (0.3306) |
| **R-free** | 0.2652 (0.3845) |
| **Number of non-hydrogen atoms** | 2165 |
| **macromolecules** | 2070 |
| **ligands** | 38 |
| **solvent** | 57 |
| **Protein residues** | 270 |
| **RMS(bonds)** | 0.002 |
| **RMS(angles)** | 0.51 |
| **Ramachandran favored (%)** | 98.86 |
| **Ramachandran allowed (%)** | 1.14 |
| **Ramachandran outliers (%)** | 0.00 |
| **Rotamer outliers (%)** | 0.00 |
| **Clashscore** | 2.60 |
| **Average B-factor** | 62.16 |
| **macromolecules** | 61.96 |
| **ligands** | 75.57 |
| **solvent** | 60.29 |

Statistics for the highest-resolution shell are shown in parentheses.
